# Weber’s law is the result of exact temporal accumulation of evidence

**DOI:** 10.1101/333559

**Authors:** Jose L. Pardo-Vazquez, Juan Castiñeiras, Mafalda Valente, Tiago Costa, Alfonso Renart

## Abstract

Weber’s law states that the discriminability between two stimulus intensities depends only on their ratio. Despite its status as the cornerstone of psychophysics, the mechanisms underlying Weber’s law are still debated, as no principled way exists to choose between its many proposed alternative explanations. We studied this problem training rats to discriminate the lateralization of sounds of different overall level. We found that the rats’ discrimination accuracy in this task is level-invariant, consistent with Weber’s law. Surprisingly, the shape of the reaction time distributions is also level-invariant, implying that the only behavioral effect of changes in the overall level of the sounds is a uniform scaling of time. Furthermore, we demonstrate that Weber’s law breaks down if the stimulus duration is capped at values shorter than the typical reaction time. Together, these facts suggest that Weber’s law is associated to a process of bounded evidence accumulation. Consistent with this hypothesis, we show that, among a broad class of sequential sampling models, the only robust mechanism consistent with reaction time scale-invariance is based on perfect accumulation of evidence up to a constant bound, Poisson-like statistics, and a power-law encoding of stimulus intensity. Fits of a minimal diffusion model with these characteristics describe the rats performance and reaction time distributions with virtually no error. Various manipulations of motivation were unable to alter the rats’ psychometric function, demonstrating the stability of the just-noticeable-difference and suggesting that, at least under some conditions, the bound for evidence accumulation can set a hard limit on discrimination accuracy. Our results establish the mechanistic foundation of the process of intensity discrimination and clarify the factors that limit the precision of sensory systems.

---

Stimulus intensity is one of the fundamental dimensions of the sensory experience. Under-standing the relationship between the physical intensity of a stimulus and the subjective intensity of its associated percept was the main driving force behind the development of the field of psychophysics (1–6). This effort was propelled by the finding that the discriminability between two nearby stimuli along a sensory continuum depends only on the ratio between their intensities, not on their absolute magnitudes. This observation was first made by Weber in 1834 (1) and was soon after further characterized by Fechner, who gave it the name of Weber’s law (WL) (2). Hundreds of studies describing sensory discriminations in audition, vision, taste, olfaction, somatosensation and temperature have replicated WL (4–6). WL embodies a non-trivial computation, since sensory receptors and sensory neurons in the periphery encode absolute magnitude explictly in the form of monotonic increases in firing-rate. The way in which the absolute magnitude of the stimulus is factored out during discrimination is not fully understood, although many mechanisms to explain it have been proposed (2, 4, 5, 7–9), (see (5) for a summary of early work).

The modern study of perception has also been motivated by understanding the limits of discrimination accuracy (10–12). However, whereas early work focused on discrimination and estimation of unstructured stimuli varying along simple sensory continua (e.g., luminance, weight) and used mainly Signal Detection Theory (SDT (13)), modern approaches have focused on the concept of bounded accumulation of evidence using temporally structured input. The resulting paradigm, which rests formally on the sequential sampling (SS) framework (14–19), has been extremely successful in guiding the development of hypothesis about perception and decision-making (20–26). The use of complex stimuli, however, makes it difficult to understand what exactly is the nature of the evidence used to guide discriminative choices, complicates teasing apart external and internal sources of noise (but see (25)), and has led to an emphasis on the differential component of discrimination at the expense of a better understanding of the role of overall stimulus magnitude. As a consequence, classic problems such as WL have not received large attention in the community studying perceptual decision making. An important exception is the work of Link, whose ‘wave theory’ showed that WL arises naturally from bounded accumulation of evidence when the intensity of the stimulus is represented by the rate of a Poisson process (4). More recently, some studies have realized the importance of reaction time (RT) as a diagnostic tool for comparing different models of WL (8), and it has been observed that Link’s model makes the prediction that overall stimulus level should rescale the RT distribution (RTD) (9). However, despite these efforts, the empirical data available does not unambiguously establish whether and how WL should be understood within a SS framework. Indeed, the most in-depth treatments still explain WL within SDT (5). Here we present empirical data and theoretical work showing conclusively that WL is an unavoidable con-sequence of exact temporal bounded accumulation of sensory evidence encoded in the brain using Poisson statistics. We studied WL by training rats to report the lateralization of binaural sounds of different overall levels. Rodents use inter-aural level differences (ILDs) caused by the acoustic shadow of the head (27) to localize sound on the horizontal plane (28, 29). ILD discrimination thus embodies a comparison of sound intensities that can be used to study WL (30). Consistent with WL, a multiplicative change in the intensity of the two sounds being discriminated did not modify discrimination accuracy. In fact, we demonstrate that the sole effect of this manipulation is to change the effective unit of time of the discrimination process. We also show that this constraint specifies the computational mechanism underlying WL in a way that permits a virtually complete quantitative description of the behavior of the rats.

## Behavioral correlates of level-invariant discrimination

Since WL is well known to robustly hold for white noise discriminations (5, 30, 31), we trained rats to discriminate the lateralization of broadband noise bursts (5-20 KHz) in a standard three-port behavioral box (Fig. 1A),. We varied ILDs while keeping the average binaural level (ABL) constant (Fig. 1C). Sounds were played through headphones to minimize uncontrolled stimulus variations (Methods; Fig. S1A). Discriminations were performed in a RT configuration (Fig. 1AB; Methods), with ILDs varying pseudo-randomly across trials and ABLs varying pseudo-randomly in blocks of 80 trials (Fig. 1C). Rats were trained to perform at their psychophysical discrimination threshold (Fig. S2B).

**Figure 1:**
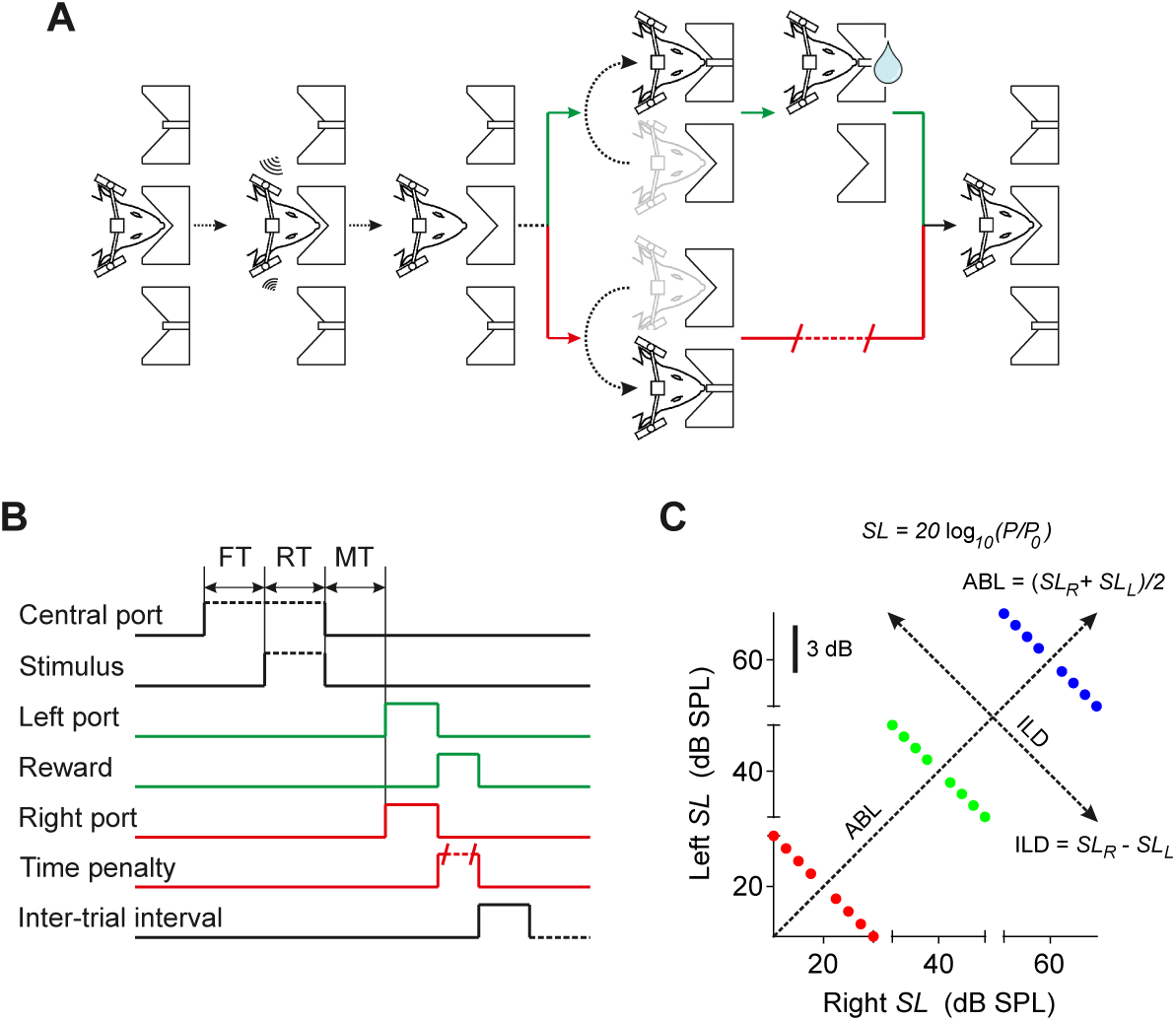
Task structure and stimulus set. (A) Schematic depiction of different task events. Rats were rewarded with water for making the correct choice and were punished with a time delay for making an error. **(B)** Time-line of relevant task events. FT, fixation time. RT, reaction time. MT, movement time. **(C)** Stimulus set. All sounds were cosine-ramped broadband (5-20 KHz) noise bursts. The ABL (ILD) of a particular stimulus is given by the average (difference – by convention right minus left) of the intensity of the sound in dB SPL (sound level *SL*) across both speakers. *P*0 = 20 *μ*Pa is the reference pressure of the SPL scale.

The accuracy of ILD discriminations did not depend on ABL (Fig. 2A-B). Differences in the *d*’ index for ILD discriminations at ABLs of 40 and 60 dB SPL did not reach statistical significance for any rat (*p >* 0.05, Fisher’s exact test, Bonferroni corrected; see Table S1). For two of the five rats, *d*’ for ABL = 20 dB SPL was significantly different than that for 40 and 60 dB SPL (in one case larger and in the other smaller; *p <* 0.05, Fisher’s exact test, Bonferroni corrected). At the group level, differences in *d*’ were again not statistically significant for any of the three ABLs (*p >* 0.05, Fisher’s exact test, Bonferroni corrected). Thus, as shown before for other species (32–34), rats also display level-invariant ILD discrimination accuracy, at least for broadband noise. Because the ILD of the stimulus is a function (logarithm) of the ratio of the root-mean-square (RMS) pressure across the two ears (ILD = 20 log(*P*_*R*_/*P*_*L*_); Fig. 1C), this result is equivalent to WL.

**Figure 2:**
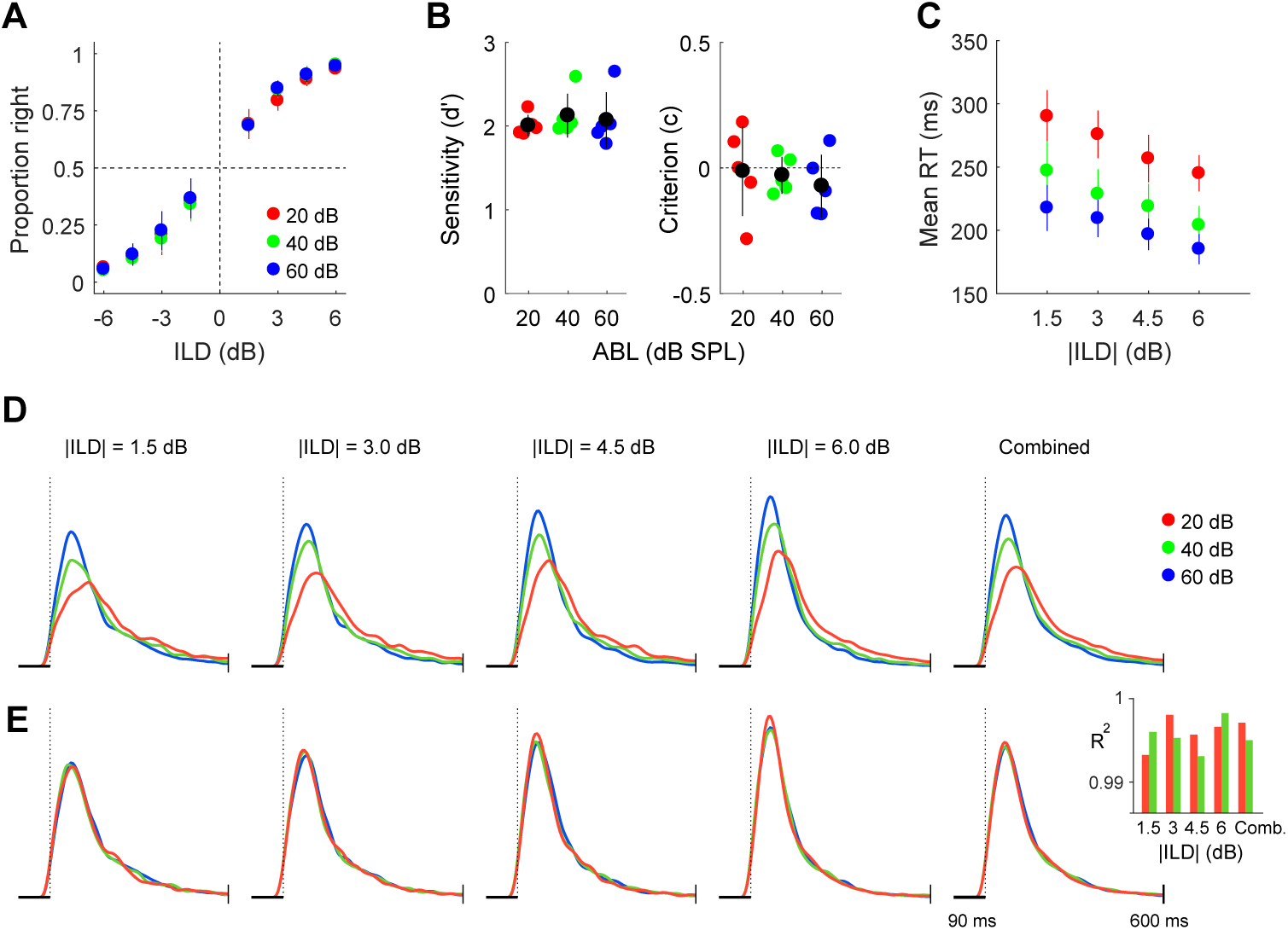
Behavioral correlates of level-invariant ILD discrimination. **(A)** Choose-right probabilities as a function of ILD for each ABL separately (mean *±* SD across rats). **(B)** Sensitivity (*d*’, left) and criterion (*c*, right) for each animal (*n* = 5; black: mean *±* SD across rats). **(C)** Reaction time (mean *±* SEM across rats; full reaction time distributions (RTDs) for individual rats are shown in Fig. S9) as a function of difficulty for each ABL separately. **(D)** RTDs for the three ABLs are shown separately for each difficulty, and combined across difficulties (right). For all RTDs, the dashed line indicates the time at which RTs become condition-dependent (90 ms, scale bar in all plots – see Methods; Fig. S3). Each RTD contains all data for that condition from all rats. **(E)** For each difficulty, we have rescaled time uniformly (see Methods; Fig. S4) to maximize the overlap of each RTD with that of the loudest sound (ABL = 60 dB SPL). **(Inset)** Accuracy of the shape-invariance of the RTDs. Each bar is the R^2^ of a linear fit of the percentiles of each RTD against those of the RTD for ABL = 60 dB SPL (see Methods; Fig. S4).

We observed that more difficult discriminations have longer RTs (Fig. 2C), a signature of the standard speed-accuracy trade-off associated to bounded evidence accumulation (14, 19, 35). How-ever, we additionally observed that discriminations involving overall quieter sounds (lower ABLs) also took longer on average (Fig. 2C; see also (8, 9, 33)). Both ILD and ABL had a significant impact on mean RT on each individual rat (Table S1) as well as at the group level (significant effect of ILD, two-way RM-ANOVA, *F*(3, 12) = 17.54, *p*=0.0001; significant effect of ABL, two-way RM-ANOVA, *F*(2, 8) = 77.12, *p <*0.0001). To further characterize the effect of stimulus intensity on RT, we examined the dependence of the full RTDs on ABL, excluding very short RTs which reflect anticipation and are condition-independent (see Methods; Fig. S3). RTDs are right-skewed, as is characteristic of integration to bound models (19, 36), and are also more narrow for louder sounds (Fig. 2D), as expected from the mean RT data (Fig. 2C). However, the shape of the distributions does not depend on ABL. To reveal this, we scaled the distributions for ABL = 20 and 40 dB SPL so as to maximize their overlap with the one for ABL = 60 dB SPL (see Fig. S4). Remarkably, the rescaled distributions are almost identical for each difficulty and for all difficulties combined (Fig. 2E). In each case, more than 99% of the variance in the shape of the RTD for one ABL could be explained by the shape of the RTD for a different ABL (Fig. 2E, inset; mean R^2^=0.996; see Methods). This result, together with the level-invariance of discrimination accuracy (Fig. 2A-B), demonstrates that the sole effect of changes in ABL is to change the effective units of time of the sensory discrimination process (for an in-depth analysis of the effect of difficulty on the RTD see Figs. S4, S6), with a shorter effective unit of time for louder sounds. This implies that, at least for the purposes of discrimination, the effective duration of a sound increases with its intensity.

## Weber’s law breaks down for controlled short sound durations

The speed-accuracy trade off (Fig. 2C) and the right-skew of the RTDs (Fig. 2D,E) suggest that rats are solving the task by a process of bounded accumulation of sensory evidence (18, 36). If this was the case, the results in Fig. 2 make the prediction that, for fixed sound durations (shorter than the time it typically takes for the rats to decide), performance should be worse for lower ABLs, since the effective duration of the integration period for lower ABLs would be shorter. Thus, WL should breakdown in an ABL-dependent manner. We tested this prediction in a series of sessions where the rats were still free to choose when to exit the central port, but if they had not exited by a maximum sound duration SD_max_, the sound stopped (see Methods). We used various values for SD_max_ within a session and considered only ABL = 20 and 40 dB SPL for this experiment, since they are the slowest conditions and would presumably be more affected by the capped sound duration. Qualitatively, performance degrades and becomes ABL-dependent for short SD_max_ (Fig. 3A). To quantify this effect, we fit a sigmoid with two parameters (asymptote and threshold; see Methods) to the difference in discriminability between the RT sessions and each SD_max_ condition, separately for each ABL (Fig. 3B). As SD_max_ decreases, performance degrades but, critically, it degrades more for the lower ABL (asympotote for 20 (40) dB SPL = 0.99 (0.59); *p <*0.0005, permutation test for a comparison of the asymptote; see Methods) for every fixed duration. Furthermore, the inflection point of the sigmoid also occurs at longer durations for ABL = 20 dB SPL (threshold for 20 (40) dB SPL = 260 (195) ms; *p*=0.003, permutation test for a comparison of the threshold). These results (Figs. 2-3) suggest that WL occurs because multiplicative changes in stimulus intensity only modify the effective unit of time of a bounded evidence accumulation process.

**Figure 3:**
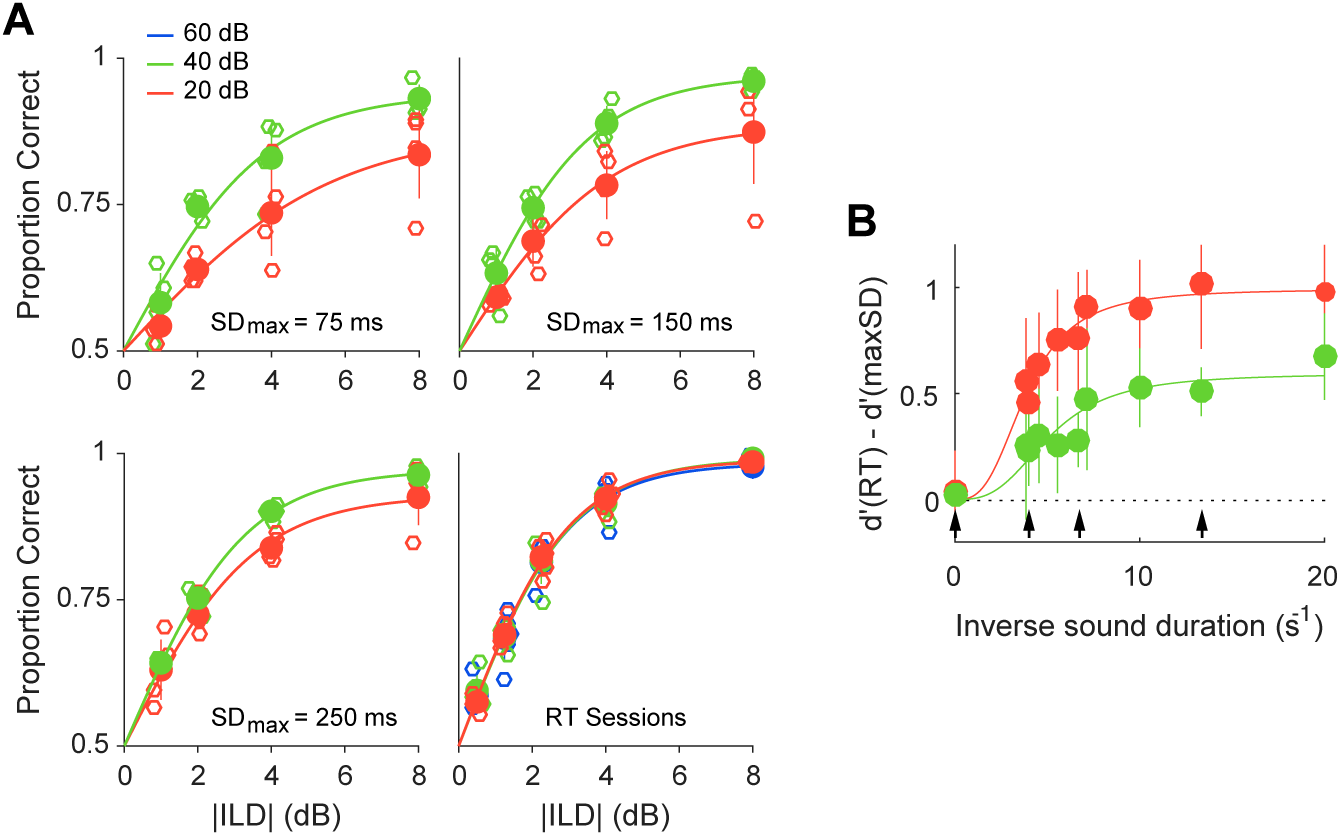
Breakdown of Weber’s law for short controlled sound durations (A) Psychometric functions for three conditions where the maximum sound duration SDmax was 75, 150, 250 ms and for the RT sessions where the sound always stopped only when the choice was made. Empty dots are individual rats (n=4) and filled dots are mean *±* SD across rats. Fits are from a two parameter logistic function (slope and asymptote; see Methods). **(B)** Difference between the discriminability index (d’) for the RT sessions and for each SDmax, separately for each of the two ABLs as a function 1/SDmax (filled circles; mean *±* SD across rats). We used 1/SDmax to avoid having to arbitrarily specify a SDmax for the RT sessions. Lines are fits to a sigmoidal function (see Methods). Arrows show the four SDmax for which psychometric functions are shown in (A).

## The mechanism underlying Weber’s law

How narrowly can the mechanism that underlies discrimination be specified solely from the scale-invariance of the RTDs with respect to ABL? Since our data (Figs. 2-3) suggests a mechanism based on bounded accumulation of evidence, our starting point was the broad class of models in which choices are triggered when the decision variable (DV) hits for the first time either of two (possibly time-dependent) bounds, and where the DV evolves in time as a continuous Markov process (CMP) (37). Most proposed models for perceptual decision-making are, or can be construed as, CMPs (4, 35, 36, 38–41). Qualitatively, the only assumptions in a CMP are that successive increments in the DV are statistically independent and of a magnitude which vanishes as the time increment goes to zero (hence, the word ‘continuous’), which we expect to be appropriate since our stimuli have constant intensity. We therefore asked: if we consider a discrimination involving a sound with RMS pressure at the two ears (*P*_*R*_, *P*_*L*_) and another discrimination involving (*kP*_*R*_, *kP*_*L*_), which members of this model class have the property that the second discrimination proceeding under a given temporal variable *t* is identical to the first discrimination proceeding under a rescaled temporal variable *t*′ = *αt*? Our analysis (Supplementary Information) reveals that this condition is very restrictive, allowing only two types of solutions. One type requires a precise relationship between the shape of the time-varying bound, the leak of the DV and the statistics of the evidence. This solution is not robust (it requires careful parameter tuning) and requires unplausible relationships between disparate processes (e.g., spiking statistics and decision bounds), so we don’t consider it further. The only alternative solution just requires the following conditions: a constant decision bound, no leak (i.e., perfect accumulation of evidence), a power-law relationship between physical stimulus intensity and sensory evidence, and a linear relationship between the variance of the evidence and its mean. Qualitatively, the essence of this solution can be traced to the fact that the spike count statistics of a Poisson process are invariant if the rate and the count window are modified by multiplicative factors *k* and 1/*k* respectively. This property extends to any process with a linear variance-to-mean relationship (i.e., constant Fano Factor) and is relevant because the DV in the model effectively ‘counts’, i.e., temporally integrates, sensory spikes. Finally, the powerlaw transformation is necessary so that multiplicative factors in stimulus intensity translate into multiplicative factors in firing rate, which can then be compensated by a rescaling of time.

#### Model parameters

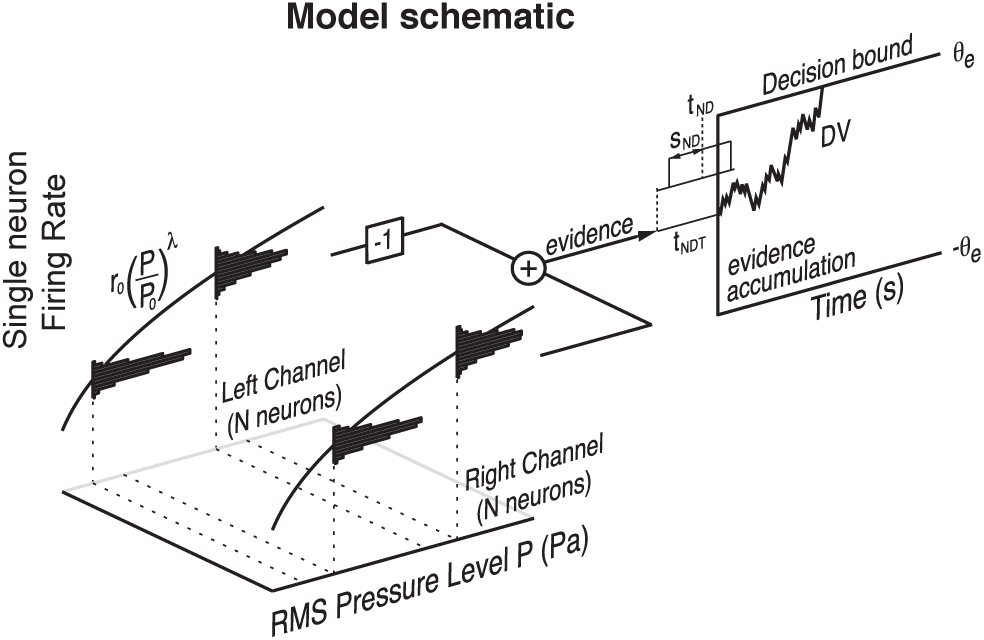

###### Discrimination process

λ amount of compression in the pressure-to-rate transformation

*Nr*_*0*_ net gain of the pressure-to-rate transformation of each channel of *N* neurons

θ_e_ decision threshold in units of the single-spike quantum of evidence

###### Non-decision time

*t* _*ND*_ across-trial mean of non-decision time

*s* _*ND*_ across-trial spread of non-decision time

#### Sensory evidence

The sound level at each ear is *SL* = *20 log*_*10*_ *(P/P*_*0*_ ) dB SPL and ABL = *(SL*_*R*_ + *SL*_*L*_ *)/2*, and ILD = *SL*_*R*_ *- SL*_*L*_. Thus, the firing rate of the neurons in each channel can be written as

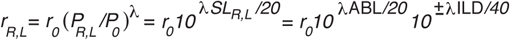

These two functions are plotted on the right (with λ = 0.3) as surfaces in the (ILD,ABL) plane. The evidence is the difference between the activity of the left and right channels.

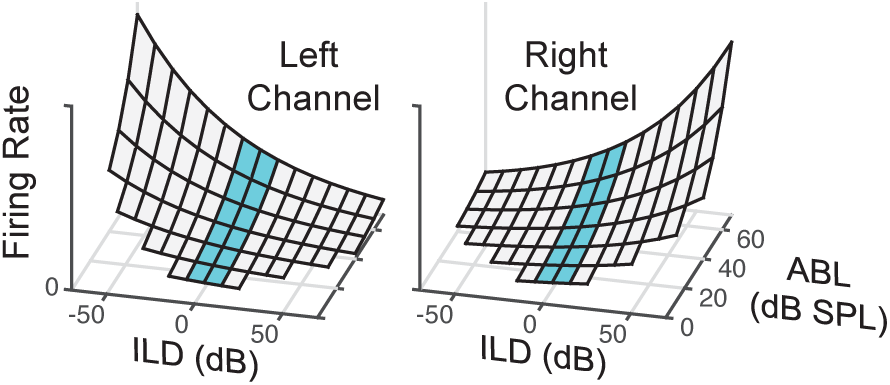

Since the firing statistics are Poisson, the mean (variance) of the evidence *e(t)* is proportional to the difference (sum) between these two surfaces. Close to psychophysical threshold (ILD close to zero; plots on the left), the mean is linear in ILD. The slope of this linear relationship is a gain factor *g(*ABL,λ) that increases with ABL (see Supplementary Information). The linear terms for the left and right channels have opposite sign an thus cancel when added, so the va riance is approximately independent of ILD, but increases with ABL through the same gain factor. Thus

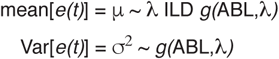

#### Discriminative choice

The instantaneous decision variable *x(t)* reflects the accumulated evidence,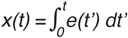. Thus, *x(t)* evolves in time according to the diffusion equation *dx/dt* = μ + σ η*(t)* where μ and σ^2^ are the just-specified mean and variance of the evidence. A right (left) choice is made when the accumulated evidence reaches a bound of θ −θ). If neurons have Poisson statistics and firing rates encode the stimulus as a power-law, variations in stimulus intensity change the mean and the variance of the evidence by the same factor *g(*ABL,λ). Thus, rescaling time as τ = *t/g(*ABL,λ), the evolution of the decision variable in the new time units close to psychophysical threshold is *dx/d*τ = *c*_*1*_ λ ILD + *c*_*2*_ η(τ), where *c*_*1*_ and *c*_*2*_ are constants independent of ABL and ILD. Therefore, discrimination accuracy and the shape of the reaction time distribution depend on difficulty only through ILD, whereas ABL only sets the effective unit of time of the discrimination process.

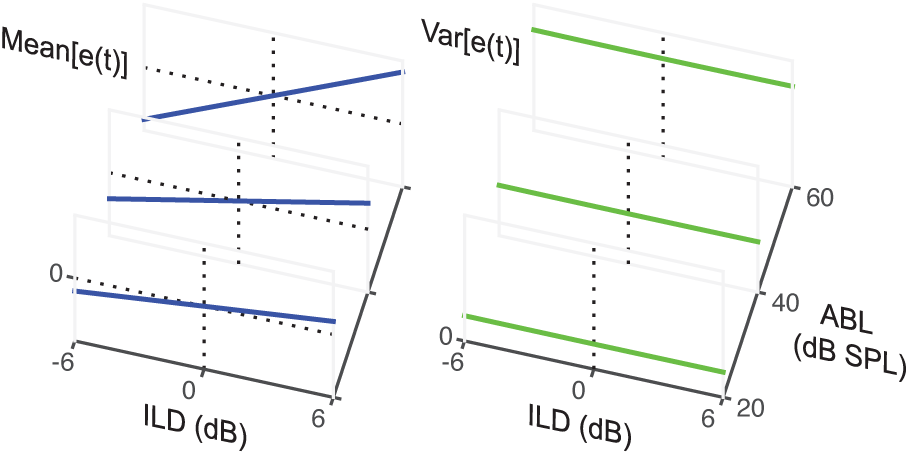

We constructed a minimal implementation of this solution (see Box, Supplementary Information) in which the evidence is the difference between the instantaneous activity of two sensory channels, corresponding to the two ears. The firing rate of the *N* neurons in each channel is a power-law of the RMS pressure level at the ear, and neurons fire with Poisson statistics. The DV integrates the evidence in time and a choice is triggered when a constant bound at DV = *±θ* is hit for the first time (decision-time). Choice and decision-time in the model depend only on three parameters (see Box): the power-law exponent (*λ*) and net gain (*Nr*_0_) of the pressure to rate transformation of each channel, and the decision bound in units of the single-spike quantum of evidence (*θ*_*e*_). This is the absolute minimum number of parameters possible in a bounded accumulation model with non-linear stimulus encoding. In the Supplementary Information we show that, close to psychophysical threshold, choice dynamics in this model are captured by a single-parameter drift-diffusion model (DDM)

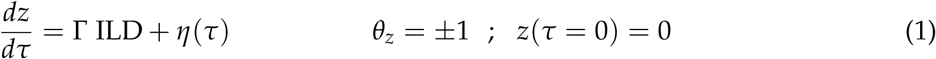

where *z* is the DV, and the single parameter Γ = *λθ*_*e*_/[40/log(10)] measures how the stimulus’ ILD sets the overall strength of the evidence of the discrimination. The stimulus’ ABL does not appear in this equation, implying that all the non-trivial properties of the discrimination (choice accuracy and the shape of the RTD together with its dependence on difficulty) depend exclusively on ILD, are fully specified by the parameter Γ, and are invariant with respect to changes in overall stimulus intensity. The temporal variable *τ* in Eq. 1 is dimensionless. The time *t* in seconds in the actual discrimination process is related to *τ* as *t* = *t*_*θ*_ (ABL) *τ*, where the effective unit of time is given by

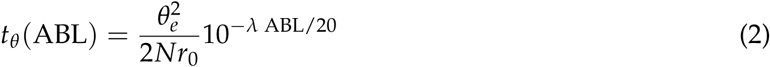

## Quantitative model fits

By construction, this model obeys WL and produces scale-invariant RTDs. It is possible, however, that the model is still unable to fit the data quantitatively (e.g., if the model produces RTDs different from the measured ones or if it fails to capture how exactly they change with ILD and ABL), specially considering that all observed quantities (except for the effective unit of time of the RTDs) depend on the single parameter Γ, and that there are twelve experimental conditions (four difficulties and three ABLs) whose full RTD we are aiming to account for. Since the measured RTs contain stimulus-independent delays (non-decision time – *t*_*NDT*_; Fig. S3) in addition to the decision-time specified by the model, we added two parameters describing the mean (*t*_*ND*_) and variance (*s*_*ND*_) across trials of *t*_*NDT*_. We used a ‘constrained’ model fitting approach (see Methods, Fig. S7) designed to challenge the model’s predictive power. Briefly, we extracted all the ILD dependence of the problem from fits to accuracy only (without using RTs), so that the shape of the RTDs and their ILD dependence are all model predictions. We then used RTs to infer the effective units of time for each ABL, but we only used data from the ABL = 20 and 60 dB SPL conditions, so that everything about the RTDs at ABL = 40 dB SPL is also a prediction.

The model’s fit at the group level are shown in Fig. 4 (see Fig. S9 for fits to the data from each rat individually). The single-parameter fit to accuracy is excellent, accounting for more than 99.8% of the variance in the rats psychometric data (Fig. 4A). Using the fitted value of Γ, the just-noticeable-difference (JND) of our rats is *∼* 2.3 dB (see Table S2), which is similar to that measured in other species (42–44). The quality of the fits to the RTDs for all twelve conditions (Fig. 4B-C) is even more impressive. Once the single number *t*_*θ*_ (ABL) (Eq. 2) for a given row in Fig. 4C is fixed by the fit, both the shape of the model RTDs as well as their dependence on difficulty is completely determined by Γ, which only depends on the psychometric function. Nevertheless they match the experimentally observed RTDs with remarkable accuracy. No RT-data was used from our ABL = 40 dB SPL condition, and still the model accurately predicts the RTDs for this intensity. Eq. 2 also makes a quantitative prediction about the exact functional dependence between ABL and RTs, which we verified is also consistent with our measurements (Fig. S8). The best-fit parameters are shown in Table S2, which also shows results for alternative fitting methods, all giving essentially identical results. The power-law exponent *λ* = 0.099 *±* 0.003, indicates a large degree of compression, which results in a relatively mild-dependence of RTs with ABL. The value of *θ*_*e*_ = 42.2 *±* 1.5 implies that each sensory spike contributes a few percent of the evidence necessary to reach threshold.

**Figure 4:**
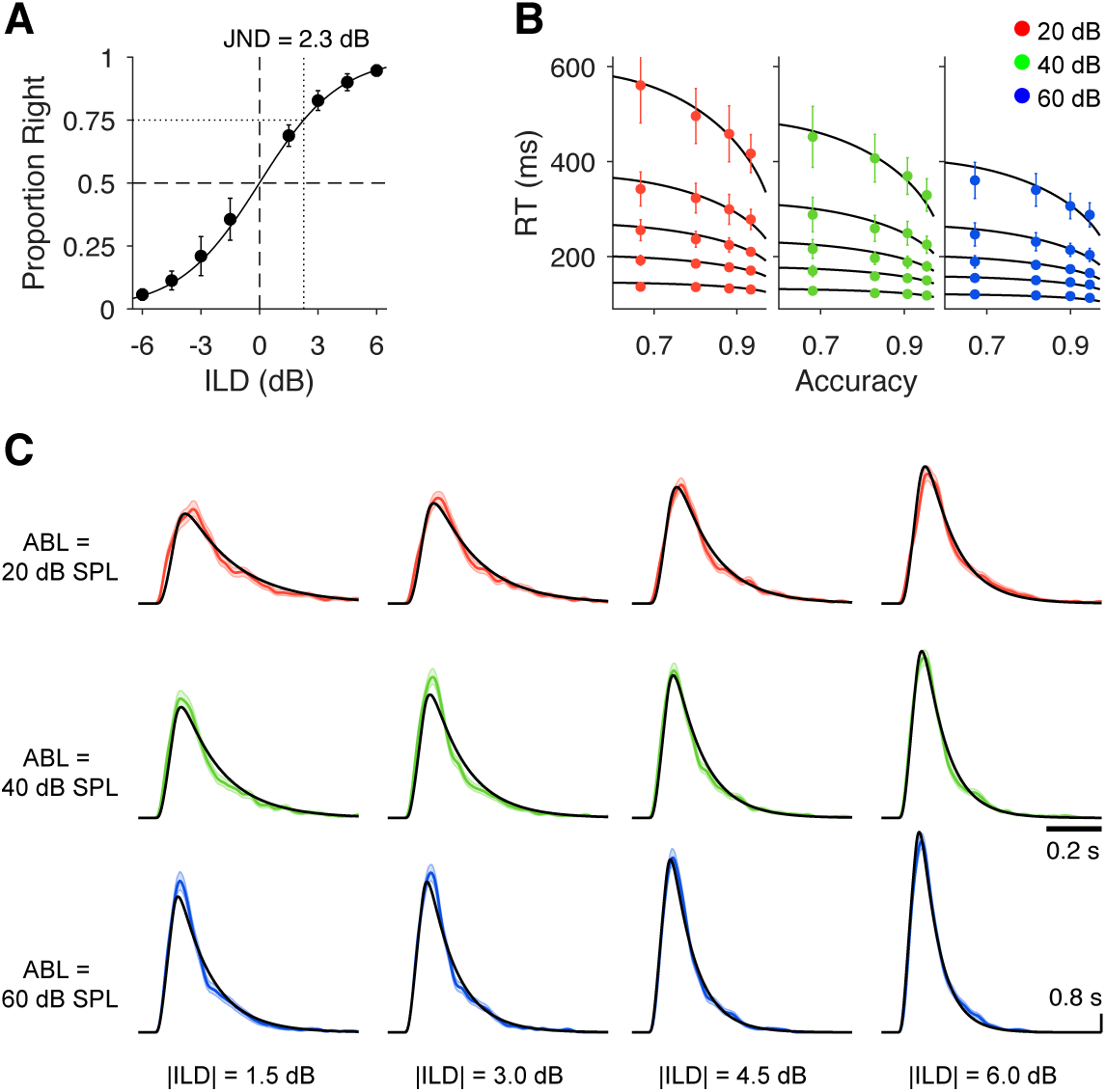
Model fits. **(A)** Psychometric function. Black circles show choose-right probabilities (mean *±* SD across rats). For each rat, the responses to the three ABLs where averaged. Line shows the best fit to this data from the single-parameter (Γ) model in Eq. 1. **(B)** Circles show the 0.1, 0.3, 0.5, 0.7 and 0.9 quantiles (mean *±* SEM across rats; Model fits for individual rats are shown in Fig. S9) of the RT distribution. Each plot is one ABL (same color code as in Fig. 2). These are the statistics that were used for fitting the remaining four model parameters (see text and Methods). Black lines represent model fits **(C)** RT histograms for each ABL and difficulty (color) and for the best fit model (black).

To validate our interpretation of the parameters, we performed further controls and analyses to prove that the rats’ behavior is fully consistent with our implicit modeling assumptions (Fig. S10). In particular we show that rats can generalize ILD discrimination to new sounds (Fig. S11B), suggesting that they fully understand the task contingencies (Fig. S11A), that performance-limiting noise has an internal origin (Figs. S11C), and that only the nature of the sound on a given trial is predictive of the rat’s behavior on that trial, i.e., that the rats display virtually no history-effects or sequential dependencies (Fig. S12) that could contaminate our estimate of the JND.

## How is the JND established?

The JND in our model depends on the power-law exponent *λ* and on the decision threshold *θ*_*e*_ (Eq. 1, Supplementary Information). The exponent *λ* presumably reflects signal-transduction biophysics and is thus not obviously plastic. The threshold *θ*_*e*_ on the other hand, similarly to the criterion of SDT, is usually assumed to be under the subject’s control and to depend on the value of outcomes (4, 13, 35, 40, 41), which, if true, would put the JND under the control of the subjects. The JND, however, is usually assumed to be a property of the sensory system and, in fact, we see very little variation in the value of the JND of our rats (essentially no variation in four out of five; Table S2). Why are the JNDs so similar and, more generally, why do they take their observed values?

One possibility is that the evidence threshold may be in some sense optimal and thus specified by the task contingencies. To explore this hypothesis, we computed the value of the threshold that would maximize reward-rate (see Methods). The optimal threshold is around twice our estimate for *θ*_*e*_ (Fig. 5A), casting doubts on the optimality of the measured decision bound. Our estimate of the optimal threshold could, however, be unreliable, because the reward-rate may depend on costs that are difficult to identify, e.g., an explicit cost of time (41) (although we note that, because RTs vary with ABL, no single threshold can maximize reward-rate for all stimulus intensities; Fig. 5A). To address this difficulty, we decided to test whether the threshold could, at least, be locally adapted to the task contingencies (a necessary condition for optimality).

**Figure 5:**
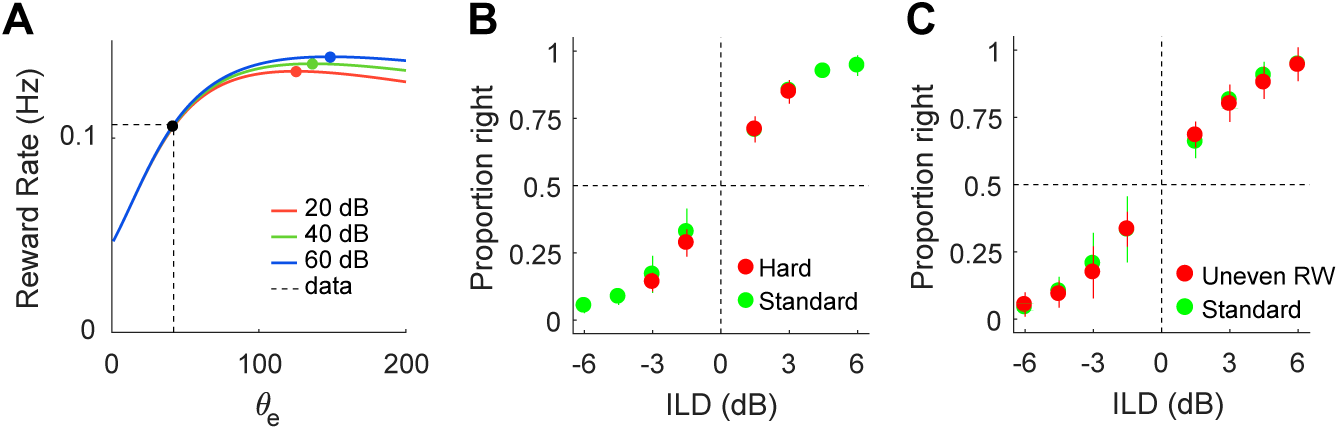
Optimal decision thresholds and role of motivation. **(A)** Curves show reward rate as a function of decision threshold for each ABL separately (see Methods). Filled colored circles are the optimal decision thresholds. Dashed line shows the actual fitted decision threshold. **(B)** Choose-right probability (mean *±* SD across rats) for hard versus control blocks. **(C)** Choose-right probabilities (mean *±* SD across rats) for standard blocks and for ‘uneven reward’ blocks where rewards for correct choices in the two hardest (easiest) conditions are 20% larger (smaller). See Methods for details on task manipulations.

We attempted to drive increases in accuracy which could be taken as evidence of raised decision thresholds by increasing the motivation of the rats. First, we presented blocks with only the two most difficult conditions, but we did not observe changes in accuracy (Fig. 5B, comparison of *d*’ between only hard and control conditions: *p* = 0.5, Fisher’s exact test; see (45) for different results on a similar manipulation). We also varied the reward magnitude as a function of difficulty on a series of sessions, making rewards larger (smaller) for the two hardest (easiest) conditions, but this also failed to produce any changes in performance (Fig. 5C, comparison of *d*’ between uneven reward and control conditions: *p* = 0.9, Fisher’s exact test). Subsequent longer-lasting bi-directional manipulations of motivation (inducing changes in reward rate of more than 100%) produced the same results (Fig. S13; see Table S3 for statistics for all behavioral manipulations). We conclude that the evidence bound is set to a fixed value for a yet-unknown purpose, and that this value of the bound in turn sets a hard-limit on discrimination accuracy.

## Discussion

Although the importance of Weber’s law derives from its generality across sensory modalities and species (5, 6) we believe that it is particular features of our task and model system that allowed us to establish such an accurate specification of the mechanism driving the behavior. In an ILD discrimination with respect to the midline, subjects report whether a sound is lateralized to the left or right. It is only from our knowledge of the auditory system that we can interpret this report as one about the relative intensity of the sound at the two ears (28, 46). We are thus recruiting a circuit designed by evolution for the purpose of comparing stimulus intensity (46), rather than trying to create or co-opt a general purpose comparison mechanisms with no particular significance for the rat. Furthermore, in an ILD discrimination with respect to the midline the categorization threshold is hard wired (46,47). We believe that it is because of this fact that rats appear to genuinely understand the contingencies of the task (Fig. S11), display negligible lapse rates (Figs. S11B-C, S13A), and be purely driven by the sensory stimulus (Figs. S10, S12). The lack of alternative sources of control over the behavior, such as trial-history effects (Fig. S12), is unusual, even in humans (48). Rats are able to sustain motivation and behave reliably and consistently for the many thousands of trials that are necessary to precisely quantify the small but systematic differences in RT that are key to our findings. Our results demonstrate that rats are an excellent model organism for RT psychophysics.

Although many models to explain Weber’s law have been proposed, most previous models and experiments (with exceptions (4, 8, 9); see below) are cast within SDT (as opposed to SS) and thus do not consider RT. Within SDT, subject behavior follows the properties of the evidence at a certain imposed time (the stimulus duration), whereas within SS, behavior derives from imposing that the evidence reaches a certain value (at an internally derived time which sets the RT). This distinction becomes relevant in comparing our results with those of Brody and colleagues, which in a series of elegant studies have been dissecting how rats discriminate the rate of lateralized discrete pulses of sensory evidence in a fixed (but variable) duration task (25, 49). Although we find, like them, that a key computation in sensory discrimination is ‘perfect’ (i.e., without time decay) accumulation of evidence, there are also important differences. In the latest instantiation of their model (50), they show that their data is well described by a SDT model with ‘scalar’ evidence statistics (SD proportional to mean; see also (8)), whereas in our model it is the variance of the evidence that is proportional to the mean – indicative of statistical independence of the evidence across time. Both (9, 51) and our results show that, within SS, scalar (in fact, perfectly sale invariant) behavior is compatible with temporally uncorrelated evidence samples. In contrast, using a SDT model forces the evidence to have the same statistics as the behavior. Furthermore, it is not clear how to mechanistically implement scalar statistics at the level of the evidence. Although Scott et al. (50) do not explicitly address WL, they show that, consistent with WL, behavior is not invariant with respect to additive changes in the evidence. It would be interesting to test whether discrimination accuracy depends only on the ratio of the number (or rate) of pulses, as WL would predict. It is possible that the brain uses fundamentally different mechanism to accumulate trains of evidence pulses (a stimulus with stochastic temporal variability) and evidence about the intensity of ‘constant’ stimuli. However, the differences between the behavioral report in the two tasks (choice at different fixed durations vs choice and RT) makes it difficult to quantitatively compare the results of Brody and colleagues with ours. Finally, more unspecific but important properties of the two tasks also appear different, as our rats in the RT task neither lapse (see Figs. S11B-C, S13A) nor display sequential dependencies (Fig. S12).

Although SS approaches to perception have a long history (16, 17), it was Link (4) who emphasized a connection between SS and Weber’s law, highlighting the importance of Poisson variability (see (8) for a thorough analysis of the connection between RT and WL within SS but with non-Poisson evidence statistics). In an important recent study, Simen and colleagues predicted that WL should be associated to scale-invariance of the RTD (9), highlighting how this unifies discrimination and temporal estimation, where scale-invariance of response times is well established (52). The evidence presented in support of the claim, however, is inconclusive, even at the level of establishing WL (perhaps because pure tone discriminations were analyzed, which are known to be one of the few exceptions to WL (53)). Moreover, Simen and colleagues did not attempt a quantitative match between their model and data.

Our work extends the contributions of Link and Simen et al. (4, 9) in important ways. First and foremost, we establish unambiguously and for the first time that, at least in our task, WL arises because changes in stimulus intensity have the exclusive effect of a uniform rescaling of the RTD. Together with the breakdown of WL for short durations, this strongly suggests that WL is a feature of a process of bounded accumulation of evidence. In addition, we show that the scale-invariance of the RTD provides a strong constraint that allows the identification of the computational mechanism at work qualitatively, without resorting to model fitting. Finally, we prove that the effectively simplest implementation of the identified mechanism displays a level of accuracy at describing the behavior of the subjects that is highly unusual. The accuracy of the model is even more remarkable given that all RT-related properties of the model (except for the effective unit of time for each ABL) are pure predictions based only on a single-parameter fit to the rats discrimination accuracy. Globally, all of these facts constitute, in our opinion, very strong evidence that the mechanism initially proposed by Link (4) and refined by us, is in fact the correct explanation for Weber’s law.

Whereas scale-invariance of the RTD for stimuli of different overall intensities is a new finding, previous work has pointed out the constancy of the coefficient of variation of the RTD with respect to changes in stimulus difficulty (54–56), another form of scale-invariance. In the model, however, the former type of invariance is exact, whereas the latter type is approximate (Supplementary Information; Fig. S6). Remarkably, the RT data allows us to also confirm this model prediction: The accuracy of the scale-invariance as a function of difficulty (but not ABL) degrades with the difference between the raw, unscaled RTDs (Fig. S4). We believe that this trend is generic: any mechanism producing approximate scale-invariance will tend to work less well as the unscaled RTDs become more different. This lends further support to the mechanism considered here, which generates exact ABL-driven scale-invariance. Interestingly, it is well known that more intense sensory stimuli are perceived as lasting longer (57), in line with the intensity-driven change in the units of time that we report. This suggests that estimation of the duration of sensory stimuli may rely on the same neural signals that are used to make discriminative choices about those stimuli (58).

Since the activity of the sensory channels grows with stimulus intensity (RMS pressure in our case), stronger stimulus intensities lead to faster drift rates. This takes the form of a multiplicative gain increase on the discriminative variable (ILD) by sound intensity (Box; Supplementary Information). Intensity-driven changes in gain are common in sensory processing (59–61), and explicit gain-normalization schemes have been proposed to remove the unspecific effect of intensity or contrast (62–64), also in the context of ILD discrimination (42). Gain normalization has also been linked to the contextual effect of stimulus conjunctions in value-based decision making (65) and attention (66). In our model, the multiplicative effect of intensity on the strength of evidence does not have any consequences in terms of discrimination accuracy without the need of any explicit normalization mechanism. Although drift rates increase with ABL, so does the noise in the trajectories, in a way that leads to exactly the same probabilities of hitting either bound. The higher drifts and noise, however, lead to earlier threshold crossings, which substantiates the ABL dependence of RTs. Typically, a single drift-rate parameter is used to measure strength of evidence (in effect measuring strength of evidence relative to a constant noise level) (35, 41, 55). However, discrimination tasks involve a comparison, which is conceptualized as as a difference in activity between two channels with opposite stimulus tuning, making the problem intrinsically two-dimensional. In our case, we use ILD and ABL as these two dimensions, which is equivalent to a change of coordinates from the activity of the two channels to their sum and difference. Our results clarify what is the contribution of each of these two dimensions to the subject’s behavior in a discrimination task, and when a one-dimensional approximation is appropriate (Supplementary Information; Fig. S5). In practice, including stimuli with different intensities in the experiment removes the freedom to choose an arbitrary value for the noise of the evidence (typically taken as *σ* = 1 or *σ* = 0.1) when using a DDM to fit the data.

Power-law transformations of stimulus intensity can be related to our results in two different ways. First, a variety of sensory receptors have been shown to encode stimulus magnitude as a power-law for a partial but significant portion of their dynamic range (67), although with exponents typically larger than our finding of *λ ∼* 0.1 (Table S2) (e.g., an exponent of *∼* 0.3 has been identified in the cochlea (68, 69)). Evidently, our model is phenomenological, but our results suggest that several cascaded compressive stages underly the representation of intensity. Second, Stevens’ ‘Psychophysical Law’, derived through variety of stimulus estimation approaches, states that the subjective intensity of a stimulus is a power-law of its physical intensity (3) (but see (70)). Although it is tempting to relate the activity of the sensory channels in our model, which is also a power-law of stimulus intensity, to the posited subjective intensity of the stimuli, an accurate interpretation of the results of estimation experiments requires a good model of the behavior, and the computational underpinnings of estimation and discrimination are expected to be different.

Studies on the neural basis of level-invariant ILD discrimination or sound localization have focused on the existence of explicitly level-invariant neural codes (e.g., the same ILD tuning regardless of ABL). Recordings from the lateral superior olive (LSO – the first station in the auditory pathway where ILDs are represented) and the inferior colliculus (IC) under anesthesia have revealed an additive, rather than multiplicative, effect of overall intensity on ILD tuning (with shifts more prevalent in the LSO (71, 72)). In the cortex, intracellular recordings reveal a lack of explicit level invariance (73). Neurons in the auditory cortex are broadly tuned to ILD and azimuth with a tendency towards preferences for sounds louder on the contra-lateral side (74, 75). Opponent-process models (similar in spirit to our proposal) employing subtraction of activity from neurons displaying opposite tuning to ILD/azimuth, allow more accurate and more level-tolerant decoding of ILD than models using the activity of similarly tuned neurons (72, 74, 76). Our results show that explicit level invariance in neural representations is not necessary for level-invariant accuracy at the behavioral level, and can instead be a property of the discrimination mechanism itself, with neither neurons encoding the evidence, nor neurons integrating it, displaying ABL-independent activity. Regarding non-sensory components of the model, the parietal cortex (77–79), and the striatum (80), are candidates for evidence accumulation, and the superior colliculus (SC) has been suggested as a possible site where a threshold-like mechanism might be implemented (81, 82). Future studies should address the involvement of these areas in tasks probing the effect of stimulus intensity on sensory discrimination.

Our failure to drive increases in performance through strong manipulations of motivation (Figs. 5, S13) suggests that rats in our task appear to have reached the limits of discrimination accuracy imposed by their sensory organs. However, their behavior is almost perfectly explained by a model which contains no such hard limits. Indeed, getting the model to perform better is trivially accomplished by raising the evidence threshold, something our rats appear incapable of doing even if strongly motivated to do so. It remains to be elucidated how the fixed evidence threshold is set and under which conditions motivation ceases to have control over its value. Although our experiments argue against an optimal value of the threshold in the context of imposed task contingencies, its value may still be adaptive in the face of longer-lasting developmental or evolutionary constraints.

Do our results shed any light on a normative explanation for Weber’s law? In ILD discrimination, intensity ratios specify angles in the horizontal plane (azimuth), since the attenuation of sound intensity by the head for a given azimuth is itself a ratio (i.e., it’s specified in dB). Thus, ratiometric intensity comparisons would seem adaptive as cues for the localization of sound. May this extend to other modalities? In vision, information about the environment is based on the relative reflectance of different surfaces. If the intensity of a light source changes, the relative luminance of different patches of an image stay constant, even if their absolute difference changes. In olfaction, molar ratios in a chemical mixture remain constant when the absolute concentration of the mixture is altered, and discrimination accuracy of mixtures depends on ratios (83). Thus, we hypothesize that the ratio-sensitivity of behavior which Weber’s law embodies is an adaptive strategy to extract information from the environment in a way that is invariant of the intensity of the sources.

## Materials and Methods

### Experimental animals

All procedures were carried out in accordance with European Union Directive 86/609/EEC and approved by Direcao-Geral de Veterinaria. Experiments were performed on 15 adult female Long-Evans hooded rats. Animals were 12-13 weeks old, weighted between 250 and 300 g at the beginning of the experiments, and were kept above 85% of the initial weight. All animals were naive to any behavioral tests. Rats had free access to food but water was restricted to the behavioral sessions, which were conducted during five consecutive days per week; animals had access to water during the sixth day and were water deprived for 24 hours before each round of five sessions. All results in the main text except for Fig. 3 came from the same batch of rats (Batch A; see Table S3). A second batch of animals (Batch B) was used for the behavioral manipulations shown Figs. 3, S11C, S13. The five rats in Batch A were tested in the reaction time sound lateralization task. Four of them performed blocks including only the hardest conditions, and three of them performed blocks with uneven RW and blocks with pure tones. The six rats in Batch B were tested in the bidirectional motivation manipulations and in the frozen noise manipulations. Four of them were tested in the capped sound duration sessions.

### Auditory Stimuli

A percept of lateralization was created by presenting broadband (5 to 20 kHz) noise with different intensities to each ear (interaural level difference, ILD). The noise was cosine-ramped and independently generated for each ear and for each presentation using a Tucker-Davies Technologies RP2 module at a sample rate of 50 kHz. The effect of loudness on sound localization was assessed by using different average binaural levels (ABLs). After training the animals with gradually smaller ILDs (see below) at an ABL of 50 dB, they were tested with a set of stimuli comprising eight ILDs, ranging from −6 to 6 dB linearly spaced in 1.5 dB steps, and three ABLs: 20, 40 and 60 dB SPL (for the main task; see below for task variants). Negative ILD values indicate higher sound intensity at the left ear; for example, an ILD of −6 dB for an ABL of 40 dB consisted in presenting the noise at 43 dB to the left ear and 37 dB to the right.

The headphones were calibrated weekly, using a Brüel & Kjaer Free-field 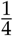 in microphone, placed in front of the speaker, 5 mm apart. For the training phase, the arena speakers were calibrated with the same microphone, placed in front of the central port, facing the speaker to be calibrated.

### Behavioral apparatus and headphone design

Rats were trained and tested on the sound lateralization task (Fig. 1A-B) using a standard Coulbourn Instruments modular box (30 × 25 × 30 cm). All components of the behavioral setup were connected to a RP2 module and accessed by a computer running Matlab 2012b (www.mathworks.com) using TDevAcc controls. The behavioral setup was placed inside a soundproof box, illuminated by infrared lights and equipped with an infrared camera to observe the animals during the sessions. The behavioral box had three ports (nose-pokes), made of stainless steel and equipped with infrared sensors, in one of the walls. The central port was used to initiate the trial and to keep the sound playing. The two lateral ports, equipped with a water spout, were used for the animal to communicate its choice (left and right ports to indicate the sound was louder on the left and on the right, respectively) and to deliver a drop of water (28 *μ*l; reward) after correct choices. The nose-pokes were conical (2.5 cm wide in the outer extreme, and 1.5 cm deep) and the distance between the central and the lateral ports was 8 cm (center to center). Two speakers (arena speakers) were placed above the lateral ports and were only used during the initial training (see below). From then on, stimuli were played trough custom made, detachable headphones.

Both the base to be implanted in the skull and the structure of the headphones were designed using Sketchup (www.sketchup.com) and 3D printed in VisiJet(R) EX200 Plastic. The structure consisted of different components that could be adjusted for each individual rat and an enclosure that fitted the size and shape of the speakers. The speakers were aligned with the ears and placed at 5 mm from the entrance of the ear canal. Once adjusted, all pieces were glued together and remained fixed throughout the experiment (Fig. S1A). The headphones were attached to the base at the beginning of each behavioral session and detached before taking the animal back to the holding cage. Our speakers were Knowles receivers (model number 2403 260 00029), which were small enough to fit in our headphone design.

## Behavioral Tasks

### Sound lateralization task: Temporal and outcome contingencies

Rats started a trial by poking in the central port within a 6 s time window (start trial waiting time) triggered by the end of the inter-trial interval (ITI, 3s), which was signaled by a light in the box turning off. After a short, variable fixation time (FT, 300-350 ms) the sound was played binaurally, through custom-made headphones, until the rat left the central port or until the maximum presentation time (6 s) was reached. Rats had to communicate, within a 2 s time window (response waiting time), whether the sound was louder at the left or right ear by poking with the snout in either the left or right ports, respectively. Correct choices were rewarded with a drop of water (28 *μ*l) and incorrect responses penalized with a 10 s timeout during which the rat was not able to start a new trial. Trials in which the rat failed to start a new trial within the start trial waiting time, broke fixation during the FT, or failed to poke in either lateral port withing the response waiting time, were considered aborts. Aborts were repeated after a 1 s time penalty.

Each session was divided in blocks of 80 trials. Within each block, the ABL was kept constant, while the ILD changed pseudo-randomly from trial to trial. Typically, sessions lasted for two hours and rats performed between 800 and 1200 trials.

### Sound lateralization task: Training

Animals were initially trained in a simplified version of the task, in which fully lateralized sounds (50 dB SPL broadband noise) were presented from either of the arena speakers. Rats understand the basic contingency of the task quickly, within a few hundred trials (Fig. S2A). The sound was played until the animal entered one of the lateral ports, and errors were repeated immediately. Short fixation times and long waiting times were used to increase the chances for the rat to complete the trial while exploring the box. Every time the rat completed a trial, the fixation time was increased by 1 ms and, once the rat completed three consecutive blocks (120 trials per block) with less than 30% abort rate, waiting times were set to their final durations and ILDs were introduced. Initially, the ABL was set to 50 dB and the ILD step was set to 4 dB; depending on the animals performance, the step was decreased gradually until the final 1.5 dB (Fig. S2B).

Once performance with 50 dB ABL and 1.5 dB ILD step was stable, the magnetic base for the headphones was implanted and the animals were allowed to recover for at least one week, during which they had free access to food and water. After this period, animals were tested, with headphones instead of the arena speakers, in the final stimuli set (24 conditions; 3 ABLs × 8 ILDs). All transitions between training steps were smooth; all rats generalized across stimuli sets and switching from arena speakers to headphones had little impact on performance even for the first block of trials with headphones.

### Block types used for the different tasks

In order to test various hypotheses about the nature of the behavior, we modified the basic task above in several ways. Rats from Batch A were tested in 5 types of blocks: (A1) “standard” blocks, in which 4 ILDs (of each sign) linearly spaced from 1.5 dB to 6 dB steps were presented. All data in the main text except for Figs. 3 and 5 came from A1 blocks. (A2) “hard”, in which only the four ILDs closer to the mid-line (*±*1.5 and *±*3 dB) were presented (Fig. 5B). We did not present only the hardest condition because when we attempted this rats “gave up” and became biased. (A3) “uneven RW”, in which we increased (decreased) the amount of water delivered after correct discriminations for the two hardest (easiest) ILD conditions by 20% (Fig. 5C). (A4) “log noise”, in which 5 ILDs (of each sign), logarithmically spaced between 1 and 8 dB, were used (Fig. S11B). (A5) “log pure tones”, identical to the previous one but using 10 kHz tones instead of broadband noise (Fig. S11B). In all types of blocks except for A5, three ABLs = 20, 40 and 60 dB SPL were used. For blocks of type A5, only ABL = 60 dB SPL was used. Rats from Batch B were tested in 6 types of blocks always with ABL = 50 dB SPL: (B1) “standard”, in which 4 ILDs (of each sign) logarithmically spaced between 1 dB and 8 dB were used (Fig. S11C, S13). (B2) “hard”, in which only the two most difficult conditions (1 and 2 dB) were used (Fig. S13). (B3) “easy”, in which the two easiest conditions (ILD = *±* 4, 8 dB) were used (Fig. S13B-D) (B4) “frozen noise (FN) coherent”, in which the exact broadband sound (out of four different examples) was presented in the two headphones appropriately scaled to produce a given ILD (ILDs were the same as in the standard blocks). (B5) “FN non coherent”, in which a different one of the four examples was played in each headphone (same ILDs). B4 and B5 where used in Fig. S11C. (B6) “easy sessions”, in which only ILDs = *±* 4.5, 6, 9, 15 dB were used (Fig. S13A). After these manipulations, four remaining rats from this batch were initially trained in a variant of the standard RT task with a different set of conditions (ILD = 0, 0.5, 1.25, 2.25, 4 and 8 dB and ABL = 10, 25, 40, 55, 70 dB SPL) designed to sample effect of difficulty and intensity more densely, and without blocks. After a few sessions we decided to perform the capped sound duration experiment and we then switched conditions to ILD = 1, 2, 4, 8 dB (of each sign) and ABL = 20 and 40 dB SPL. The range of maximum sound durations (SD_max_) tested was 50, 75, 100, 140, 150, 180, 220, 250 and 260 ms. These sessions are used in Fig. 3.

### Task variants: Discrimination of the ILD of pure tones

As a control to rule out stimulus-response associations as determinants of performance, and to test generalization, three of the rats were tested with blocks of type A4 and A5 mixed pseudo-randomly.

### Task variants: Manipulations of motivation

We performed several types of task manipulations to study whether and how changes in motivation would affect performance in the task. For rats in Batch A, blocks of type A2 where randomly included among standard blocks within a set of sessions, in such a way that every time ‘hard’ conditions were tested, at least three consecutive blocks (one for each ABL) were used. Results from this manipulation are in Fig. 5B. These same rats also performed a series of sessions where all blocks were of type A3 (Fig. 5C). Longer-lasting and bi-directional manipulations of motivation were tested with rats of Batch B. These rats were tested in a series of sessions where an initial standard block was followed by blocks of type B2 until the end of the session, and on sessions where only blocks of type B6 (easy and very easy trials) were used (Fig. S13A). They were also tested in a series of sessions where motivation was manipulated bidirectionally within a single session: after a standard block, two blocks of type B2 (or B3) were used, then another standard block was used, then two blocks of type B3 (or B2) we used, and so forth, until the end of the session (Fig. S13B-D).

### Task variants: External vs Internal noise

To test the contribution of external noise to the width of the psychometric function, we selected four broad band noise samples and either used the same or different samples at the two ears. Rats of Batch B were tested in a series of sessions with blocks of type B4 and B5 alternating pseudo-randomly within the same session (Fig. S11C). It should be noted though that because our headphones are not inserted in the ear canal, we can’t fully control that the waveforms at the two ears are completely coherent, but at least we can restrict a possible contribution of external variability to the changes in the same exact waveform arising from differential interaction with the pinnae.

### Task variants: Maximum sound duration

We investigated whether rats could extract the same information from stimuli of the same duration at different intensities by capping the maximum stimulus duration (Fig. 3). The task was still in reaction time configuration (rats choose when to leave the central port freely) but, for a given SD_max_, the sound was stopped at that SD_max_ if the rat had not left the central port at that time. For choices with RT *<* SD_max_, the sound offset was triggered by the central port exit. Rats performed the task in mini-blocks of 16 trials. Each mini-block contained two permutations of all the ILDs at fixed ABL and SD_max_, and ABL and SD_max_ were chosen randomly from their possible values each mini-block.

## Data analysis

### Isolating stimulus-dependent reaction times

In order to exclude those trials in which behavior was not driven by the stimulus, we looked for the minimum RT for which there was evidence of condition-dependence. To this end, we used two-sample Kolmogorov-Smirnoff test to compare the distribution of RTs corresponding to the two conditions with shortest (ILD = 6 dB, ABL = 60 dB SPL) and longest (ILD = 1.5 dB, ABL = 20 dB SPL) mean RT. Starting at 50 ms, we systematically included longer and longer RTs. As evident in Fig. S3B, if the maximum RT is sufficiently short, the two distributions are not significantly different but for RTs *>* RT_min_ = 90 ms, they become different. For all analyses, we excluded trials with RT *<* RT_min_. In addition, since the shape of the RT distributions is very well behaved and understood in our study, we also excluded trials witch exceedingly large RTs, which presumably reflect disengagement. For all analyses except model fitting, we chose a conservative value of RT_max_ = 1000 ms (the fraction of trials with RT *>* 1000 ms was always very small: 0.39 *±* 0.39 % – mean *±* SD across rats). For model fitting, trials with RTs above the 97% percentile in the RT distribution of each rat were excluded. Empirical estimates of the RT distribution (Figs. 2, 4, S9) were made using kernel density estimation (84) as implemented with custom `matlab` code.

### Accuracy

Accuracy was assessed using the SDT statistics for sensitivity (*d*’) and criterion (c) (13). Completed trials within the valid range of RTs were divided into four categories: hits (correct responses to the left); false alarms (incorrect responses to the left); correct rejections (correct responses to the right); and misses (incorrect responses to the right). Sensitivity and criterion were then estimated by applying the standard z-transform formulas (13).

We used sensitivity and criterion to have a robust measure of accuracy within each 80 trial block. Level-dependence was tested by comparing *d*’ and c as a function of ABL (three comparisons: 20 vs 40 dB; 20 vs 60 dB; and 40 vs 60 dB) using Fischer exact test (i.e., testing the different in the means between two arrays (with elements given by the corresponding statistic *d*′ or *c* in each block) corresponding to a pair of conditions, with respect to a null distribution of no-difference obtained by randomly shuffling the conditions labels 20000 times). Alpha level was set to 0.05 and adjusted using Bonferroni correction for multiple comparisons.

Because “easy” blocks barely contain any errors, for comparisons between “easy” and “standard” blocks (Fig. S13) the statistic was % Correct instead of *d*’, and significance was evaluated again using Fisher’s exact test.

In Fig. 3A we fit a two-parameter logistic function of the mean across rats of the percent correct of as a function of abs(ILD) (difficulty). In addition to the slope parameter, we included an asymptote because we observed that rats in the SD_max_ sessions sometimes display not-negligible lapse rates (unlike in the RT task).

For the calculation of *d*′ in Fig. 3 we did not use 80-trial blocks for the statistics since the task only involved mini-blocks of 16 trials (too short to estimate *d*′). A single *d*′ was calculated from all trials in each condition (SD_max_, ABL) for each rat. Since the SD_max_ sessions and the just-preceding RT sessions used different values of ILD (see above), we fit a four-parameter logistic function (two asymptotes, slope and bias) to the behavior of each rat on the RT sesions. We then estimated the performance of each rat at the same values of ILD used in the SD_max_ sessions and created datasets with the same number of trials as the actual RT sessions but with ratios of correct and incorrect trials consistent with the interpolated performances. These new datasets were then used for calculating the *d*′ of the RT sessions. Finally, to estimate the uncertainty of the green and red leftmost points in Fig. 3B (RT sessions), we compared within these sessions, for each rat, the performance in trials with ABL = 20 (40) dB SPL with that in trials with ABL = 40 (20) and 60 dB SPL merged.

To quantify the effect of sound duration on difficulty, we fit a sigmoid to the average across rats of the difference of the sensitivities *d*′ (RT) *-d*′ (SD_max_) = *a*([1/SD_max_]^3^)/(*b* + [1/SD_max_]^3^), separately for the two ABLs. We used the inverse of the sound duration in the abscissa to avoid having to arbitrarily assign a SD_max_ to the RT sessions. The parameters *a* and *b* represent the asymptote and the threshold of the sigmoid. In the legend of Fig. 3 we report the threshold in ms, which is equal to *b*^-1^/^3^. The exponent was chosen by inspection because it provided reasonable fits. The fit was made to the average difference in *d*′ across rats (filled circles in Fig. 3B). To assess whether the parameters of the fit where different for the two ABLs, we used the difference in the value of each parameter for the fits corresponding to each ABL as a test statistic. We then generated surrogate datasets by permuting the ABL label of each trial separately for each rat and for each SD_max_. Each permutation generated, for each rat, a surrogate 20 and a surrogate 40 dB SPL dataset (which were composed of trials belonging to the real 20 and 40 dB SPL conditions picked randomly). We then performed fits of each surrogate exactly in the same way as for the real data and computed the value of the test statistic. Repeating this process across 2000 permutations gave us an estimate of the distribution of the test statistic under the null hypothesis of no difference between the two ABLs. The *p*-values reported are two-tailed against this null distribution.

### Reaction Time

Two-way ANOVAs (3 × 4) were used to describe the effects of ABL and ILD on RTs in Fig. 2C reported in Table S1. Group level comparisons were made using Two-way repeated measures ANOVAs.

### Accuracy of temporal rescaling

We assessed how accurately the RTDs for different experimental conditions resembled a uniform scaling of time (Fig. 2D-F, Fig. S4) using the fact that when two distributions are related by a uniform scaling, their quantiles are proportional (Supplementary Information). For fixed ILD (ABL), we regressed the percentiles of each condition on those of the fastest, i.e., 60 dB SPL (6 dB). Since the RTDs are significantly right-skewed, higher percentiles have more uncertainty. Thus, we used weighted least-squares to perform the linear fit. To compute the uncertainty in the percentiles, we generated 1000 bootstrap re-samples from the RTD of each rat for each condition. For each re-sample, we computed the percentiles and averaged them across rats, obtaining a sampling distribution of 1000 percentiles. These distributions are very well approximated by Gaussians. To avoid the complication of having uncertainty in both the dependent and independent variables, we followed the following heuristic approach: We assumed that the independent variable (percentiles of the fastest condition) had no uncertainty, and assigned an uncertainty 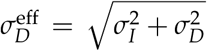 to the dependent variable, where 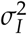 are the variances of the sampling distribution of the percentiles of the independent and dependent variables respectively. We then applied the standard weighted least-squares algorithm to find the value of the slope and its associated *R*^2^, which we report in Fig. 2E-inset, Fig. S4. The shaded areas in Fig. S4 correspond to 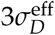.

### History effects

We used logistic regression (85) to investigate the effect of trial history on the performance of the rats in single trials from the standard A1 blocks. We used the model to predict the performance of completed trials within the valid range of RTs. We used 12 predictors from the current trial, corresponding to the four conditions of difficulty (absolute value of ILD) at each ABL (each coded as a [-1, 0, 1] identifier). Four aspects of trial history were evaluated: history of the stimulus lateralization (the sign of ILD regardless of ABL), history of response side, and two response-outcome predictors: history of response side after correct choices, and after errors. For a given trial *i*, the stimulus history predictor for trial *i − j* was coded as a [-1, 1] indicator, where 1 (−1) indicates that the stimulus at trial *i − j* had positive (negative) ILD (by convention, we predict the choose-right probability ‘R’, i.e., the probability of making a response to the side that is correct when ILD is positive). The response history predictor was coded as a [1, 0, -1] indicator, where 1 (−1) indicates that the rat made a response to R (-R) in trial *i − j* regardless of the stimulus and 0 means the rat made an abort. The response after correct (incorrect) choices predictor was identical to the response predictor except that the indicator was set to zero when the trial was either an abort or an incorrect (correct) trial.

For each of these four aspects, we fit a kernel extending 15 trials into the past (0 *< j <* 16 in the previous paragraph), in such a way that the effect of each aspect on the current trial is the convolution of the kernel with the corresponding predictor across trials into the past. We represented all history kernels as linear combinations of five unit-norm decaying exponentials with time constants of [0.5, 1, 2, 4, 8] trials (Fig. S12G inset). In practice, we used an orthonormal basis in the space spanned by these five exponentials. Thus, all in all, our model fits 32 coefficients plus the bias: 12 from predictors associated to the current trial and 5 coefficients for each of the 4 history kernels. All coefficients and kernels for all animals are shown in Fig. S12.

Model fits were performed using the matlab version of the free software package `glmnet` (86) (https://web.stanford.edu/∼hastie/glmnetmatlab/). Regularization was implemented using the elastic net method with parameter *α* = 0.5. All predictors were subject to regularization. All fits used cross-validation as implemented by the funtion `cvglmnet` and we used the `lambda_min` option to select the hyper-parameter that minimizes prediction error. In addition, for assessing the predictive power of the fits, we manually implemented nested cross-validation with five outer folds. Briefly, model coefficients and hyper-parameters were sequentially fit using four fifths of the data and prediction was evaluated on the remaining fifth until all the data had been used both for fitting and prediction. Predictive quality was assessed using the ROC Area Under the Curve (AUC – Fig. S12B).

To estimate the relative predictive power of different types of predictors we used the relative variance of the linear component of the model for a particular set of predictors. After linear combination or convolution for both current-trial coefficients and the four different history kernels (best fits) we obtain five time series across trials. For obtaining a final predictive probability these would be added. Instead, we compute the variance across trials associated to each of the five time series and report its value relative to the summed variance across the five. The fraction of variance for each of the five type of predictors for each rat are shown in Fig. S12A.

We used bootstrap to estimate uncertainty on the model parameters (87). Briefly, we obtained 2000 surrogate datasets from the actual dataset by resampling with replacement. For each surrogate dataset we evaluated predictive accuracy and model coefficients, and from these, we computed 95 % confidence intervals (shown as error bars in Fig. S12).

To assess whether trial history made a significant contribution to the predictive power of the model, we compared the AUC of the actual data to that of shuffled surrogate datasets (Fig. S12B). In each shuffled surrogate dataset, history predictors were randomly permuted across the whole set of trials, with the constraint that the four history predictors associated to a particular trial were in the same position of the permuted sequence for the surrogate dataset. For each surrogate, we estimated the uncertainty in the coefficients again using bootstrap. Trial history was deemed to make a significant contribution if there was no overlap between the 95% CI of the real data and shuffled surrogates.

## Model fitting

Our model is specified by five parameters: *θ*_*e*_, *λ, T*_0_ (the inverse of the effective gain of the pressure-to-rate transformation *Nr*_0_ – see Box), *t*_*ND*_ and *s*_*ND*_. This embodies the assumptions that the decision process is symmetric and that every choice is driven by the stimulus. Since rats don’t display significant lapse-rates or biases we did not need to consider lapse-rate parameters nor parameters specifying non-zero initial conditions or asymmetric decision thresholds. The first four parameters are the absolute minimum number that is required to fit a sequential sampling model (with non-linear encoding of physical stimulus intensity) to data. Including trial-to-trial variability in non-decision time (specified by *s*_*ND*_) led to a small but clear improvement the accuracy of the fits. We used several strategies for evaluating the accuracy of our model.

### 1. Constrained model fitting approach

In order to test the whether (a) the speed accuracy trade-off as a function of ILD, and (b) the dependency of RTs on ABL, which are structural features of the model, were also present in the data, we used an unorthodox ‘constrained’ model fitting approach. Here, we first used choice data to fit the single parameter Γ from the logistic function

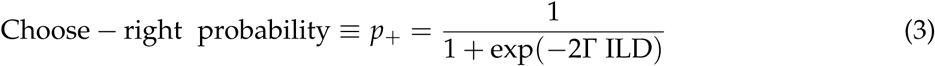

associated to Eq. 1.

Next, we fit the RTDs. We fit the RTDs of correct trials only. Although we attempted to fit incorrect trials as well, the RTDs for incorrect trials, specially for low ABLs, are severely distorted due to the rats anticipation of the sound onset discussed in the text and on Fig. S3. We could not find a principled way of finding and excluding anticipatory error RTs that would lead to well behaved RTDs for errors.

In the constrained fitting approach, we considered Γ was fixed (to its estimated value from fits to accuracy) for fits of the RTDs (implying that all dependence of the RT distributions on ILD was not available for the model fitting process to modify). To further test the ABL-to-RT relationship predicted by the model, we excluded data from the ABL = 40 dB SPL condition for these fits.

RTDs were fit using the *χ*^2^ method for its robustness and efficiency (88). In this method, one uses the *χ*^2^ statistic for a dataset composed of measurements of *N*_*p*_ proportions. We used ten proportions (split by the 0.1, 0.2, 0.9 quantiles of the RT distribution) per condition (although in Figs. 4B and Figs. S7B1-5 we only show the 0.1, 0.3, 0.9 quantiles to avoid visual clutter), and considered eight conditions (the four difficulties for ABL = 20 and 60 dB SPL). Thus, in the constrained approach we adjust the value of four free parameters (*λ, T*_0_, *t*_*ND*_ and *s*_*ND*_) to fit the results for *N*_*p*_ = 80 proportions/data-points. As discussed above, only RTs *>* RT_min_ and RTs *<* RT_max_ were considered.

The cost function for the fit was

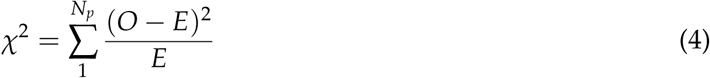

where *O* are the observed proportions and *E* are the expected proportions given the model. To estimate *E*, we used the two series approximations of the RTD of a drift-diffusion process *ρ*_DT_(*t*) for short and long decision-times (89, 90), truncated both series at five terms, and selected between the two according to a threshold in dimensionless time of 0.25, which guarantees a smooth overlap and ensures that the relative error of the approximation is always below 10^−3^ for all decision-times. The distributions were fit in dimensionless time. For a given non-decision time *t*_*NDT*_ and measured reaction time RT, the dimensionless decision-time is *τ*_*DT*_ = (RT *-t*_*NDT*_)/*t*_*θ*_, where *t*_*θ*_ is defined in Eq. 2. The expected proportions *E* were calculated by numerically integrating *ρ*_DT_(*t*) over the value of *t*_*NDT*_, which was assumed to have a uniform distribution of mean *t*_*ND*_ and half-width *s*_*ND*_ (see Box).

The *χ*^2^ cost function was minimized numerically in MATLAB with the function fminsearch, which uses an implementation of the Nelder-Mead simplex algorithm (91). The initial conditions for each parameter were selected randomly within a reasonable interval each time the function was executed, always converging to the same global minimum.

### 2. Unconstrained model fitting approach

We also fit the data in a standard unconstrained fashion. In this approach, we also used the *χ*^2^ method, but in order to fit accuracy and RT simultaneously, we scaled the RTDs so that the area under each distribution would be equal to the probability of the corresponding choice given the model. Accordingly, for each condition, we used 11 proportions, ten for the RTs of correct trials, and a single one for the incorrect trials (in this way, incorrect trials are only used for the estimation of accuracy, see above). Thus, in this approach, we adjust the value of all five free parameters to fit the results for *N*_*p*_ = 132 proportions (eleven per condition for twelve conditions). All other aspects of the *χ*^2^ fit were the same as for the constrained approach.

To obtain error bars for our estimates and an estimate of the joint distribution of the parameters from the fit (Figs. 4B, S7, S8, S9), we used used bootstrap (85), performing subsequent fits for *N*_*r*_ = 1000 resamples (with replacement) from our dataset.

### Maximum Likelihood fits

To ensure that our results were robust to the model fitting method, both constrained and unconstrained approaches were used using the Quantile Maximum Likelihood method (92). This method maximizes the likelihood of the data under the model, but also grouping RTs into quantiles to minimize problems due to outliers. The results of the fits using this method (reported in Table S2) are effectively identical to those obtained using the *χ*^2^ method.

## Optimal thresholds

We computed the reward rate as a function of the decision threshold *θ*_*e*_ (Fig. 5A), assuming other model parameters take on their best-fit values. We define the reward rate *RR* as the expected reward in a trial divided by the expected duration of a trial

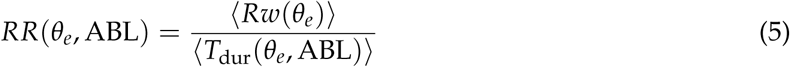

The average reward is

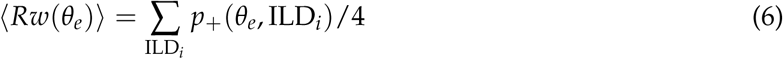

where *p*_+_(*θ*_*e*_, ILD_*i*_) is given in Eq. 3 and ILD_*i*_ = 1.5, 3, 4.5, 6. The average trial duration is

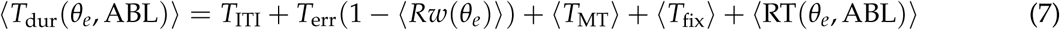

where *T*_ITI_ = 3 s is the inter-trial interval, *T*_err_ = 10 s is the time penalty for an error, *⟨T*_MT_*⟩* = 450 ms is the average movement time of the animals (which we measured empirically), *⟨T*_fix_*⟩* = 325 ms is the mean fixation time, and *⟨*RT(*θ*_*e*_, ABL) *⟩* is the mean RT, given by

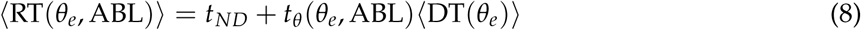

where *t*_*θ*_ (*θ*_*e*_, ABL) is the temporal rescaling factor in Eq. 2 and ⟨DT(*θ*_*e*_) ⟩ is the mean decision time, given by (17, 40)

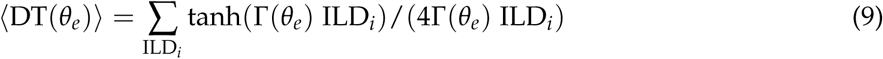

## Acknowledgements

We thank Davide Reato, Jaime de la Rocha, Gautam Agarwal, Eran Lottem, Angel Nevado, Zach Mainen and Leopoldo Petreanu for helpful comments on the manuscript. We thank Patrick Simen for pointing out relevant references, Tor Stensola for help with behavioral boxes, the Vivarium platform at Champalimaud Research for support, Manuel Bayonas for help with headphone prototyping, and Dmitry Kobak for comments on the manuscript and advice on statistical analyses.

## Author Contributions

J.PV and A.R. conceived the project. J.PV and M.V conducted the experiments. A.R., J.C and T.C conceived the theory. J.C, J.PV, M.V and A.R. analyzed the data. A.R. wrote the manuscript with feedback from all authors.

## 1 Supplemental Information

### 1.1 Stimulus delivery and Training

#### The acoustic shadow of the head

We delivered sounds through custom-made headphones (Fig. S1A) which could be easily attached and detached every behavioral session to a magnetic base chronically implanted to the skull (Fig. S1A; see Methods). Throughout this study we’ve assumed that the intensity of the sound at each ear was equal to the calibrated intensity of its corresponding speaker. However, because we decided not to insert the headphones on the ear canal for reproducibility of the stimulus and comfort of the rats, the signal from each speaker will, in principle also contribute to the intensity of the sound at the contra-lateral ear. To investigate if this is a concern, we measured the attenuation (acoustic shadow) produced by the head on an anesthetized rat using the same 5-20 KHz band-pass noise discriminated by the rats, placing our microphone at the entrance of one ear canal and the speaker at its standard position in the contra-lateral side. For these measurements we used high intensities to stay above the noise floor of the microphone. The attenuation due to the head and near-field placement of the speaker is *A* = 22 dB (Fig. S1B-C). Using this value it is straightforward to calculate what is the actual level difference ILD_a_ experienced by the rats as a function of the intended level difference ILD_i_.

**Figure S1.**
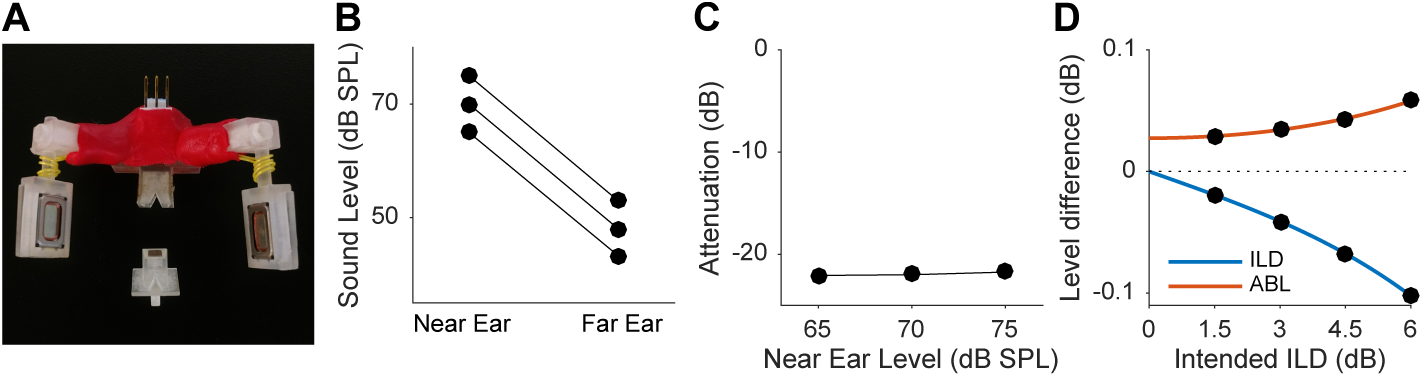
Headphones and acoustic shadow. **(A)** Headphones used for stimulus delivery, and base (bottom). The base is chronically implanted to the skull of the animals and contains a strong miniature magnet which both holds the speakers in place during task performance but also allows easy attachment/detachment. The actual positioning of the speakers was adjusted under anesthesia to match the position of the pinna for each rat individually relative to the base. The red material in the picture is a moldable glue (Sugru) used for strengthening the headphone structure. **(B)** Measurements of the acoustic shadow generated by the head. Broadband noise stimuli were played at 65, 70 and 75 dB SPL and sound level mas measured placing the microphone by the ear canal of the ‘far’ ear. **(C)** The head plus near field positioning of the speakers causes an attenuation of 22 dB. **(D)** Actual minus intended ILD and ABL as a function of intended ILD.

Assuming additivity of the squared pressure RMS from each of the two sources, simple algebra leads to the following expressions

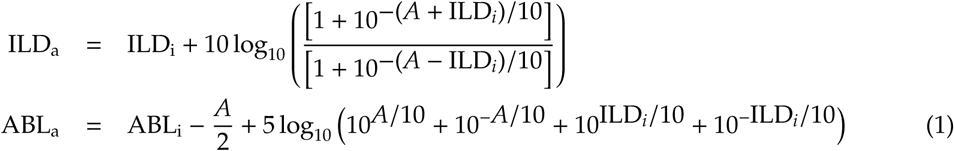

The difference between the actual and intended ILDs and ABLs is shown in Fig. S1D. Since it is always less than 0.1 dB (which is less than an order of magnitude smaller than the just noticeable difference (JND) for ILD of our animals) we have, for simplicity, ignored the difference between actual and intended levels throughout this study.

#### Training

As described in the methods, we trained our animals first with speakers on the sides of the behavioral box before implanting headphones. Withing a few hundred trials of their first exposure with the task, the rats understand the contingencies of the task (Fig. S2A). After this period, we introduced ILDs by playing the stimuli from both lateral speakers. ILD steps were initially large (4 dB) and were made progressively smaller (conditional on performance) until reaching their final value of 1.5 dB. The fact that no significant changes in performance are observed when changing the ILD (Fig. S2B) step further suggests that animals interpreted the stimuli as lying along a continuum (as intended) and did not develop specific behavioral strategies for each stimulus independently.

**Figure S2.**
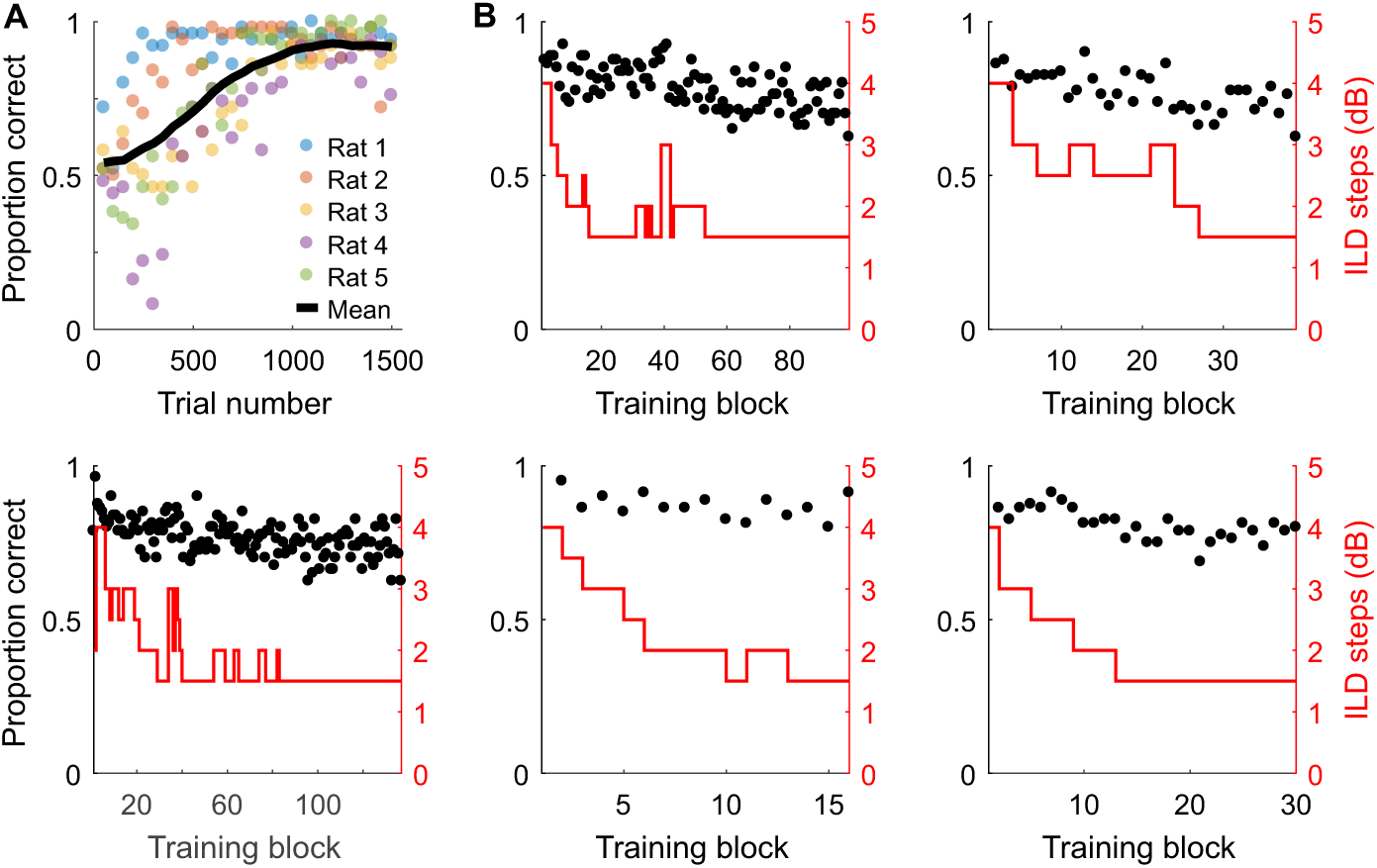
Training. **(A)** Accuracy as a function time in the first sessions where sounds (50 dB SPL broadband noise) come from either of two speakers on the side of the box. **(B)** Taking the animals to psychophysical threshold. Each panel is one rat. Red line is the ILD step used in the block. The smaller it is, the more difficult the task. Black points indicate performance.

### 2 Empirical reaction time analyses

#### 2.1 Short reaction times and intensity-dependent processing delays

The fact that animals could somewhat anticipate the timing of the sound stimulus onset (the pre-stimulus fixation period was uniformly distributed between [300 350] ms) and that we did not impose a minimum reaction time (RT) had the consequence that a fraction of valid completed trials are based on decisions made, and responses initiated, before stimulus onset, but in which sound onset still took place before the rat exited the central port (counting as valid trials). These trials contaminate our results. Fortunately, they have a clear signature at the level of the RT distribution. The cumulative RT distributions are shown in Fig. S3A separately for each difficulty. It is obvious from the plots that the first portion of the distribution comprising the shortest RTs is the same for all conditions and thus stimulus-independent. We estimated the minimum reaction time RT_min_ in which the distributions become stimulus dependent, and focused our analysis only on trials with RT > RT_min_. Although it is likely that in this way we are excluding a small fraction of very fast but still stimulus-driven trials, we decided to use this criterion as it is simple and unambiguous. The very high accuracy of the model fits suggests that the distortion introduced by eliminating these very short RTs is probably quite small.

**Figure S3.**
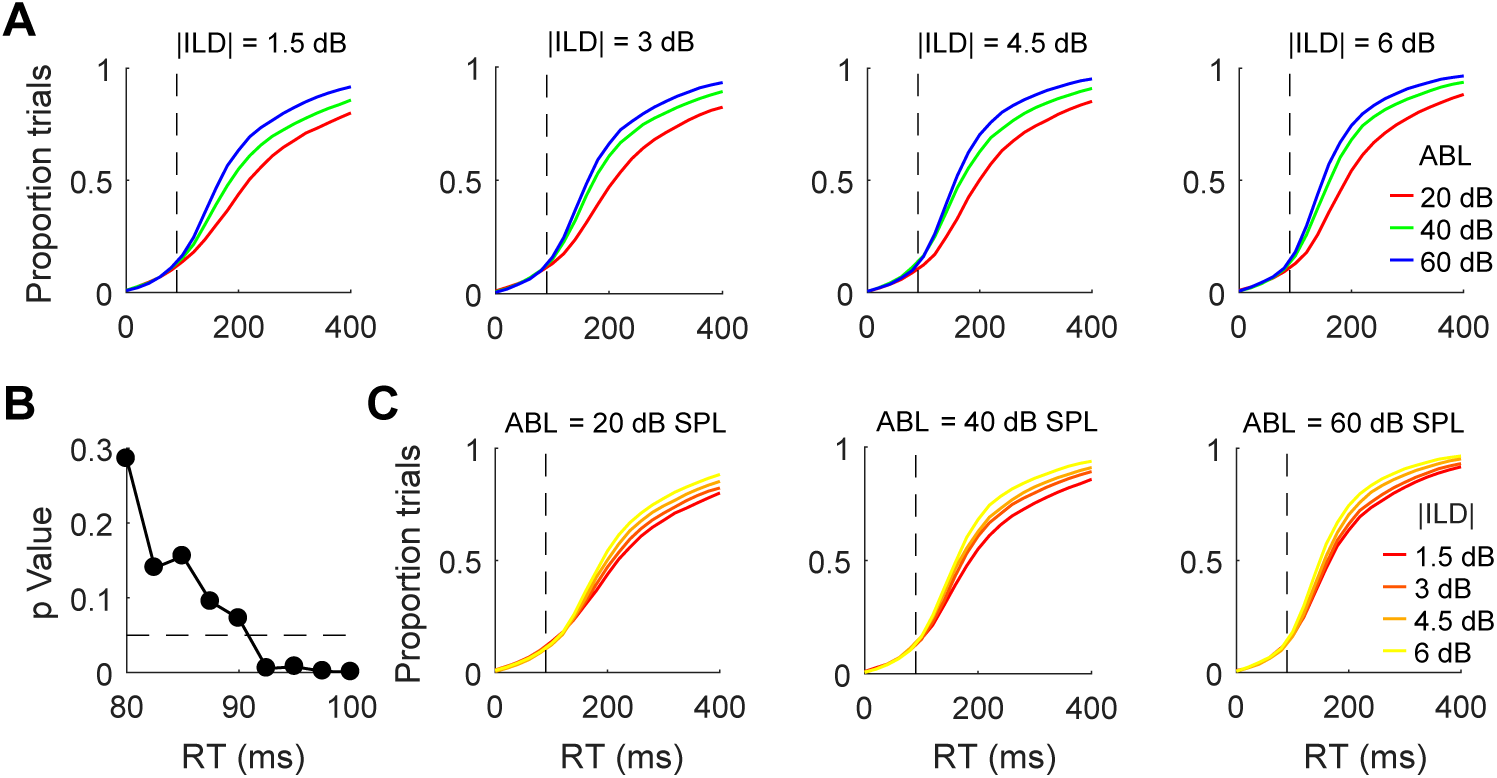
Short RTs and processing delays. **(A)** Each plot shows the cumulative distribution of RTs for a given difficulty (absolute value of ILD) for the three ABLs. The shape of the distribution for the shorter RTs are the same across all conditions. **(B)** *p*-Value of a two tailed Kolmogorov-Smirnoff test comparing the RTs for the fastest and slowest conditions (|ILD| = 6 dB and ABL = 60 dB SPL versus |ILD| = 1.5 dB and ABL = 20 dB SPL) as a function of the maximum RT included in the comparison. We defined RT_min_ as the value at which this comparison becomes significant (90 ms). This value is shown as a dashed vertical line in (A) and (C). **(C)** Same as (A) but grouping the cumulative distributions by ABL. The distributions start to diverge later for fainter sounds.

We estimated RT_min_ as the first RT for which the RT distribution corresponding to the shortest and fastest conditions become significantly different (Fig. S3B). This value is shown as the vertical dotted lines in Fig. S3A,C.

We have argued in the main text that changes in RT as a function of stimulus intensity (ABL) result from a temporal rescaling of the evidence accumulation process. However, an alternative explanation is that this finding simply reflects intensity dependent delays in the auditory pathway. Indeed, neural response latency is typically correlated with stimulus intensity across sensory modalities. Such delays would be expected to extend the period of condition-independent RTs in an intensity dependent manner. Plotting the cumulative distributions grouped by ABL (Fig. S3C) shows that indeed there appears to be an ABL-dependent component to the period of stimulus independence (dashed vertical line is again RT_min_). However, it is also apparent that the rescaling of the RT distributions (in this case as a function of |ILD|, see also Fig. S4B-C) still takes place for RTs > RT_min_ across all ABLs, even for the faintest sounds. Processing delays *per se*, thus, appear not to be sufficient to explain our findings (a result supported by the results in Fig. 3). The intensity-dependence of RT_min_ suggests using a different value for this parameter for each ABL. Accordingly, we estimated an RT_min_ separately for each ABL in the same way as above, and re-did our model fits with ABL-dependent non-decision times, but the quality of the fit improved only marginally at the expense of six new parameters (RT_min_, *t*_*ND*_ and *s*_*ND*_ for each ABL instead of a single value of these parameters across ABLs). Thus, we have kept a single RT_min_ across all conditions.

### 2.2 Empirical assessment of RT scale-invariance

When two distributions in time are related to each other through a stretching of the time axis (a rescaling of time), their percentiles are proportional. More explicitly, assume that distribution ρ′(*t*) obtained from ρ(*t*) by stretching of the time axis by a factor α, so that ρ(*t*) = αρ′(α*t*). It follows that

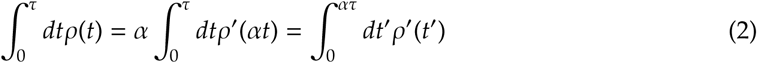

for all τ. If the value of the previous integrals is *Q*/100, then τ is the *Q*^*th*^ percentile *PC*_ρ_(*Q*) of ρ(*t*) and ατ is the *Q*^*th*^ percentile *PC*_ρ_, (*Q*) of ρ′(*t*), i.e., their percentiles are proportional. This fact allows us to use the slope of a linear fit of the percentiles of the two distributions to identify the (putative) temporal rescaling factor α corresponding to a given change in ABL, and the R^2^ of the fit to quantify its precision (Fig. S4A,C). Since difficulty has been suggested to also lead to an approximate rescaling of the reaction time distribution (RTD) (1–3), we performed the same analysis for pairs of distributions corresponding to different difficulties (for each ABL separately; Fig. S4B,C). We assessed the uncertainty in the estimation of of each percentile using bootstrap (see Methods). Since the uncertainty clearly grows with the percentile (4) (Fig. S4A-B), we used weighted least squares to perform the linear fit. Although the fits regress the percentiles of the two RTDs in question, in the Figure we actually plot *PC*_ABL_(*Q*) - *PC*_60dB_(*Q*) as a function of *PC*_60dB_(*Q*) (in panel A, and *PC*_abs(ILD)_(*Q*) - *PC*_6dB_(*Q*) as a function of *PC*_6dB_(*Q*) in panel B), which illustrates the linear dependency more clearly.

**Figure S4.**
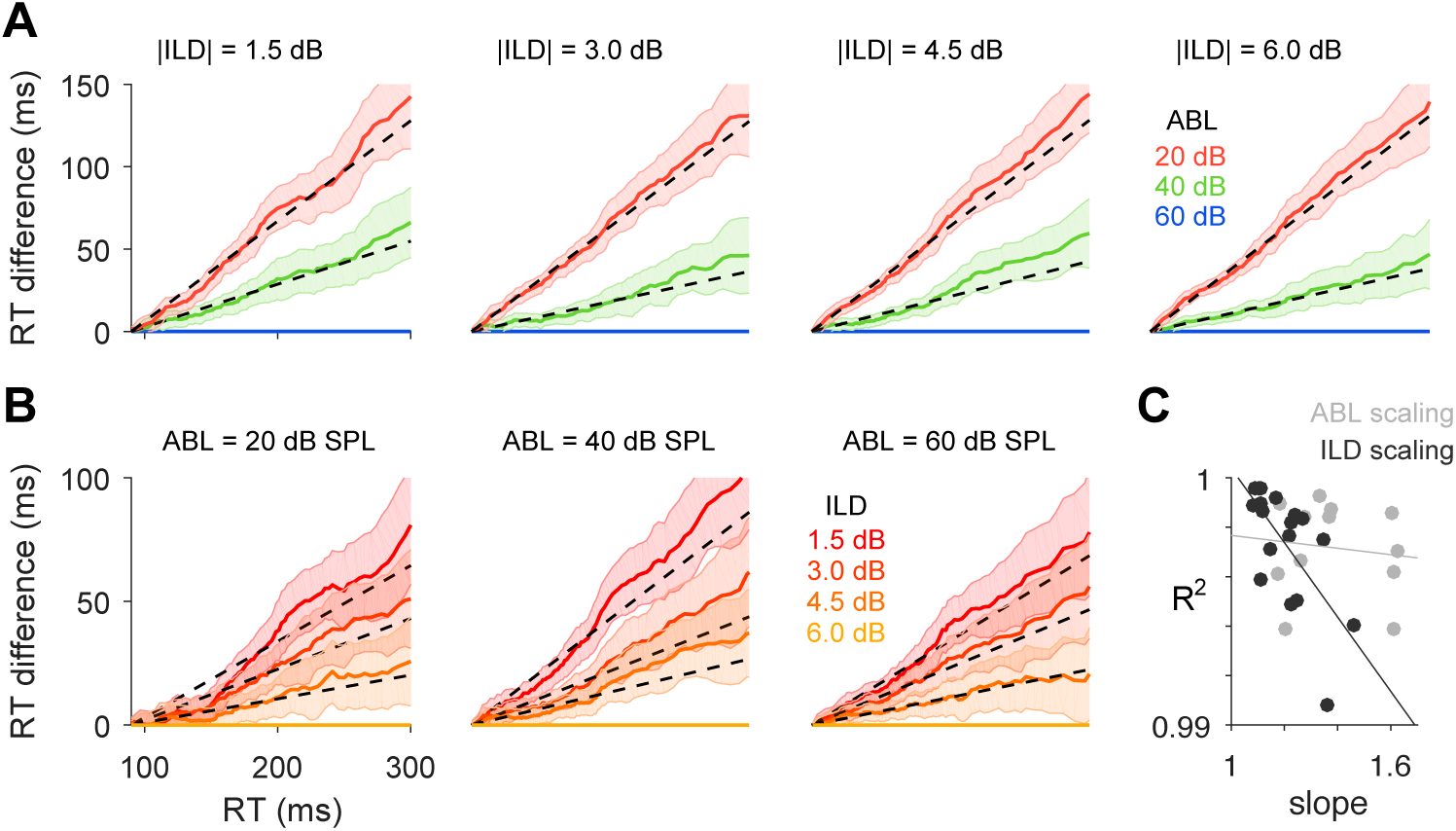
Empirical assessment of RT scale invariance. **(A)** Each plot shows, for each difficulty, *PC*_ABL_(*Q*) - *PC*_60dB_(*Q*) as a function of *PC*_60dB_(*Q*), where *PC*_ABL_(*Q*) is the 100*Q*^*th*^ percentile of the RT distribution for the corresponding ABL of that difficulty (see text). Only RTs > RT_min_ are considered in this analysis. Dashed line is the outcome of the linear fit. Shaded regions represent three standard deviations of the sampling distribution of the percentiles (see Methods). Note that higher percentiles have more error. Because of this, we used weighted least squares to do the linear fit. **(B)** Same as (A), but testing the scale invariance of the reaction time distribution (RTD) induced by changes in difficulty for each ABL separately. **(C)** R^2^ of the weighted least squares fit vs slope of the fit for the effect of changes on ABL (gray) or ILD (black) on the RTD. Lines show linear fits.

Both changes in ABL and changes in ILD are very well approximated by a uniform scaling of the RT distribution (Fig. S4). However, there is an important difference. First, the values of the slopes of the fit are closer to unity for ILD-than for ABL-induced scaling (Fig. S4C, x-coordinate of the dots), because changes in difficulty have an overall smaller effect on RT than changes in sound level. Obviously, the RTDs will look similar to each other if the variable of interest does not have a strong impact on RT. In order to understand the contribution of this effect to the quality of the linear fit, we plotted the R^2^ of each fit versus its slope. The results show clearly that difficulty and sound level have a different effect on the RTD. For changes in difficulty, the RTDs become more different in shape the larger the ILD-induced change in RT, as evident by statistically significant negative slope of a linear fit of R^2^ versus slope (*p* = 0.003, Permutation test), showing that R^2^s in this case are only large because the speed accuracy trade-off in the task is weak. In contrast, for ABL-induced changes, slopes reach higher values, but nevertheless are unrelated to the quality of the fit (slope of the fit in Fig. S4C not significantly different from zero; *p* = 0.7, Permutation test). This is in agreement with the predictions of (5) and our model, in which ABL-induced scale invariance is exact, whereas difficulty-induced scale invariance is approximate and degrades with the difference in difficulty between the two RTDs. For a thorough treatment of the nature and form of the changes in the RTD induced by difficulty, see below (Fig S6).

### 3 Mathematical Analyses

#### 3.1 Mechanistic implications of temporal rescaling by stimulus intensity

In this section we explore what can be inferred about the mechanisms underlying the discrimination process from the fact that the overall stimulus intensity exclusively changes the units of time of the discrimination problem. Without loss of generality we assume that the task is to discriminate between the intensity of two simultaneously presented stationary physical signals *s*_1_ and *s*_2_. For the moment, we assume that *s*_1_ and *s*_2_ are scalars even if the stimulus can be stochastic (e.g., in our case *s*_1,2_ represent the RMS pressure of the sound at the right and left ears). Below, we consider the implications of the stochastic nature of the stimulus.

Motivated by our findings pointing towards bounded evidence accumulation (Figs. 2-3), we consider the class of models in which the decision variable (DV) can be represented as a Continuous Markov Process (CMPs) (6), and choices are triggered when the DV hits either of two bounds for the first time (this is the decision-time). In a CMP, increments in the DV across consecutive infinitesimal time intervals are statistically independent and ‘small’ (vanishingly small as the time interval goes to zero). This model class is very flexible, allowing the temporal evolution of both the mean and the variance of the DV to depend on the instantaneous value of the DV itself and of the evidence, and also explicitly on time. We also allow for time-dependent decision bounds, which appear in normative accounts of perceptual decision making (7). Most models of decision making can be construed as a CMP (7–13).

In a CMP, the differential increment in the DV is Gaussian and thus the process can be completely specified by its mean and variance, called the drift and diffusion coefficients (6). The temporal evolution of the DV *x*(*t*) is given by the stochastic differential equation

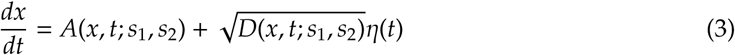

where η(*t*) is a white noise process (i.e., ⟨η(*t*)η(*t*′) ⟩ = δ(*t* - *t*′)), and *A*(*x, t*; *s*_1_, *s*_2_)*dt* and *D*(*x, t*; *s*_1_, *s*_2_)*dt* are the mean and variance of the Gaussian increment of *x* between *t* + *dt* and *t*. We make the reasonable assumption that the joint dependence of *A* and *D* on the evidence on the one hand, and on (*x, t*) on the other, arise from two separate statistically independent signals (each with white-noise variability). This just entails the assumption that the evidence *e*(*t*) and the DV *x*(*t*) exist and are well defined on their own, and that the DV can only be a function of the evidence in the past. In order to remain within the CMP framework, these two white noise processes are added, implying that the drift and diffusion coefficients are given by

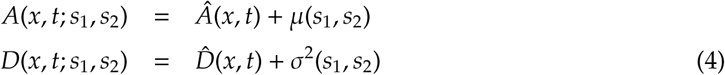

The quantities μ(*s*_1_, *s*_2_) and σ^2^(*s*_1_, *s*_2_) are the mean and variance of the evidence *e*(*t*)

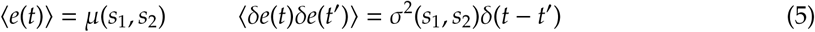

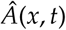Â (x, t) describes a (possibly non-linear and time-dependent) feedback (positive or negative) term of the DV on itself, and 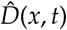 describes a possible contribution of the instantaneous state of the process and of time to the overall noise integrated by the DV.

We posit that the subject commits to one of the two available choices when the DV *x*(*t*) hits either of two bounds (or thresholds) θ_1_(*t*) and θ_2_(*t*) for the first time. Since our rats do not display significant biases, we consider unbiased discriminations in which the initial condition of the process is *x*(*t* = 0) = 0 and where the two thresholds are symmetric, i.e., θ_2_(*t*) = -θ_1_(*t*) ≡ -θ(*t*).

The analysis simplifies if one maps this problem onto one with a constant threshold, which can be done by changing variables so that the new DV 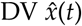 measures the fractional distance between *x*(*t*) and the instantaneous threshold, i.e.,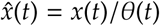. Using Ito’s formula (14) to perform the change of variables, the diffusion equation for the new DV reads

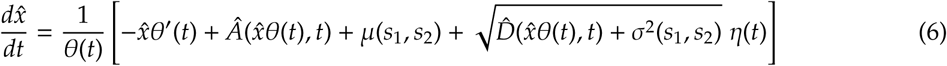

with a constant threshold of one, i.e.,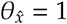.

##### 3.1.1 Problem Formulation

- WL can be formally stated as saying that the accuracy of the discrimination between *s*_1_ and *s*_2_ and that between 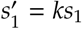 and 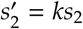 should be the same. Thus, since our hypothesis is that this is the case because the only effect of the change in stimulus intensity is to change the units of time of the discrimination process, we would like to find the most general conditions under which a discrimination between *ks*_1_ and *ks*_2_ proceeding under standard time *t* is identical to a discrimination between *s*_1_ and *s*_2_ proceeding under a uniform rescaling of time *t*′ = α*t*. We therefore seek functions θ(t), Â, 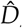, μ and σ^2^ such that the probability distribution of the differential increment of the DV in these two discriminations is identical. Since the increment is Gaussian in a CMP, this amounts to equality between their means and variances

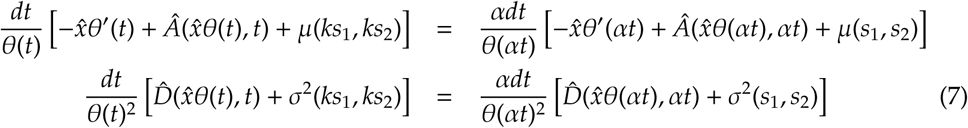

##### 3.1.2 Problem Solution

- The second equation only has a solution if 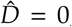, because the temporal rescaling cannot both compensate the change in the stimulus-dependent diffusion term σ^2^ and at the same time leave untouched the stimulus-independent term 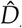.
- The first equation admits a solution where

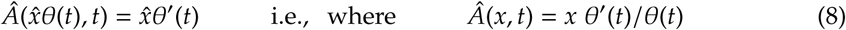
- In this case, Eqs. 7 reduce to

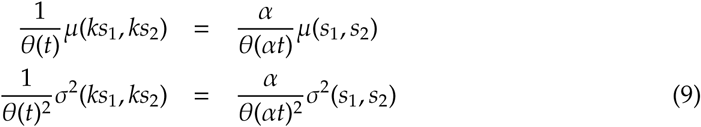 In order for these equations to be satisfied, μ and σ^2^ need to depend on the stimuli through a power-law, and θ(*t*) also needs to be a power-law function of time. This is because a power-law is the only function for which

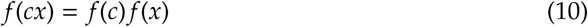 In addition, μ and σ^2^ can be any polynomial of these power-laws as long as it only contains terms of the same order. Let’s denote the order of the polynomial by *n*_*p*_, and let’s denote as λ, λ_σ_, and λ_*t*_ the exponent of the power-laws for μ and σ^2^ and θ(*t*) respectively. Then, the previous equations simplify to

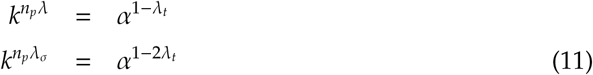 In order for both of these equations to be satisfied simultaneously, it needs to be the case that

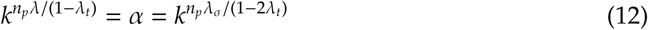

which implies that

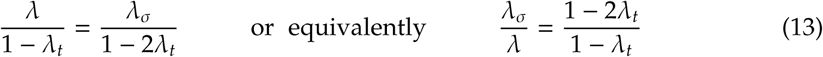 The solutions of this equation with λ_*t*_ equal or different from zero are qualitatively different.
- If λ_*t*_ is different from zero, then it has to be negative, otherwise θ(*t* = 0) = 0. This solution thus implies a decaying decision bound. From Eq. 8, the drift coefficient needs to be

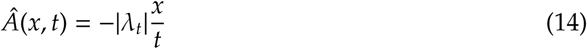

which can be interpreted as a temporally decaying standard linear leak term. The term λ_σ_/λ increases with the magnitude of λ_*t*_ and is bounded between 1 (for λ_*t*_ = 0; in this case the variance and the mean scale in the same way with the stimulus) and 2 (for λ_*t*_ = −∞; in this case the standard deviation and the mean scale in the same way with the stimulus).
- If λ_*t*_ = 0, the solution is much simpler. In this case the decision bound is constant, 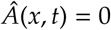 and λ_σ_ = λ. This solution corresponds to a particular form of the drift-diffusion model (DDM) which we consider below.
- Finally, symmetry considerations can further specify some properties of the mean and variance of the evidence. For unbiased decisions, swapping the two stimuli should simply reverse the agent’s choices leaving everything else the same. Since the two choices are associated to different signs of the DV, this means that the mean and the variance should be such that

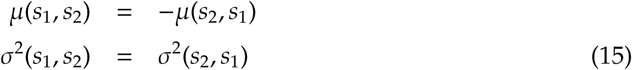

##### 3.1.3 Implementation

Here we consider whether there are simple, biologically plausible mechanisms, relying on as few hypothesis as possible, that might generate a CMP with the properties outlined in the previous section. Since our model is phenomenological, not biophysical, our goal is not mathematically prove the necessity of a particular implementation, but rather to consider the plausibility of the different solutions.

Solutions with non constant bounds, e.g., 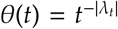 require both a specific form of the leak *A*(*x, t*) = −|λ_*t*_| *x*/*t* and a power-law relationship between variance and mean with an exponent λ_σ_/λ = (1 − 2λ_*t*_)/(1 −λ_*t*_), which is a function of the power-law exponent of the decision bound. It is not clear how such disparate physiological processes could be related, and even if this was possible, it would require their relationship to be finely tuned. Thus, although interesting mathematically, we consider this solution implausible and non-robust, and do not consider it further. From here on, we focus on the solution with a constant bound, i.e., λ_*t*_ = 0, where as we’ve shown above there is no leak term and λ_σ_ = λ.

The first step is to identify the power-law dependence of μ and σ^2^ on the stimulus as a non-linear transformation mapping stimulus intensity to firing rates. The idea is that the evidence is a function of the stochastic activity of two populations of neurons. Thus

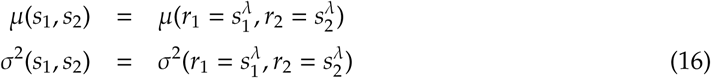

How should the activity from the two populations be combined? As mentioned above, in principle any polynomial of *r*_1_ and *r*_2_ is allowed as long as it only contains terms of the same order. Higher-order terms, however, don’t have a straight forward implementation and imply potentially complex transformations between the statistics of the activity of each population of neurons separately and the statistics of their combination. The simplest and most plausible solution is therefore a linear combination, and the anti-symmetry condition (15) implies that μ has to be proportional to the difference between the mean activities of the two populations. In the resulting model, the evidence is proportional to difference between the instantaneous activity of two populations of neurons whose mean firing rate is a power law of the stimulus intensity. Critically, the variance-to-mean relationship of the neurons needs to be linear, i.e., the Fano Factor (FF) needs to be constant.

Using Eq. 9, one obtains that, in these conditions, a multiplicative factor *k* on the stimulus intensity is identical to a rescaling of time by a factor

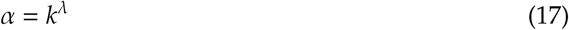

Qualitatively, *any change in the evidence that modifies its mean and its variance by the same mutiplicative factor can be absorbed by a change in the units of time*, as we show more explicitly below when describing the bottom-up construction of this model.

This implementation of the general solution to the problem is a generalization of the one proposed by Link (8) and recently also studied by Simen et al. (5), who both assumed a linear stimulus to firing rate transformation, i.e., λ = 1. A power-law transformation between stimulus intensity and firing rate has been often used in Signal Detection Theory models (see e.g., (15)). In (16), a power-law transformation in a sequential-sampling model was considered, but the authors focused on spiking statistics with non-constant FF.

#### 3.2 Temporal fluctuations in the amplitude of the stimulus

We have considered discriminations of stationary sensory stimuli. However, stationary stimuli may display temporal fluctuations in their instantaneous value. In fact, in our experiments we use broad-band noise with constant RMS, in which the instantaneous pressure level changes in time. Intuitively, temporal fluctuations in the stimulus would seem to be problematic for the evidence to meet the conditions outlined in the previous section. To see this, consider a physical signal *s*(*t*) scaled by a factor *k*, i.e., *s*′(*t*) = *ks*(*t*). This will scale by *k* the standard deviation (across time) of *s*′, not its variance, as required. Let us thus consider this situation in more detail.

For simplicity we consider the problem as discrete, so the signal is generated by sequentially drawing independent samples from a distribution on every time step. The sensory neurons have a short but finite integration time constant, which we model as a local averaging of the stimulus waveform over *n* steps. For instance, if the integration time-constant is ∼ 2 ms, for our stimulus, which has power up to 20 KHz, *n* ∼ 40. As required, sensory neurons have a constant FF, which we take for simplicity equal to 1. We thus model them as inhomogeneous Poisson processes whose rate changes every *n* steps. The time-varying firing rate is the local average of the stimulus raised to some power λ.

In these conditions, the variance in the spike counts of these neurons has two sources: one is the Poisson variability itself (which would be present even if the stimulus was completely constant in time), and another one caused by the temporal fluctuations in the stimulus, which lead to temporal fluctuations in the rate of the neurons. We are interested in knowing under which conditions this compound spike count variance is proportional to the mean, as required to obtain the shape invariance of the RTD (strictly speaking, we should refer to this as de decision-time distribution, but we will still use the acronym RTD for this quantity. In general it should be clear by the context whether we are referring to a decision-time (when talking about a mathematical model of the discrimination process) or to RT (when talking about experiments)).

Since we expect *n* to be large, the local average *s*_*n*_(*t*) of the stimulus will generally have a Gaussian distribution. We write it as *s*_*n*_(*t*) ∼ *N*(*A, A*^2^/*n*) since, as outlined above, one expects that the stimulus intensity *A* will in general scale both the mean and the standard deviation of *s*(*t*). For zero-mean sensory stimuli (such as sound pressure), *A* would represent the stimulus RMS and the sensory system would perform rectification. However, the central limit theorem guarantees that the distribution of the local average will still be Gaussian regardless of the specific form of the instantaneous distribution of *s*(*t*). For large *n*, the Gaussian distribution of *s*_*n*_(*t*) can be very well approximated by a Gamma distribution

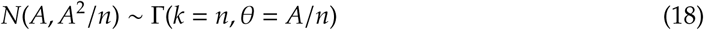

where *k* and θ are the shape and scale parameters of the Gamma distribution. We perform this approximation because the Gamma distribution is a special case of the generalized Gamma distribution, which is closed under a power transformation. It is straightforward to check that the distribution of 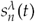 is a generalized Gamma (GG) of the form

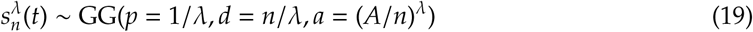

and has therefore mean and variance given by

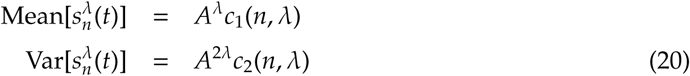

Where

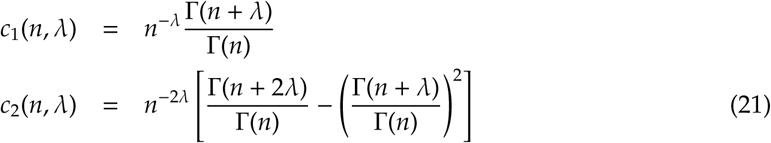

where Γ(*x*) is the Gamma function. This is the mean and variance of the time-varying firing rate of the sensory neurons.

Let us denote as ⟨*x*⟩_*r*_ and ⟨*x*⟩_*c*|*r*_ the mean over the distribution of the time varying firing rates and over the Poisson stochasticity of a neuron for a given fixed firing rate, and let us denote as ⟨ (δ*x*)^2^⟩_*r*_ and ((δ*x*)^2^)_*c*|*r*_ their respective variances. If *c* is the spike count (per unit time) of a neuron, then its overall mean and variance are given by

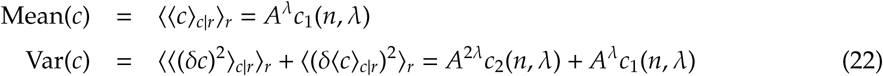

Thus

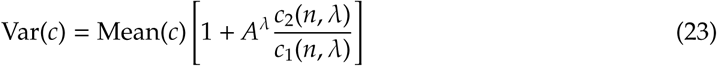

This equation clarifies the issue. Although the compound variance in fact contains a term proportional to the mean and another one proportional to the mean squared, because the coefficient *c*_2_(*n*, λ) tends to zero whereas *c*_1_(*n*, λ) tends to one as *n* becomes large (as can be verified using Stirling’s approximation), it will in general be the case that

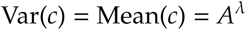

Qualitatively, as long as the temporal fluctuations in the stimulus are such that the stimulus signal-to-noise is high when it is low pass filtered by the time-constant of the sensory neurons, Poisson variability dominates and a constant FF ensues. Since these conditions are expected to hold in our experiments, it is appropriate to assume, as we have done, that the firing rate of the neurons is a power-law of the constant pressure RMS.

#### 3.3 Bottom-up construction of the model

Here we describe explicitly the implementation of the model we used to fit the experimental data.

##### 3.3.1 The evidence

The instantaneous value of the evidence *e*(*t*) is given by the difference between the instantaneous activity of two channels, corresponding to the two ears,

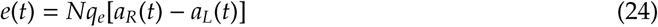

where *N* is the number of neurons in each channel, *q*_*e*_ is the quantum of evidence per spike per neuron, and *a*_*R,L*_(*t*) are the instantaneous activities (i.e., short-term spike counts) associated to firing rates *r*_*R,L*_. The firing rates are power-law functions of the RMS pressure level at each ear

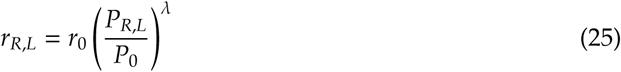

where *P*_0_ = 20μ Pa is the reference pressure level of the SPL scale. The firing rate *r*_0_ measures the overall gain of the pressure-to-rate transformation. If the power-law exponent λ is less than one, the non-linearity is compressive.

We can express the statistics of the evidence in terms of properties of our stimuli by recalling that the sound level (*SL*) at each channel is given by *SL* = 20 log_10_(*P*_RMS_/*P*_0_) and it can be expressed as *SL*_*R,L*_ = ABL ± ILD/2, so that

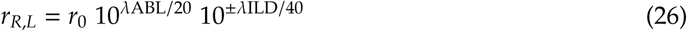

The spiking statistics of the neurons encoding the stimulus are Poisson (in principle, any process with a constant FF is appropriate. However, it is easy to show that if we assume FF ≠ 1, we cannot fit the value of FF separately from the value of other parameters. Thus, for simplicity, we assume FF = 1, i.e., Poisson statistics). We also assume that the neurons are uncorrelated. Instantaneous spiking correlations do not destroy the shape invariance of the RTD, but the model fits are already excellent without correlations, so we did not attempt to include them in our description.

Under these assumptions, the mean and auto-correlation function of the evidence across the Poisson stochasticity of the neurons can be written as

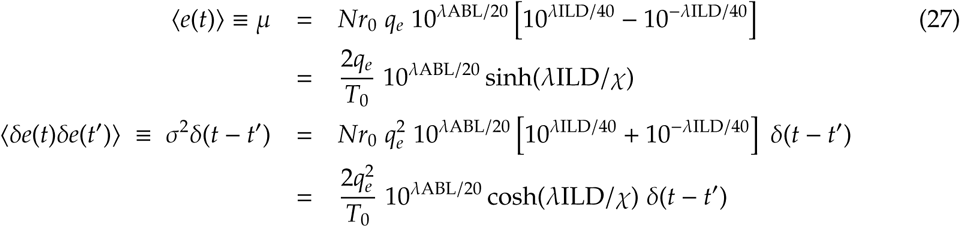

where δ*e*(*t*) ≡ *e*(*t*) -⟨*e*(*t*) ⟩ and where we have defined *T*_0_ = (*Nr*_0_)^-1^ and χ = 40/log(10) = 17.37.

##### 3.3.2 Choice

Choices are triggered at the moment the DV hits for the first time a constant decision bound ± θ (decision-time). Thus, the dynamics of the DV, whose instantaneous value we denote by *x*(*t*), are given by

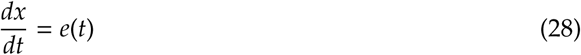

Using the diffusion approximation (17, 18) of the previous equation (which is expected to be accurate if the number of neurons in each channel is not too small and the quantum of evidence is not too big), we obtain a CMP, and re-write Eq. 28 as

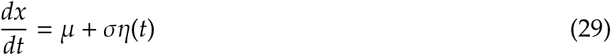

where η(*t*) is a white-noise process and μ and σ are given by Eqs. 27. We always assume that the decision variable starts at zero at *t* = 0.

In order to understand the separate contributions of ABL and ILD to the discrimination process, it is revealing to perform a change of variables. To this end, we measure the decision variable in units of the threshold, and time in units of the quantity *t*_θ_ ≡ (θ/σ)^2^ which, physically, is the mean decision-time of the process when ILD=0, and which is equal to

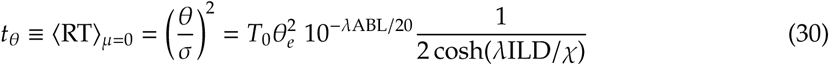
where θ_*e*_ ≡ θ/*q*_*e*_ is the threshold measured in units of the quantum of evidence. Thus, in the new dimensionless variables

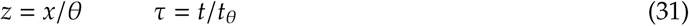

Eq. (29) reads

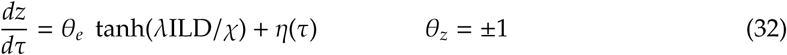

This formulation makes it clear that: (i) The dimensionless process does not depend on ABL. Changes in ABL affect exclusively the units of time of the problem and thus result only in a rigid stretching of the RTD. In general, any quantity which changes the mean and the variance of the evidence by the same multiplicative factor can always be compensated by a change in the effective unit of time of the problem. (ii) Changes in ILD affect both the strength of evidence *and* the effective units of time (see Eq. 30). Changes in the strength of evidence lead to changes in accuracy and in the shape of the RTD (see below for a quantification of the RTD depends on changes in the strength of evidence).

For a generic DDM with evidence *e*(*t*) = μ + ση(*t*), bounds at ± 1 and initial condition at zero, the probability *p*_+_ of hitting the upper bound first is a logistic function of argument 2μ/σ^2^ (13, 19). Thus, the choice log-odds for this model is given by

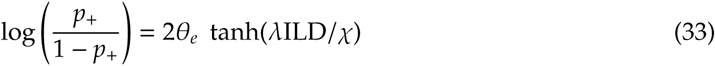

##### 3.3.3 Model behavior at psychophysical threshold

We now consider the case of decisions made close to psychophysical threshold, i.e., for ILDs small enough such that

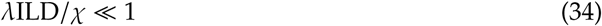

This is a very good approximation given the stimuli that we use since, considering the value of λ that we obtain from our fits, λILD/χ < 0.05 for the ILDs that we use in our task. Developing the hyperbolic functions in Eqs. 27 to first order in ILD we obtain

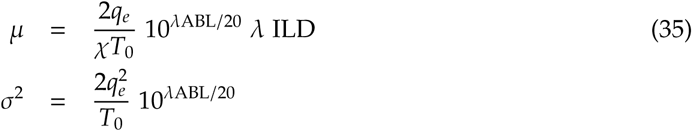

which are to the two equations in the Box of the main paper with *g*(ABL, λ) = (2*q*_*e*_/*T*_0_)10^λABL/20^. Performing the same change of variables described in the previous section, the DDM is now

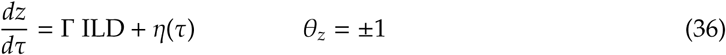

where Γ = λθ_*e*_/χ, and the effective unit of time of the problem now reads

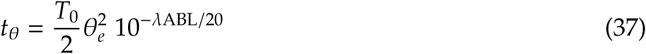

Thus, close to psychophysical threshold, it is possible to fully decouple the effect of changes in overall intensity (ABL, which only rigidly stretches the RTD) and the effect of changes in the strength of evidence (ILD, which changes accuracy and the shape of the RTD). Remarkably, the discrimination process depends only on the single parameter Γ which specifies how much evidence is associated to a given increment in ILD. A full specification of choice and decision-time depends on three parameters: θ_*e*_, λ and *T*_0_ = 1/(*Nr*_0_), as outlined in the main text and in the Box. Eqs. 36-37 correspond to Eqs. 1 and 2 of the main text. To be able to model RTs, we consider a stimulus-independent non-decision time *t*_*NDT*_ which captures sensory and motor delays unrelated to the discrimination process. We assume

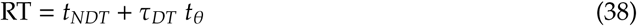

where τ_*DT*_ is the stochastic decision time associated to Eq. 36. We postulate that the non-decision time in each trial is drawn from a uniform distribution between *t*_*ND*_ - *s*_*ND*_ and *t*_*ND*_ + *s*_*ND*_ (Box). The two parameters *t*_*ND*_ and *s*_*ND*_ complete the specification of the model which we fit to the data. Close to psychophysical threshold, the choice log-odds ratio of the model is

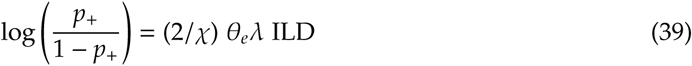

This equation makes precise the notion of a discrimination task being “sensory limited”: the choice log-odds is linear in the three critical quantities that determine accuracy: the effective decision threshold (θ_*e*_), the exponent of the compressive non-linearity (λ), which specifies how physical intensity is mapped to the magnitude of the internal representation of the stimulus, and the logratio of the physical stimulus intensities (ILD; see next section for a clarification of the relevance of the log-ratio in this expression). Our assumption that the rats operate at psychophysical threshold is validated by the accuracy of our model fits and, more directly, by the linear growth of the stimulus coefficients with ILD in our logistic regression analysis of the task (see Fig. S12B below).

### 3.4 Relationship to the standard DDM and relevance of the log-ratio

In the standard use of the DDM, only the drift coefficient μ depends on the stimuli (through an anti-symmetric function such as μ ∼ *s*_1_ - *s*_2_ or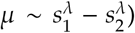, whereas the variance σ^2^ does not. However, we have shown that, in general, the model should be conceived as two-dimensional, with both the drift as well as the diffusion coefficients μ and σ^2^ being a function of the sensory stimuli *s*_1_ and *s*_2_. If the evidence is a function of the activity of neurons with constant FF, under which conditions is it appropriate to think of the model as one-dimensional? Or put slightly differently, we have been referring somewhat loosely to the ‘overall stimulus intensity’. What exactly is the overall intensity and under what conditions does it stay constant when the stimulus changes?

From the arguments we have presented, one can assume that the dependency of the mean and the variance of the evidence on the stimuli will in general be of the form

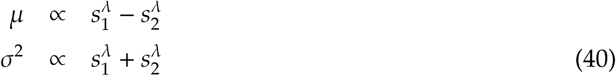

Consider the DDM in the space (*s*_1_, *s*_2_) of the two sensory stimuli to be discriminated, and a change in the stimuli from (*s*_1_, *s*_2_) to 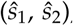, to which one can associate a change in the statistics of the evidence from (μ, σ2) to 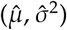. These two discrimination problems are represented in Fig. S5A. If we consider the line in stimulus space that goes from the origin to 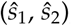, we know that moving along this line keeps the ratio of the stimulus intensities constant, which correspond to a uniform stretching of the RTD; neither choice accuracy nor the shape of the RTD changes along this line. We can also consider a curve in stimulus space that keeps the variance of the evidence constant and equal to its initial value σ^2^. Moving along this line we change the ratio of the stimulus intensities, and thus both choice accuracy and the shape of the RTD vary. We can go from any two points in stimulus space moving along these two lines, e.g., first from (*s*_1_, *s*_2_) along the constant-σ^2^ line until we cross the constant-ratio line (thick black line in Fig. S5A), and then along this straight line towards 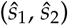 (thick gray line in Fig. S5A). Note that the RTD changes along both lines, but its shape only changes along one of them.

**Figure S5.**
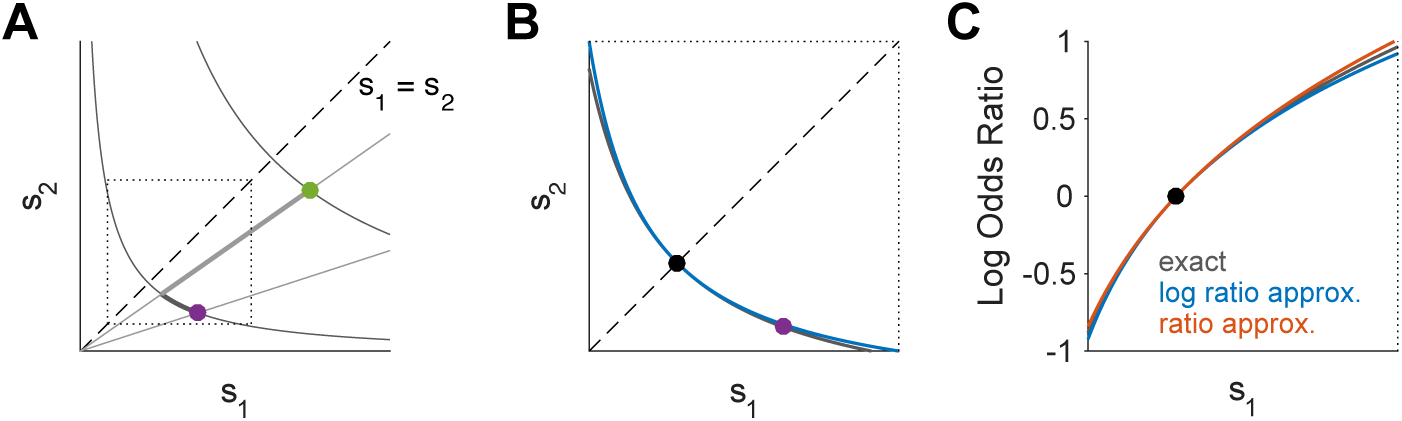
One versus two-dimensional DDM. **(A)** Two discrimination problems (filled circles) are shown in stimulus space (*s*_1_, *s*_2_). The difference between any two problems can be decomposed into a difference in the difficulty of the problem (gray curved lines, constant variance) and a difference in the effective time units of the problem (straight black lines, constant ratio). The dotted square corresponds to a maximum difference of ± 8 dB in stimulus intensity, as we sometimes use in our experiments. **(B)** Zoomed version of the dashed square region in (A), representing discrimination problems close to psychophysical threshold (i.e., *s*_1_ ∼ *s*_2_). Gray line is as in (A), and blue line is the line representing constant ABL which we use in our experiments. **(C)** Choice log-odds ratio (LOR) as one varies one stimulus (*s*_1_) while moving along the gray (gray line) and blue (blue and red) lines in (B). Black is the exact LOR. Blue is the linear approximation in log(*s*_1_/*s*_2_), equivalent to Eq. 39. Red is a linear approximation in (*s*_2_ - *s*_1_)/*s*_2_. For all panels in this figure, we used λ = 0.1.

The standard use of the DDM where changes in the stimulus only affect the drift coefficient are equivalent to moving along the constant-σ^2^ line. From the point of view of the behavior of the DDM, this is a pure change in ‘difficulty’ at fixed ‘overall intensity’. Note that this notion of intensity is related to the total number of spikes fired by the two channels on average, and this may not correspond to the subjective notion of intensity of the subject. However, from the point of view of the behavior of the model, changes in the stimulus at constant-σ^2^ are the only stimulus manipulation for which no other manipulation exists which has (i) the same performance and (ii) a different effective unit of time.

In general, keeping the variance of the evidence constant while the stimuli change requires a good understanding of how the evidence relates to the stimulus. In our simple model, it requires knowing the value of λ. However, close to psychophysical threshold, keeping the variance constant is easier, since the constant-σ^2^ line becomes perpendicular to the identity *s*_1_ = *s*_2_. Thus, any changes which keep constant any symmetric function of the stimulus are expected to also keep the variance approximately constant. In Fig. S5B we show that keeping ABL constant is a very good approximation to keeping the variance constant within the range difficulties in our experiment. Keeping *s*_1_ + *s*_2_ constant would be a worse approximation, but which would nevertheless be appropriate if *s*_1_ and *s*_2_ are sufficiently similar.

Finally, we considered whether there is anything special about the logarithmic transform. In a signal detection theory model with additive noise and a logarithmic encoding of the stimulus intensity (Fechner’s model (20)), the *d*′ of a discrimination is proportional to the log of the ratio of the stimulus intensities. We also obtain that the choice-LOR close to psychophysical threshold is proportional to ILD (Eq. 39), i.e., the log of the ratio of the RMS pressure level of the two stimuli (Fig. 1C). However, we have not explicitly invoked a logarithmic transformation. Logarithms only appear in our description because sound level is typically measured in dB. In deriving Eq. 39, which is valid near threshold, we took the exact expression for the choice-LOR (Eq. 33) and developed it to first order in ILD. However, one can also develop this expression to first order in the small quantity 1 - *s*_1_/*s*_2_. Fig. S5C shows that these two approximations are similarly accurate, at least for the range of difficulties that we have used. Thus, although the the mathematical description of our model is compact when measuring sound intensity in dB, there is nothing unique about the logarithmic transformation.

### 3.5 RT scale invariance as a function strength of evidence

Several authors have pointed out that changes in the strength of evidence of a DDM lead to changes in mean RT while preserving the coefficient of variation (CV) or the overall shape of the RTD (1–3). Importantly, scale invariance with respect to the strength of evidence (ILD for us) is a *quantitative* result (holds approximately), whereas scale invariant with respect to ABL in our model is *qualitative* (i.e., exact), as shown in the main text (see also Fig. S4). Here we quantify the accuracy of this approximation and provide an intuition for why it happens and when it fails. Across this section, we study changes in the shape of the RTD holding the variance of the evidence fixed (i.e., moving along the hyperbolic contours in Fig. S5A).

As long as accuracy is not nearly perfect (up to 90-95 % correct), the RTDs are approximately scale invariant with respect to changes in the strength of evidence (Fig. S6A-B). In Fig. S6B we used the scaling of time that minimizes the Kolmogorov-Smirnoff (KS) distance between the each RTD and the fastest one, as in the main text (similar results hold if one uses the method in Fig. S4 to calculate the temporal rescaling factor). As one increases the strength of evidence, accuracy grows (Fig. S6A-inset). If the variance of the evidence is fixed, there is a one-to-one mapping between strength of evidence and accuracy. We will use accuracy, instead of strength of evidence, to label the different RTDs since it is bounded and since it is a more natural quantity to reveal the different regimes of the RTD. The KS distance between RTDs as a function of their respective accuracies is shown in Fig. S6C. After scaling, these distances are greatly reduced (Fig. S6D), showing that RTDs corresponding to different accuracies can be approximated by scale transforms of each other (1–3).

Looking at the figure, it is evident that when the accuracy approaches one, the quality of the approximation degrades. To understand why, it is instructive to look at the analytical expression of the RTD (21)

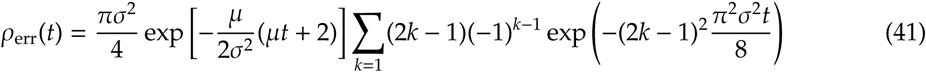

This is the probability density that the process *dx*/*dt* = μ + ση(*t*) starting from zero with bounds at ± 1, hits the lower bound at time *t* while not having hit either bound before (the unnormalized RTD for errors). We can normalize the distribution by the probability of making an error

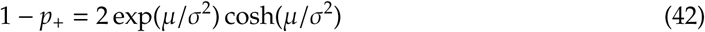

(recall that when the bounds are constant the RTD is the same for hitting either bound, i.e., correct versus error trials). Furthermore, extracting a factor exp(π^2^σ^2^*t*/8) from the infinite series, which is common to all terms, the normalized RTD can be written as

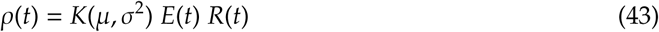

Where

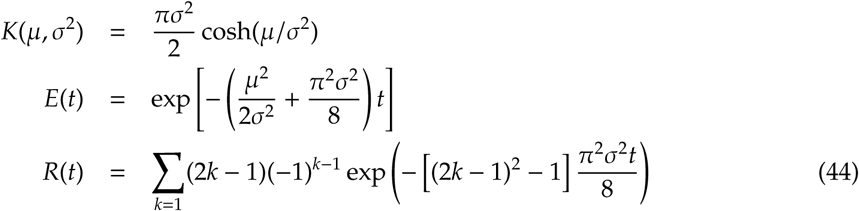

The quantity *K*(μ, σ^2^) is a normalization constant. The term *E*(*t*) is a pure exponential decay, whereas the term *R*(*t*) has the form of a ‘refractory’ period, which sets the exponential probability to zero initially (to account for the fact that the process starts away from the bound and thus, for sufficiently small times, the bound crossing probability is arbitrarily small) and quickly converges to one, setting the tail of the distribution to a perfect exponential. The exponential and refractory terms of the RTDs in Fig. S6A are shown in Fig. S6E. Importantly, only *E*(*t*) depends on the strength of evidence, in such a way that the weaker the strength of evidence, the slower the exponential decay rate, and the longer the decision-time.

**Figure S6.**
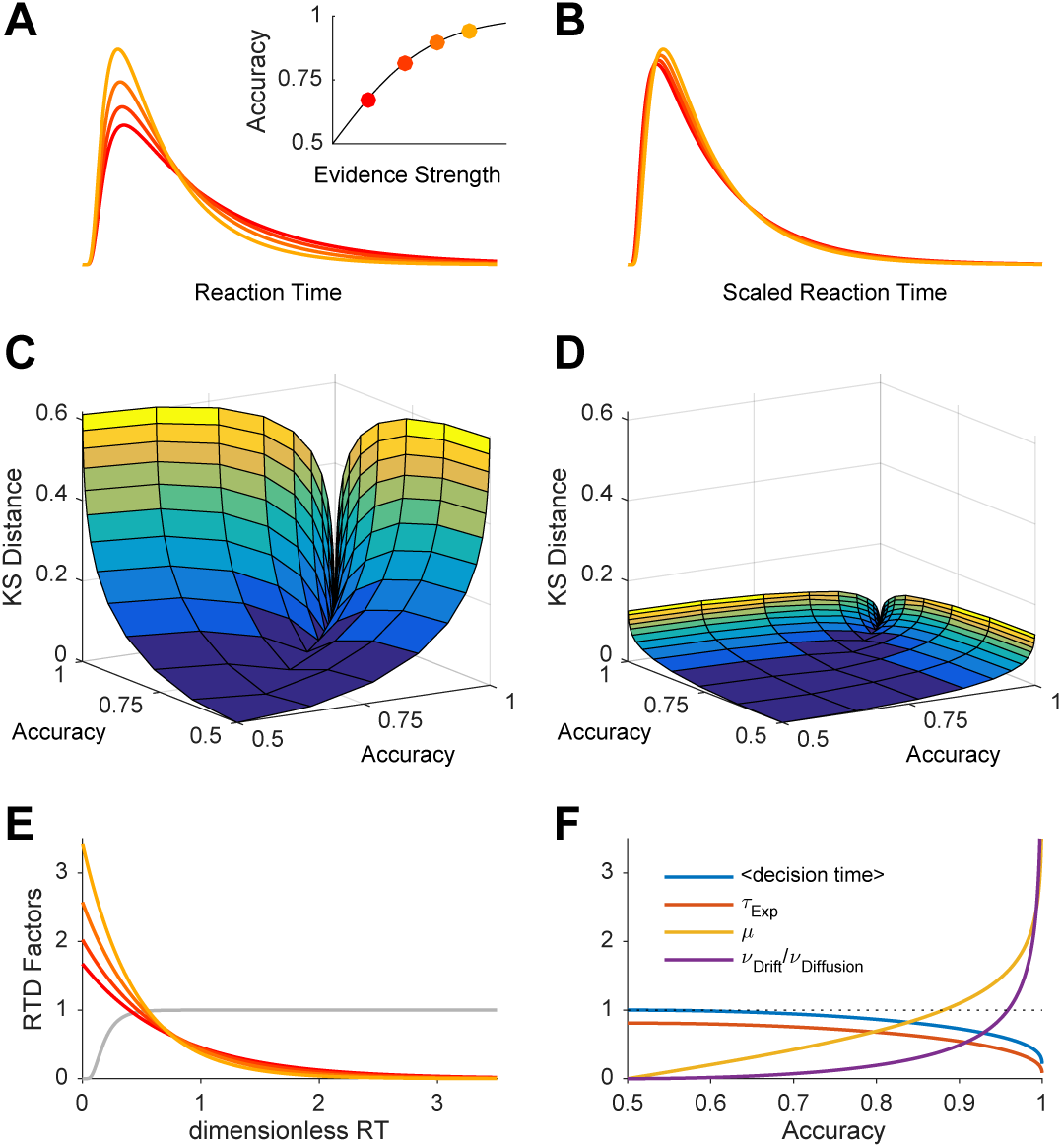
RT scale invariance with respect to changes in strength of evidence. **(A)** Four representative RTDs from the DDM. **Inset.** Accuracy as a function of strength of evidence. **(B)** Same distributions but in rescaled time. The time axis for the three slowest distributions was changed so as to maximize the overlap (minimize the Kolmogorov Smirnoff (KS) distance, see text) with the fastest one. **(C)** KS distance as a function of the accuracy associated to any pair of RT distributions. **(D)** Same after temporal rescaling. **(E)** Exponential and refractory components of the RTDs shown in (A), with the same color code. The refractory term (gray) is the same for all of them as it does not depend on the strength of evidence. The color lines show *K*(μ, σ^2^)*E*(*t*) (see Eq. 43), so that the product of the two lines is the actual RTD. **(F)** Properties of the exponential decay term *E*(*t*) as a function of accuracy. First two lines are the mean decision-time and the time constant of the exponential decay term τ_Exp_. Third line is the strength of evidence. Fourth line is the ratio ?_Drift_/?_Diffusion_ (see Eq. 45) Only for accuracies very close to unity does the strength of evidence come to dominate the shape of *E*(*t*) and thus of the RTD. The different quantities in this plot are all dimensionless and have comparable values; thus we’ve used a single y axis whose label should be inferred from the legend. For all plots in this figure we’ve assumed without loss of generality that σ^2^ = 1.

The decay rate of the *E*(*t*) term has the form

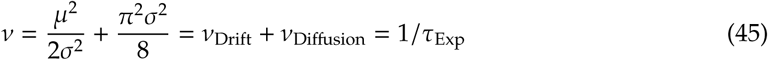

This effectively sets the rate of threshold crossings and thus the mean decision-time. The inverse τ_Exp_ of this rate, together with the mean decision-time, is shown in Fig. S6F as a function of accuracy. They depend almost identically on accuracy, although the mean decision-time is slightly longer because the small-decision-time portion of *E*(*t*) is removed by the refractory term *R*(*t*).

The decay rate *v* is the sum of two terms, one associated to the strength of evidence μ, which we call *v*_Drift_ and another one *v*_Diffusion_ associated to diffusion of the decision variable. When *v*_Drift_ « *v*_Diffusion_, the time constant τ_Exp_ and thus the mean decision-time is almost independent of the strength of evidence. To understand whether this regime takes place for any conditions of accuracy, we plotted *v*_Drift_/*v*_Diffusion_ as a function of accuracy in Fig. S6F (recall that the strength of evidence can be written in terms of accuracy as μ = (σ^2^/2) log[accuracy/(1 - accuracy)] from the equation for the psychometric function). The figure shows that the two terms only become similar at values of accuracy larger than 0.95 and that their ratio is < 0.1 for values of accuracy less than the JND. In contrast, for very high values of accuracy *v*_Drift_ grows much faster than *v*_Diffusion_ (in fact diverges).

Thus, the scale invariance of the RTD shown in Fig. S6D can be understood as deriving from the very small change in the shape of the RTD as a function of accuracy for all accuracies except those very close to one. Although perhaps intuitively one would have expected that the JND, which separates the low signal and high signal regimes in terms of accuracy, would have also separated those regimes at the level of the RT distribution, this turns out not to be the case. Instead, for almost the whole dynamic range in terms of accuracy, the RT distribution is in the low signal regime, with its associated right-skewed exponential tail characteristic of a Poisson distribution. Our empirical RTDs fully support this conclusion, as shown in Fig. S4C.

### 4 Model fitting

In this section we expand on several considerations related to the model fitting process and provide model fits for the individual rats.

#### 4.1 Constrained versus unconstrained model fitting

In the main text we have provided results (Fig. 4) using a ‘constrained’ model fitting approach. For this approach, we first fit Γ ∼ λθ_*e*_ from the psychometric function of the rats. Then, assuming that Γ is now fixed, we fit the remaining four parameters λ, *T*_0_, *t*_*ND*_ and *s*_*ND*_ using only the two extreme values of ABL = 20 and 60 dB SPL. Alternatively, one can fit all five parameters simultaneously from all the data (see Methods). Here we provide the results of the model fits for both approaches and compare them. Fig. S7A-B shows the RT quantile fits and psychometric functions for both fitting approaches. They model fits are almost identical. The estimated parameter values from the two approaches are shown in Fig. S7C and in Table S2, and are again almost identical for the constrained and unconstrained approaches, and also very similar across rats. Table S2 also shows the values of the estimated parameters using the Quantile Maximum Likelihood method (22), which are effectively identical to those obtained using the χ^2^ method.

**Figure S7.**
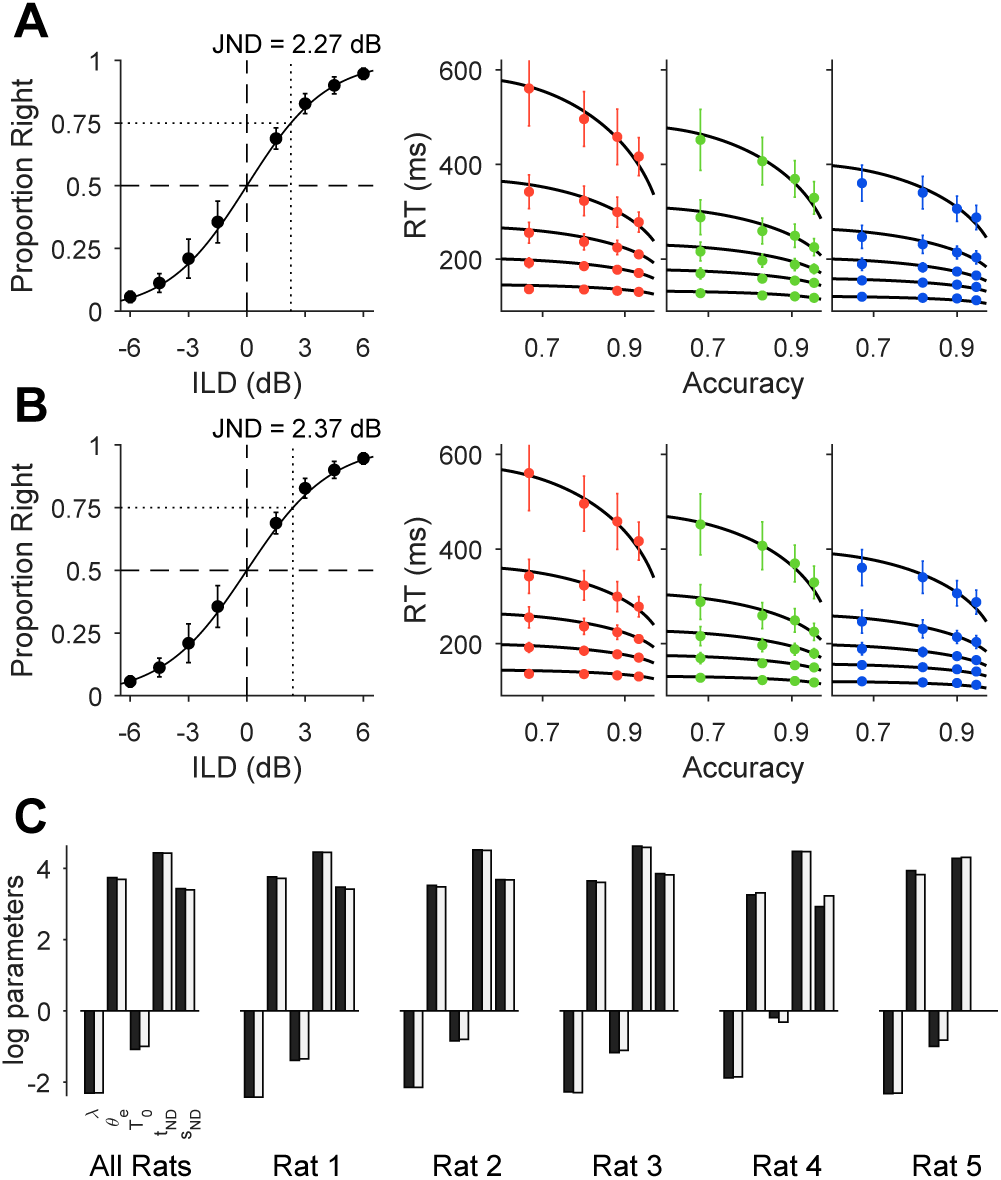
Constrained versus Unconstrained model comparison. **(A)** Constrained model. This is the same data shown in Fig. 4A-B. **(B)** Same format but for the unconstrained model. RT and accuracy were fit simultaneously and all ABL conditions were used. **(C)** Log parameter estimates for the constrained (black bars) and unconstrained (white bars) models for the pooled data and for individual rats. For rat 5, the best fit gave *s*_*ND*_ = 0 (see Table S1), so we exclude it from the plot.

We have not attempted to provide a quantitative comparison between the goodness of fit of the two approaches since the constrained model uses data from individual trials in two different fits (one for accuracy first and then another one for RT). It is more appropriate to think of the constrained approach as a method for testing model predictions.

The fact that an unconstrained fit produces the same result as the constrained fit means that both the coupling between speed and accuracy as a function of ILD, as well as the effect of ABL on RTs, have an identical form in the model and in the data.

#### 4.2 Uncertainty on parameter estimates

The fitting of non-linear models is an inherently difficult problem, as there can be large regions of parameter space which make no difference to the quality of the fit (23, 24) leading to large parameter uncertainties. We obtained an estimate of the joint probability distribution of the parameters from (unconstrained) model fits of bootstrap re-samples of our dataset (Fig. S8A). The results show that, indeed, there are strong correlations between the three parameters that describe the decision process. In contrast, the mean *t*_*ND*_ of the non-decision time *t*_*NDT*_ is very well determined and only very slightly anti-correlated with its spread *s*_*ND*_. Where does the correlation between λ, θ_*e*_ and *T*_0_ come from? The negative correlation between λ and θ_*e*_ is easy to explain, since accuracy only constraints their product. Fig. S8A shows that this product (or, equivalently Γ, which sets the rat’s JND) is very well specified by the data.

To gain an understanding of the relationship between *T*_0_ and λ we explored how they jointly determine the dependency of RT on ABL. Recall that we have assumed that

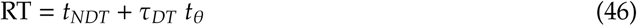

and that this temporal rescaling relationship describes our data very accurately (Fig. 4). In order to obtain a model-independent estimate of *t*_θ_, we used a procedure analogous to the one in Fig. S4, and inferred this estimate 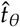 assuming Eq. 46 is true and linearly regressing the quantiles of the actual RT distributions for each value of ABL on those for the ABL-independent distribution of τ_*DT*_ specified by the dimensionless DDM in Eq. 36. We performed a single fit for all difficulties with a given ABL. Thus, we have

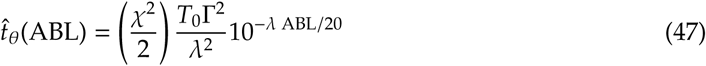

which we view as a non-linear equation with ABL as the independent variable for which we have three data points per rat. Since Γ is very well specified by the data, we assume it is constant and equal to its best-fit value and consider *T*_0_ and λ as the parameters of interest. We then use non-linear least squares to study how well λ and *T*_0_ are specified by Eq. 47. The values of 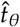 (ABL) obtained in this way together with the fit from the model in Eq. 47, and the joint distribution of the estimates of log λ and log *T*_0_ from bootstrap re-samples are shown in Fig. S8B and S8C respectively. For this model, we computed the Fisher Information matrix (FIM) at the best fit. The directions of the eigenvectors of the FIM are shown as black lines in Fig. S8C (with lengths proportional to the covariance of the parameters), and the eigenvalues are shown in the inset. The eigenvectors span several orders of magnitude, a signature of that the there is a ‘sloppy’ direction in parameter space in the model (the one along which log λ and log *T*_0_ are positively correlated) (23, 24). Inspecting the functional form of *t*_θ_(ABL), we see that the curvature of the lines in Fig. S8B is determined by λ, and that λ and *T*_0_ jointly determine the overall range of 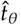 (ABL). Sloppiness arises because similar curves can be produced by small correlated changes in curvature and range. Are curvature and range actually correlated across rats? Fig. S8C suggests that the answer is yes (different rats lay along the same sloppy direction for each individual animal). Furthermore, the functional form of *t*_θ_(ABL) suggests that this correlation should exist, because the model can only produce linear functions of ABL with very small slope compared to the abscissa (because of the λ^2^ term in the denominator of *t*_θ_(ABL)). However, we don’t think the current data provides strong enough evidence of a correlation between curvature and range. On the one hand, because of the sloppy direction in parameter space, the correlation between λ and *T*_0_ is enhanced by the model fit. Also, the strong correlation depends heavily on data from a single rat (the one corresponding to the blue dots in Fig. S8C). To establish conclusively that this correlation exists we would need more rats and more values of ABL and thus this issue stands as a model prediction right now.

**Figure S8.**
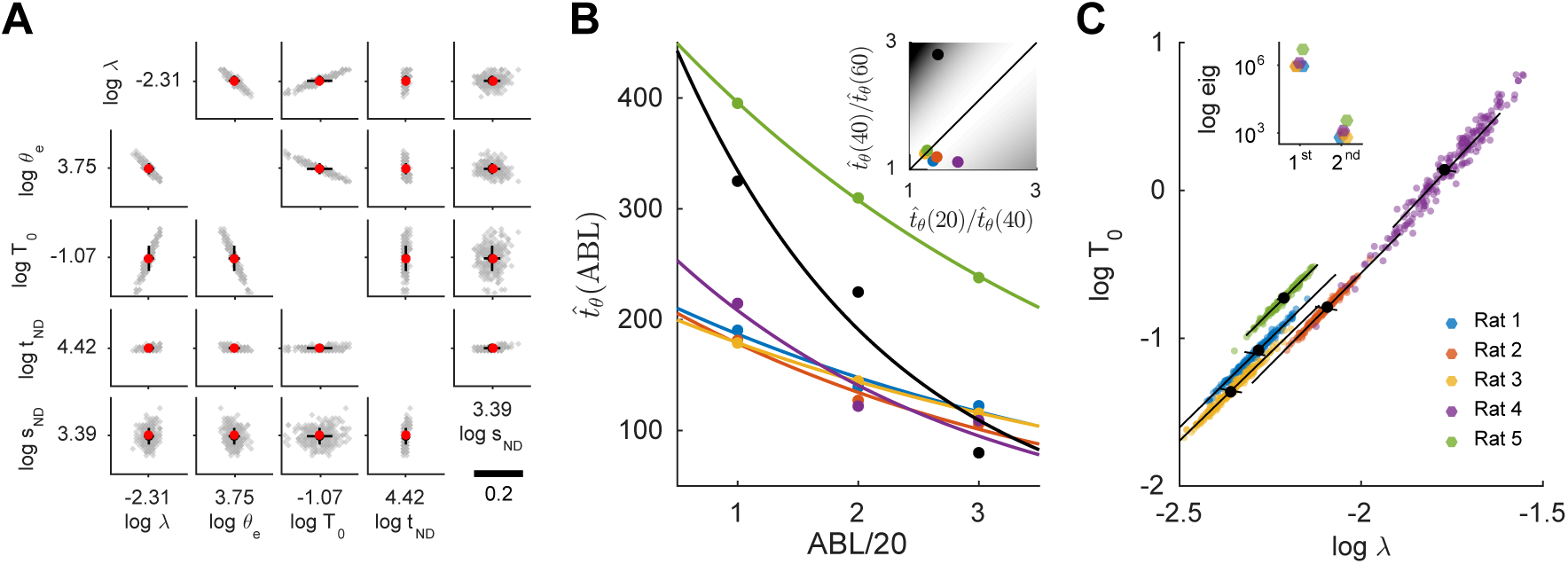
Uncertainty in parameter estimation. **(A)** Log-parameter estimates from bootstrap re-samples of the data using the unconstrained model fitting approach (see Methods). Gray: results from 200 resamples. Black: mean ± standard deviation across 1000 re-samples. Red dot is the fit to the actual data. **(B)** Empirical time-scale factor 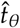 (ABL) for each of the three ABL conditions (see text). Each color is for an individual rat. Black dots are artificial data to show that only certain temporal scaling factors can be accommodated by the model. **Inset.** Ratio of temporal scaling factors for our two (equal) ABL increments plotted against each other for each rat. Model predicts they should be the same. Background color is the error in the fit (darker is more error). **(C)** Estimates of log *T*_0_ versus log λ from Eq. 47. Colored points are estimates from bootstrap re-samples. For each color, black full circle is the mean across re-samples. Black lines represent the directions of the eigenvectors of the Fisher Information Matrix (FIM) evaluated at the best fit, with lengths proportional to the covariance of the parameters (they are not orthogonal because the x-and y-axis are stretched. **Inset.** The two eigenvalues of the FIM for each rat.

Finally, we not that, despite sloppiness, the model is still quite sensitive values of 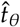 (ABL). For instance, from Eq. 47 it is straightforward to check that the following equality should hold

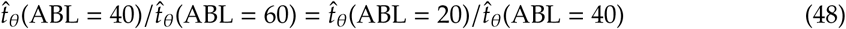

Values of 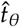 (ABL) perfectly within the range of what we observed, but which strongly violate this equality (black circles in Fig. S8B), give poor model fits (Fig. S8B inset).

#### 4.3 Model fits for individual rats

Here we show the results of the model fits for each of the five rats individually.

**Fig S9.**
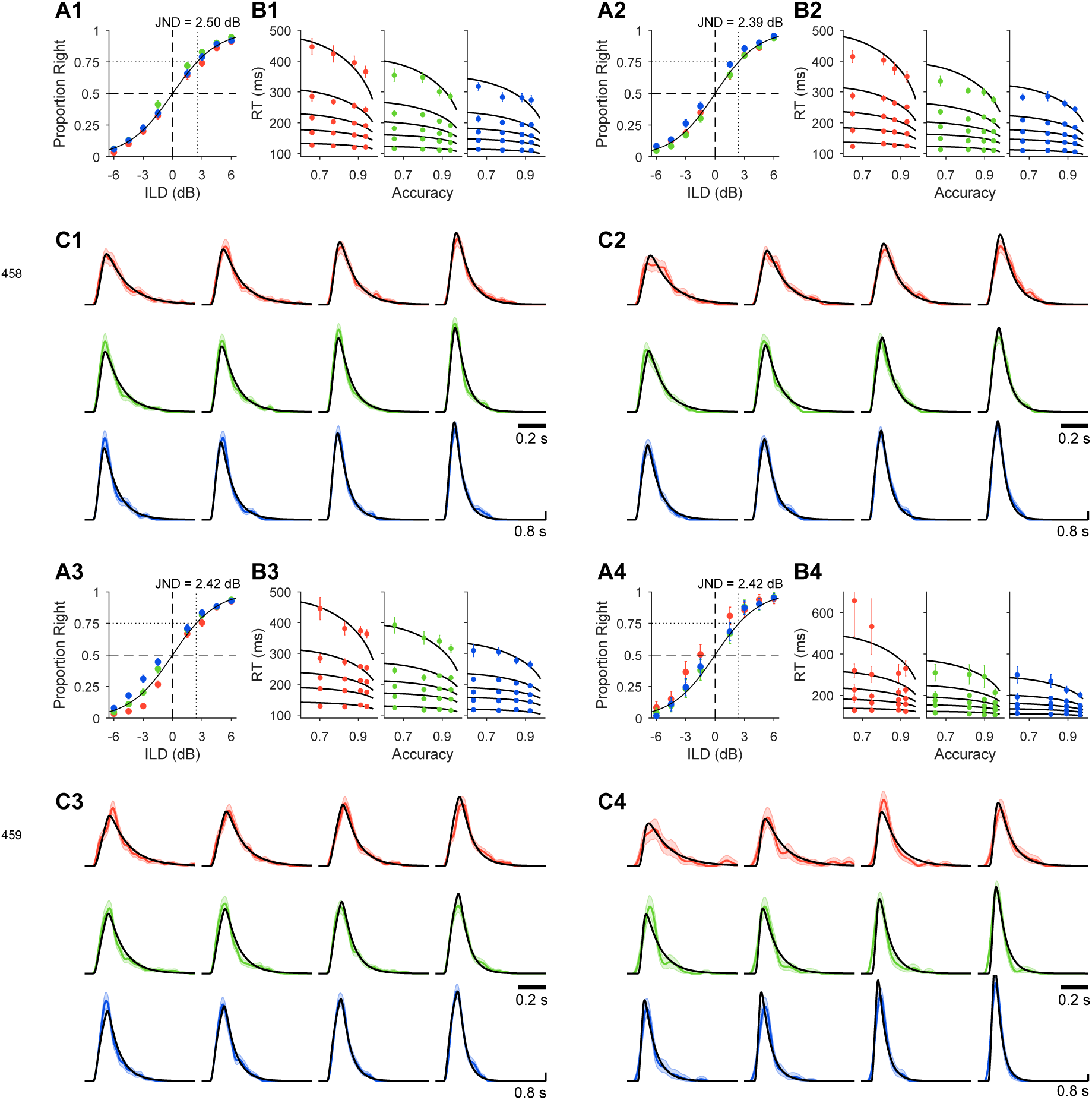
Model fits for individual rats. Data for Rats 1-4. Continues in the next page.

**Fig S9.**
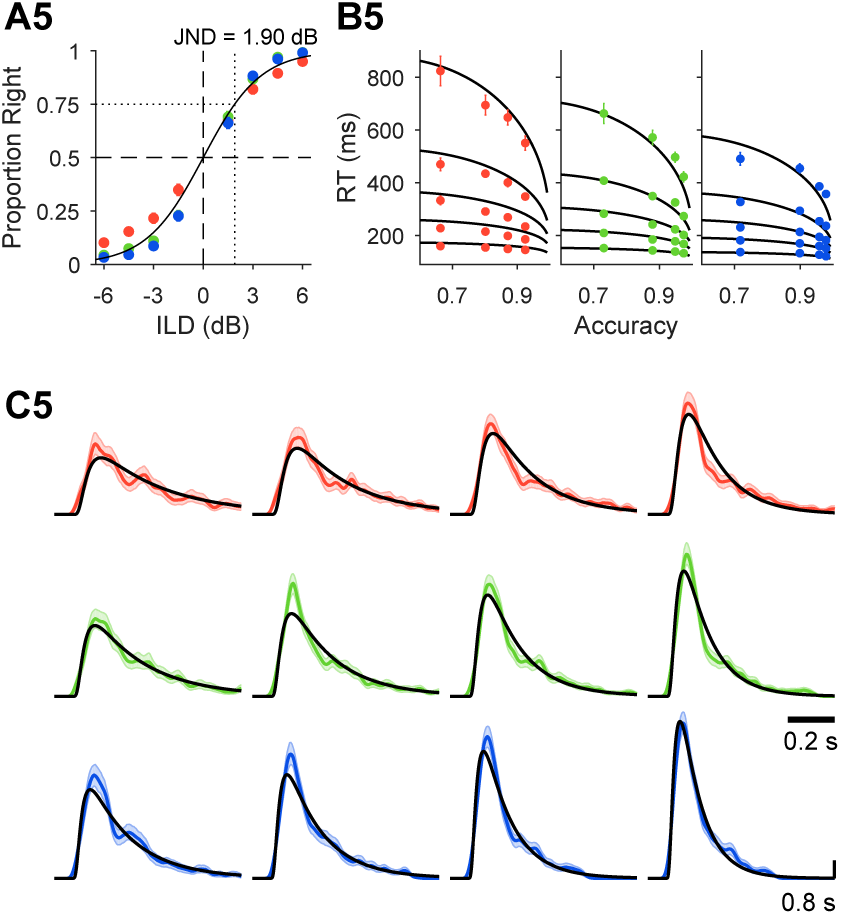
Model fits for individual rats. Same format as in Fig. 6 of the main text. **(A1-5)** Psychometric functions. Dots are choose-right probabilities separately for each ABL (same color code as Figs. 2-3 main text). A single psychometric function has been fit to pooled data across ABLs. **(B1-5)** The five quantiles for the RTDs for the three ABLs and their model fit. Error bars are standard deviations across bootstrap resamples (*N*_*r*_ = 1000). **(C1-5)** RTDs (kernel density estimates, see Methods) for each rat and their corresponding model fit across all twelve conditions. Results are more noisy than for the pooled data but the model can still fit the behavior of each single rat accurately. For all plots, we show the range 0-800 ms.

### 5 Task validation and behavioral manipulations

#### 5.1 Sources of uncertainty in a sensory psychophysics task

In order to interpret correctly the width of the psychometric function, it is important to understand what are the sources of uncertainty that contribute to its value. For instance, we are using choices from our rats to fit parameters that specify how evidence accumulation of noisy sensory input determines accuracy, which implicitly assumes that it is factors related to the specific sensory discrimination process that we model that set the accuracy of the rats’ choices. Fig. S10 shows schematically factors that may contribute to the slope of the psychometric function in a signal detection theory setting. Here, the goal may be to measure ‘sensory’ uncertainty, i.e., the trial to trial variability in the decision variable. This would be associated to the black psychometric function in the figure. If the task is not fully understood by the subject or the decision boundary has to be retrieved from memory, this may lead to what we call ‘decision uncertainty’, i.e., “I know what’s out there but I don’t know what I should do to get reward”. If decision uncertainty is present, this will decrease the slope of the psychometric function (blue psychometric). Furthermore, the actions that the subject needs to perform in the task may be under the control of a variety of mechanisms, some of them independent of the stimulus in the current trial. For instance, subjects may tend to repeat an action if it led to reinforcement in the past regardless of the current stimulus. The extent to which these alternative mechanisms will take control of the current action will be modulated by a number of different factors. One series of factors may have to do with trial history. Additionally, some discriminations may be particularly effortful, which could bias control away from the task. Finally, general uncertainty about the right course of action may also recruit alternative strategies. All of these factors would further decrease the slope of the psychometric fuction (dark blue psychometric). In our task, we have systematically considered these factors (Figs. 5, S13) and found that they do not contribute to the discrimination accuracy of the rats.

**Figure S10.**
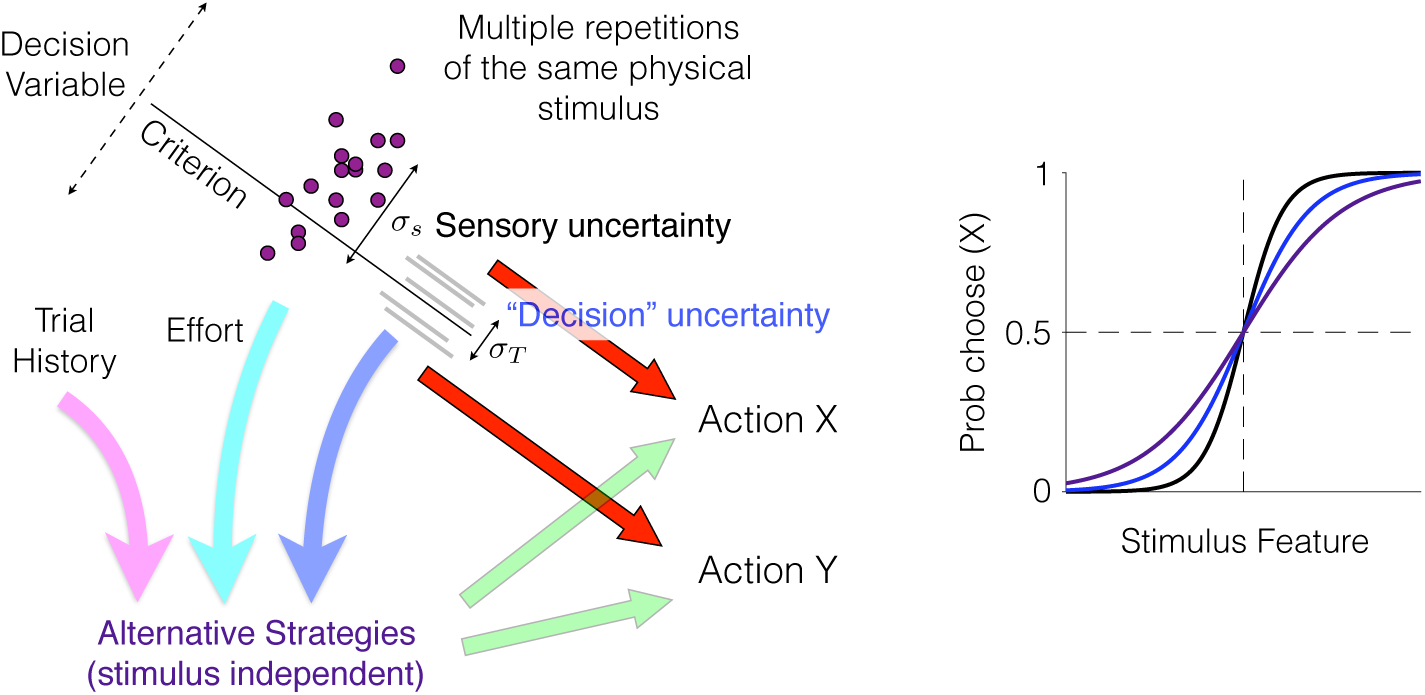
Different sources of uncertainty contribute to the slope of the psychometric function. Left. Schematic description of how task contingencies in addition to other behavioral strategies might all lead to the same two actions studied in a binary choice experiment. **Right.** Each additional source of uncertainty or stimulus-independent behavioral strategy broadens the psychometric function. Black, sensory uncertainty. Blue, idem plus decision uncertainty. Dark-blue, idem plus stimulus-independent behavioral strategies.

### 5.2 Stimulus generalization and role of external noise

In order to test that rats understood the task contingencies and responded exclusively based on the ILD of the stimulus, we first compared performance across block transitions corresponding to changes in ABL (Fig. S11A). Sensitivity (*d*′) in the first and last 24 trials of each block was not statistically different (*p* > 0.5, Fisher’s exact test). Next, we tested whether rats trained with the standard broad band noise stimulus could discriminate the ILD of pure tones (ABL = 60 dB SPL).

**Figure S11.**
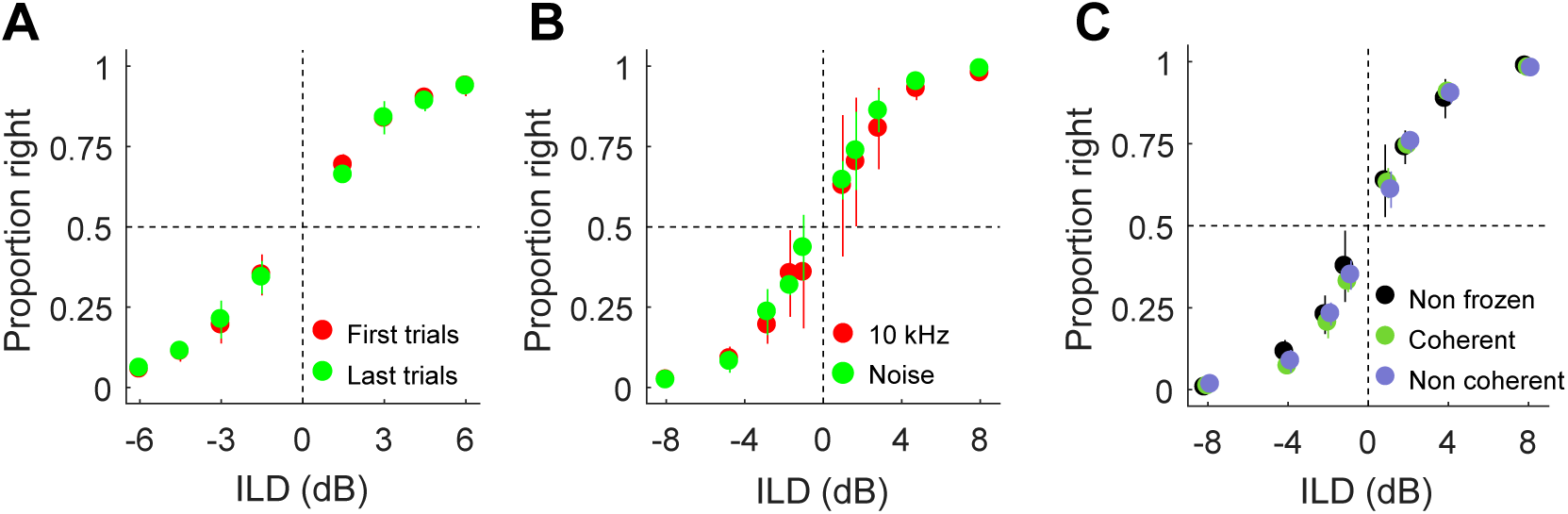
Generalization and external noise. **(A)** Choose-right probabilities for the first and last 24 trials of each block. **(B)** Same but for standard blocks versus blocks where the stimulus was a 10 KHz pure tone at 60 dB SPL. **(C)** Same comparing the standard noise stimulus and frozen noise stimuli, which could be coherent or non coherent across the two ears (see Methods).

In this case, again, sensitivity did not differ significantly across conditions (Fig. S11B; *p* > 0.5, Fisher’s exact test), demonstrating that the rats understand the task contingencies and are able to generalize, extracting and reporting the relevant feature (ILD) of sounds of different spectral content.

We also tested the implicit model assumption that discrimination accuracy is limited by vari-ability in the spiking of sensory neurons. Since the broad band noise stimulus is stochastic in nature, this stochasticity could in principle contribute to the measured value of the JND (25). We tested this hypothesis by applying the same ‘frozen noise’ stimulus (appropriately scaled in amplitude) to each ear. Sensitivity for frozen and non-frozen noise stimuli was not significantly different (Fig. S11C; *p* > 0.1, Fisher’s exact test), confirming that, at least in our experimental conditions, rats are indeed using the constant RMS pressure-level of the stimulus to perform the task, and that stochastic temporal fluctuations in the stimulus *per se* do not limit accuracy.

#### 5.3 Trial history effects

In order to reveal the extent to which variables different from the sensory stimulus in the current trial had control over the rat’s responses (26, 27), we quantified the predictive power of the history of stimuli, responses, responses after correct outcomes and responses after errors on choices in the current trial using logistic regression (see Methods). The fraction of variance of the linear component of the logistic regression model captured by the stimulus and each of the four types of history effects we considered, shows that, for four out of the five rats, trial history only marginally affected the responses of the animals in the current trial (Fig. S12A). To formally evaluate whether trial history carries predictive power, we compared prediction accuracy using the area under the curve (AUC; nested cross-validation; see Methods) between the actual data and surrogate data-sets where the trial history is randomly shuffled. Only for one rat (rat 1) there is a small but significant effect of trial history (Fig. S12B, no overlap between the 95% CI of the AUC). For this rat, the correct-response history predictor stands out and captures ∼ 8% of the variance (Fig. S12A, inset). Inspecting the four history kernels in Fig. S12D-G we can see that this effect has the form of a win-stay strategy (the correct history kernel for this animal is the outlier reaching ∼ 0.45 at trial one into the past). Anecdotally, this rat was ‘lazy’, i.e., it had a tendency to keep the back of her body close to the lateral port it had just consumed reward from and thus oriented its head towards the central port at an angle in the next trial (data not shown). It thus incurred in a higher physical cost changing responses than responding to the same side, and this is evident in the kernel.

As expected from the level invariance of the accuracy, the magnitude of the linear coefficients associated to the stimulus in the current trial for the different ABLs are very similar (Fig. S12C). Interestingly they are also approximately linear as a function of ILD, in accordance with the predictions of the theory for discriminations that take place at psychophysical threshold.

We conclude that alternative strategies have no control over the responses of the rats in the task, and that their choices reflect exclusively their perception of the ILD of the sounds in the current trial, as implicitly assumed by our model.

**Figure S12.**
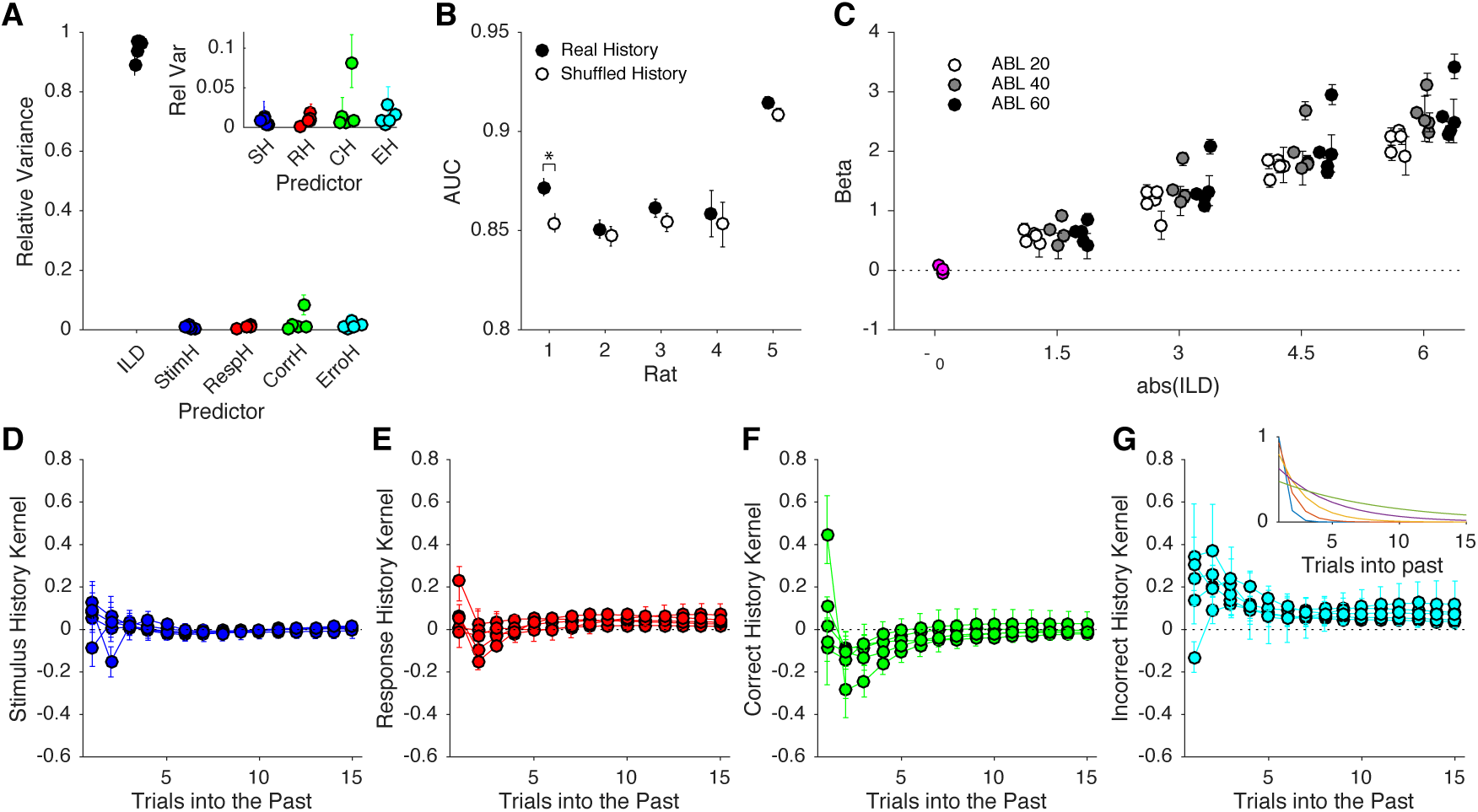
Trial history effects. **(A)** Fraction of variance of the linear component of the logistic regression model captured by the stimulus and the four types of history effects we considered. Each dot is a single rat. **Inset.** Zoom in on the fraction of variance associated to the history effect predictors. **(B)** Area under the curve (AUC) for each rat for the actual data and for surrogate datasets where history predictors have been shuffled (see Methods). **(C)** Current trial stimulus predictor coefficients separately for each ILD and ABL. **(C-F)** Kernels for each of the four types of history predictors. Notice the difference in the scale of the y-axis between this panel and panel (C). Dots connected by lines indicate the kernel for each single rat. **Inset** in (G) shows the basis of decaying exponentials used to express all kernels. Across the figure, error bars are bootstrap 95% CI (see Methods).

#### 5.4 Further manipulations of motivation and effect of priors on difficulty

We sought to further establish the surprising lack of effect of motivation on performance. The batch of animals used in the main text experienced sessions where some blocks had only the two hardest conditions (Fig. 5). For a new batch of animals, we tested longer manipulations of motivation and we also changed motivation bi-directionally. We assessed the effect of increased motivation in sessions in which only the first block was standard, and the rest of the session only used stimuli from the hardest conditions (which in this case were ILDs of 1 and 2 dB, below the JND). Although there was a trend towards an increase in sensitivity for the only-hard blocks, the increase did not reach significance (Fig. S13A; *p* = 0.064, Fishe’s exact test). We further tested whether experiencing only easy stimuli would decrease performance. For this, rats experienced many sessions of only easy stimuli (ILDs of 4.5, 6, 9, 15 dB). Against our expectations, lapse rates did not increase and percent correct was not significantly different (Fig. S13A; comparing difficulties of 4.5 and 6 dB with our standard dataset, *p* > 0.1, Fisher’s exact test. Since there were few errors in these conditions, assessments of sensitivity were unreliable).

In addition to testing the effect of motivation, changes in the overall difficulty of a block can be used to test the effect of priors on difficulty. Normative bayesian models of perceptual decision-making (7, 28) predict that the psychometric function should change depending on the subjects prior on the difficulty of the stimulus. We reasoned that the best chance to see this effect would be at transitions between blocks with different overall difficulties. Thus, we trained rats in a series of sessions with transitions between standard, hard-only and easy only conditions. Consistent with our previous results, we saw no statistically significant differences in sensitivity (for standard versus only-hard) or percent correct (for standard versus only-easy) at block transitions (see Table S3 for statistical comparisons).

**Figure S13.**
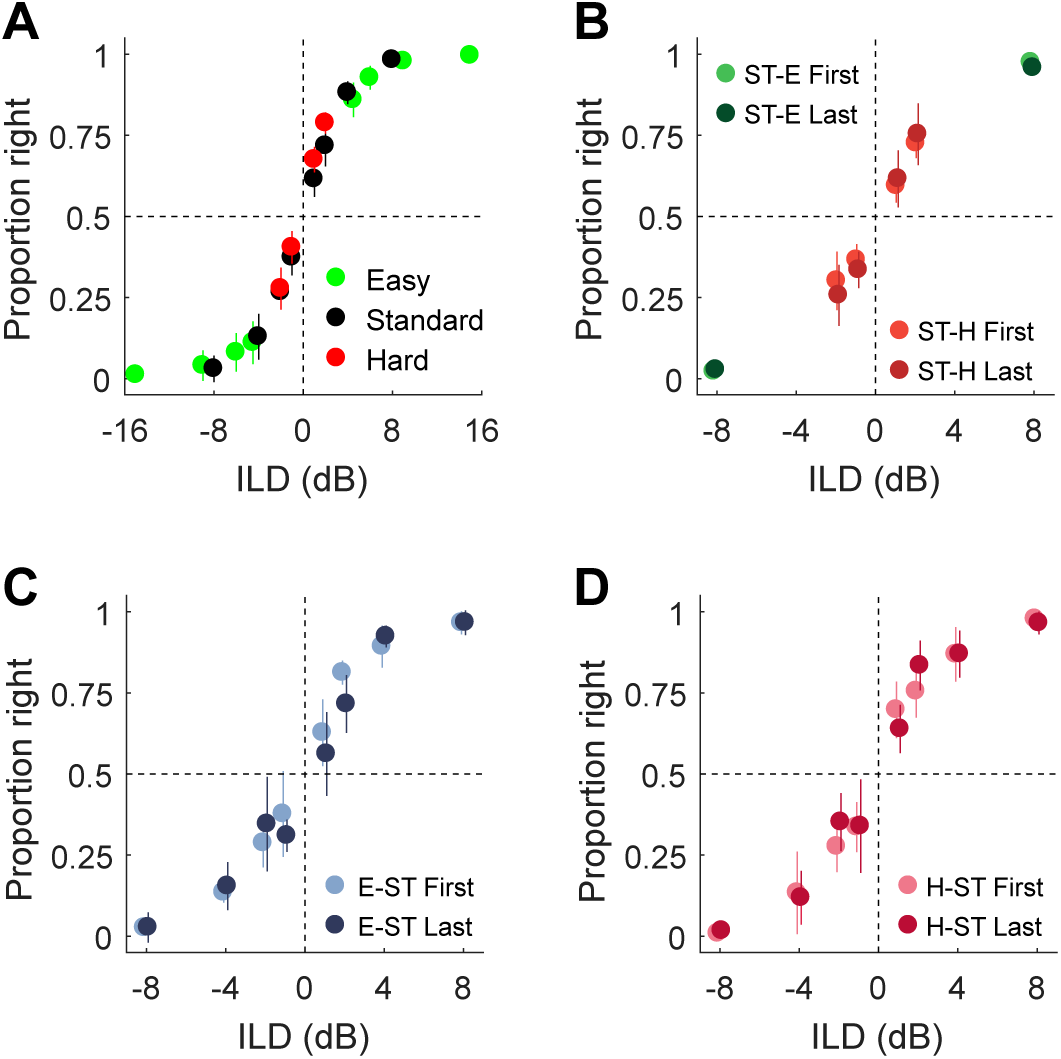
Stronger bi-directional manipulations of motivation and effect of priors on difficulty. **(A)** Choose-right probabilities for standard, only-hard or only-easy sessions (see Methods). Choose-right probabilities around changes from: **(B)** standard -ST-to easy blocks -E- and from standard to hard blocks -H-; **(C)** easy to standard blocks; and **(D)** hard to standard blocks. In all figure legends, First and Last are the first and last 24 trials of each block, respectively.

Overall we conclude that the evidence threshold for discrimination is remarkably robust against changes in motivation or priors on difficulty.

### 6 Supplementary Tables

#### 6.1 Accuracy and RT comparisons

**Table S1.**
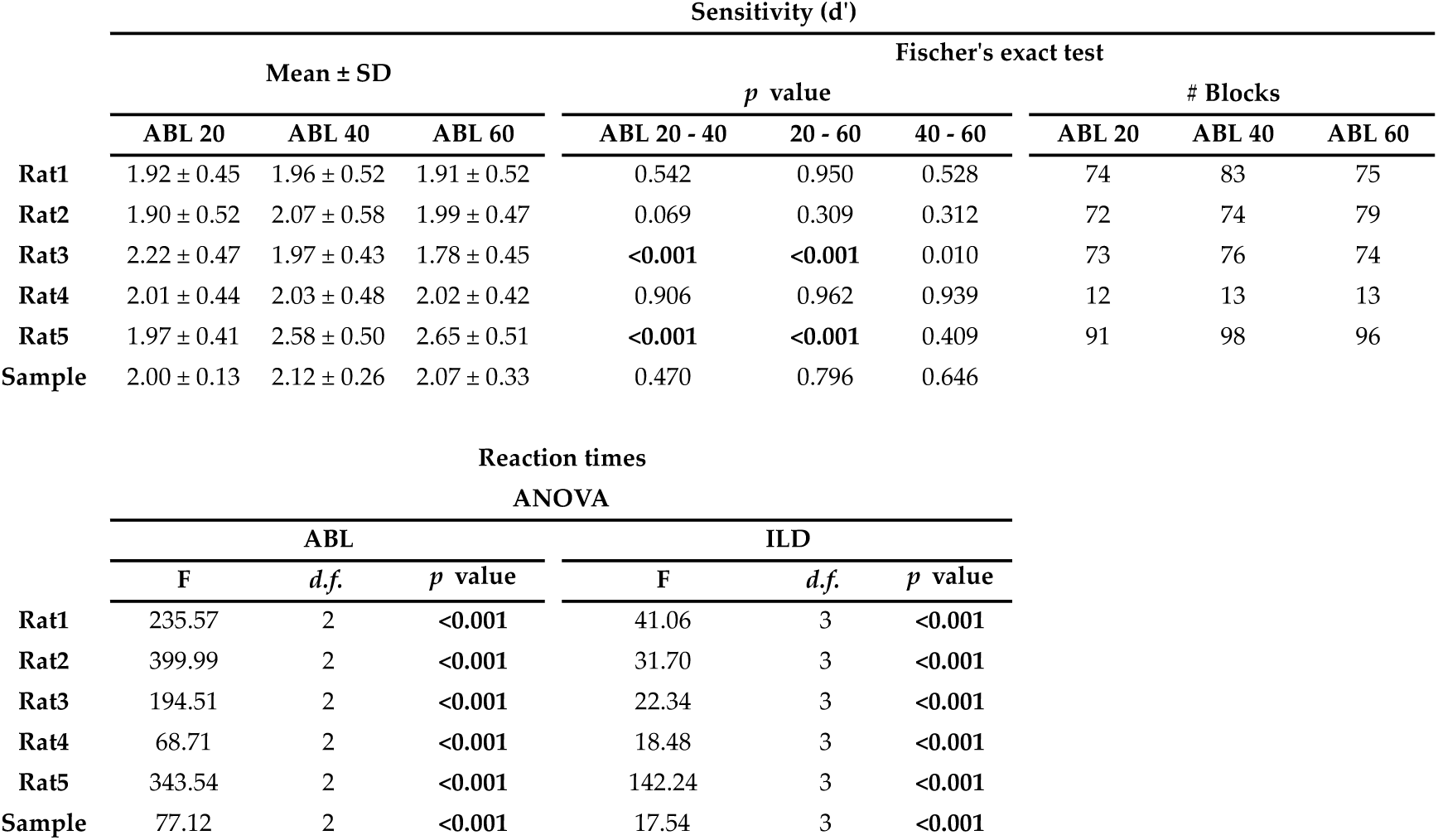
Accuracy and reaction time results. **Top.** Accuracy. For each rat, we report sensitivity (*d*’) for each ABL (first 3 columns), *p*-Value of a sensitivity comparison for each pair of ABLs testing the null hypothesis that they are the same (Fisher’s exact permutation test, Bonferroni corrected), and number of blocks in each condition (each block is 80 trials). Rat 4 lost the implant early. **Bottom.** Results for two-way ANOVA testing differences in mean reaction time across ABL (left three columns) and ILD (right three columns).

| | Sensitivity ($d'$) | | | | | | | | |
| --- | --- | --- | --- | --- | --- | --- | --- | --- | --- |
| | Mean ± SD | | | Fischer's exact test | | | | | |
| | | | | $p$ value | | | # Blocks | | |
|  | ABL 20 | ABL 40 | ABL 60 | ABL 20 - 40 | 20 - 60 | 40 - 60 | ABL 20 | ABL 40 | ABL 60 |
| Rat1 | 1.92 ± 0.45 | 1.96 ± 0.52 | 1.91 ± 0.52 | 0.542 | 0.950 | 0.528 | 74 | 83 | 75 |
| Rat2 | 1.90 ± 0.52 | 2.07 ± 0.58 | 1.99 ± 0.47 | 0.069 | 0.309 | 0.312 | 72 | 74 | 79 |
| Rat3 | 2.22 ± 0.47 | 1.97 ± 0.43 | 1.78 ± 0.45 | <0.001 | <0.001 | 0.010 | 73 | 76 | 74 |
| Rat4 | 2.01 ± 0.44 | 2.03 ± 0.48 | 2.02 ± 0.42 | 0.906 | 0.962 | 0.939 | 12 | 13 | 13 |
| Rat5 | 1.97 ± 0.41 | 2.58 ± 0.50 | 2.65 ± 0.51 | <0.001 | <0.001 | 0.409 | 91 | 98 | 96 |
| Sample | 2.00 ± 0.13 | 2.12 ± 0.26 | 2.07 ± 0.33 | 0.470 | 0.796 | 0.646 | | | |

**Table S1.** Accuracy and reaction time results.
|  | Reaction times |  |  |  |  |  |
| --- | --- | --- | --- | --- | --- | --- |
|  | ANOVA |  |  |  |  |  |
|  | ABL |  |  | ILD |  |  |
| | F | $d.f.$ | $p$ value | F | $d.f.$ | $p$ value |
| Rat1 | 235.57 | 2 | <0.001 | 41.06 | 3 | <0.001 |
| Rat2 | 399.99 | 2 | <0.001 | 31.70 | 3 | <0.001 |
| Rat3 | 194.51 | 2 | <0.001 | 22.34 | 3 | <0.001 |
| Rat4 | 68.71 | 2 | <0.001 | 18.48 | 3 | <0.001 |
| Rat5 | 343.54 | 2 | <0.001 | 142.24 | 3 | <0.001 |
| Sample | 77.12 | 2 | <0.001 | 17.54 | 3 | <0.001 |

#### 6.2 Parameter estimates from model fits

**Table S2.**
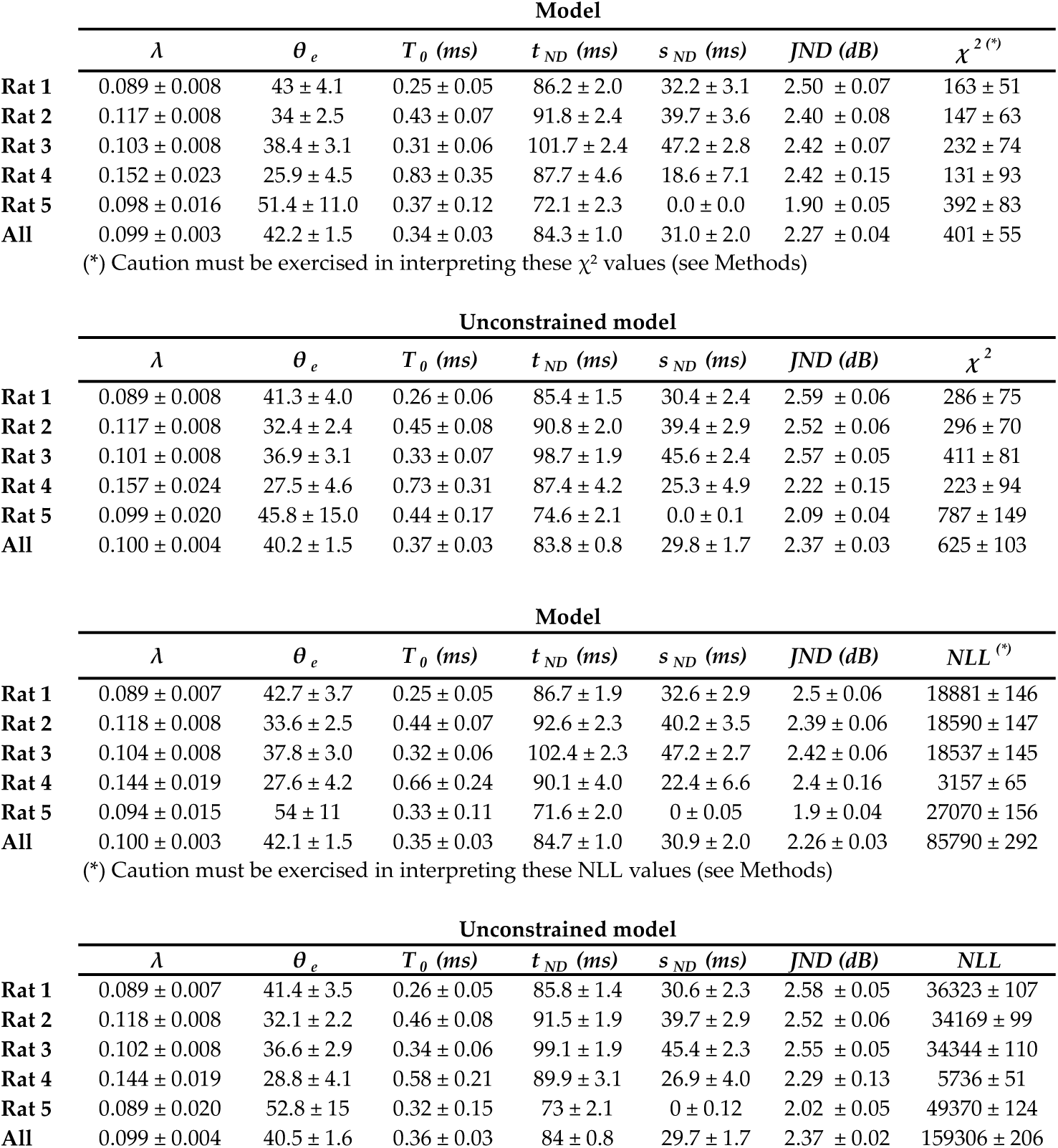
The top two tables list the model parameter estimates for the constrained and unconstrained models using the χ^2^ method for all rats individually and for the pooled data across rats. The log of these estimates are represented graphically in Fig. S7C. For the unconstrained model, the χ^2^ values are statistically meaningful. The bottom two tables show the same thing but using the quantile maximum likelihood method (see Methods), instead of the χ^2^ method. The results are essentially identical regardless of which method is used.

#### 6.3 Behavioral manipulations

**Table S3.**
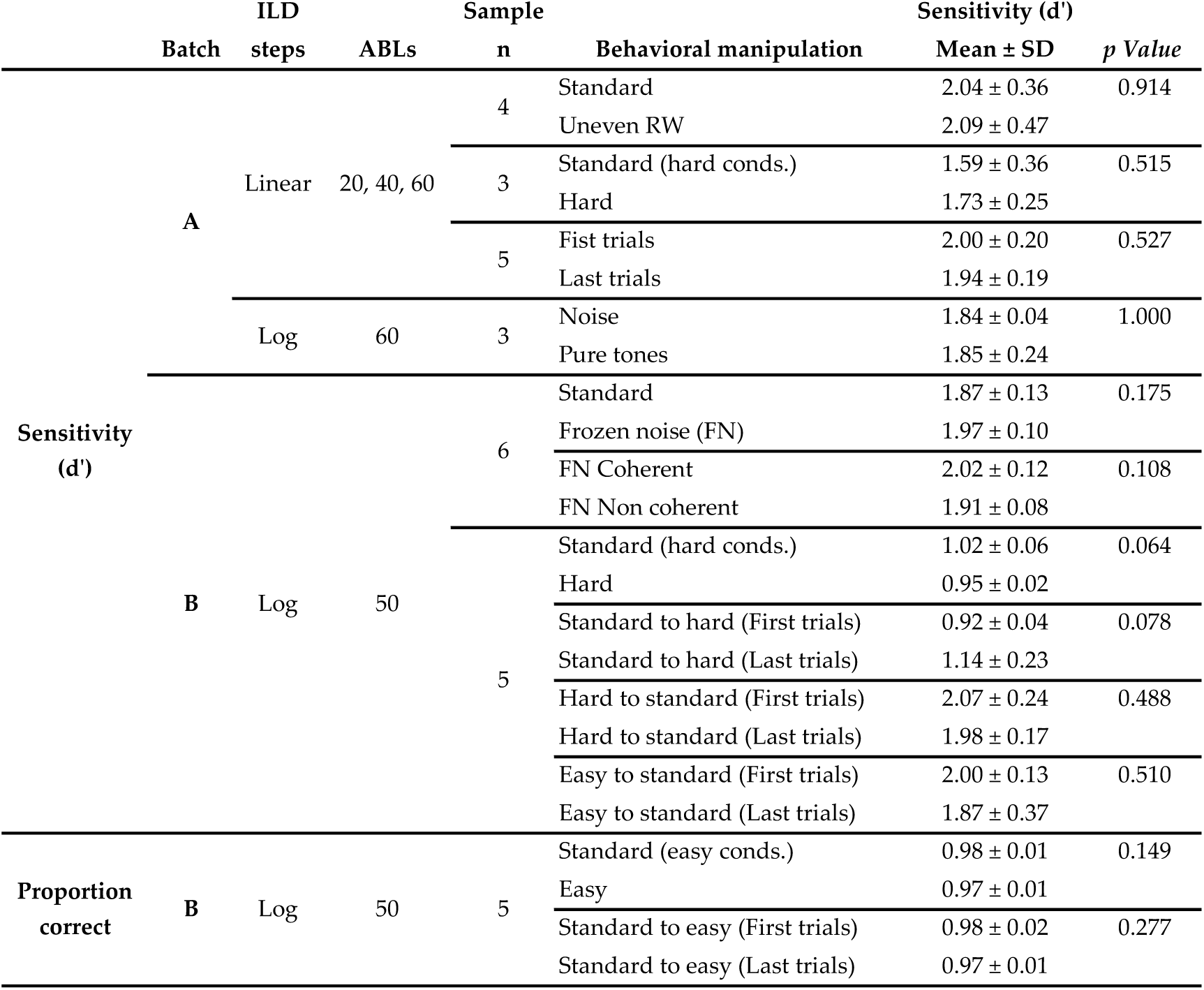
Results for the statistical comparison of sensitivity (*d*’) across all behavioral manipulations. See Methods for precise definition of these manipulations and for a description of the procedure used to obtain *p*-Values.

